# The splicing paralogues *SNRPB* and *SNRPN* control differential metabolic states

**DOI:** 10.64898/2026.02.11.705284

**Authors:** Feyza Polat-Haas, Ana Villalba, Polina Rusina, Anusha Gopalan, José Héctor Gibrán Fritz García, Azamat Akhmetkaliyev, Frank Rühle, Anna Einsiedel, Anna Szczepińska, Fridolin Kielisch, Jia-Xuan Chen, Susanne Nguyen, Thierry Schmidlin, Simon Hippenmeyer, M. Felicia Basilicata, Claudia Isabelle Keller Valsecchi

## Abstract

Gene duplication underlies evolutionary innovation, yet many paralogues remain highly similar, raising questions about their functional divergence and physiological relevance. The spliceosomal Sm core protein SNRPB and its mammalian-specific paralogue SNRPN share over 90% sequence identity, but their distinct expression patterns - SNRPB being ubiquitous and SNRPN confined to the brain - suggest specialized functions. Why mammals have two different spliceosomes has remained obscure. Here, we generated isogenic human cell lines expressing ectopically either SNRPB or SNRPN exclusively and found that SNRPN stabilizes transcripts involved in energy metabolism and mitochondrial function, leading to increased mitochondrial abundance and oxygen consumption. Despite similar spliceosomal interactomes, SNRPN more strongly associates with the PRMT5 methylosome complex and exhibits dynamic arginine methylation in its C-terminal region that is sensitive to translation inhibition and amino acid availability. The SNRPN-dependent transcriptome responds to translation inhibition by stabilizing long, intron-rich genes involved in amino acid and energy metabolism. Our findings reveal a nutrient-sensitive, methylation-dependent mechanism that differentiates the two paralogues. This suggests that SNRPN functions as a metabolic-specialized spliceosomal subunit thereby providing tissue-specific adaptation of RNA processing in mammals.

## Introduction

Gene duplication is a widespread phenomenon that facilitates functional diversification and adaptation, with numerous reported examples of new gene functions arising after duplication ^1–4^. However, there are also many cases where gene duplicates show little signs of divergence, indicating that a substantial fraction of highly similar gene duplicates can be redundant or at least partially redundant. Determining whether genes are fully interchangeable is challenging, as functional overlap can be context-dependent. The budding yeast transcription factor paralogues Msn2 and Msn4 are an illustrative case, as they were initially thought to regulate overlapping sets of stress-response genes. However, systematic analysis across conditions revealed that retaining both factors resolves regulatory conflicts that emerge from transcriptional noise when environments fluctuate ^5^. Besides their relevance in evolution and regulatory complexity, gene paralogues also receive high attention in a clinical context. This is exemplified by the antisense oligonucleotide Spinraza, which treats spinal muscular atrophy caused by loss of SMN1. Spinraza exploits the functional overlap of *SMN1* with the *SMN2* paralogue to restore functional SMN protein production ^6^.

The spliceosome is an essential multisubunit RNA-protein complex that catalyzes the removal of introns to create functional mRNA transcripts. SNRPB, along with other Sm proteins (SNRPD1, SNRPD2, SNRPD3, SNRPE, SNRPF, SNRPG), forms the core of the U1, U2, U4, and U5 small nuclear ribonucleoproteins (snRNPs). Because splicing is an essential process, the absence of *SNRPB* in species with only a single Sm orthologue causes lethality (e.g. fruit flies ^7^). Different from most other species, mammals possess a duplicated version of the *SNRPB* gene known as *SNRPN*, with the two paralogs sharing more than 90% sequence identity ^8^. While most tissues express *SNRPB*, in the brain it is replaced by *SNRPN* ^9–11^. To date, it is unknown whether these mutually exclusive subunits give rise to two distinct mammalian spliceosomes and what their respective functions are.

Cerebro-Costo-Mandibular syndrome (CCMS) is a congenital skeletal disorder caused by heterozygous mutation of *SNRPB*, and therefore represents a condition in which *SNRPN* does not functionally compensate for *SNRPB* loss ^12^. SNRPN, in turn, is encoded from the imprinted 15q11-q13 locus whose mutation causes Prader-Willi syndrome (PWS), a neuroendocrine and neurobehavioral disorder. PWS patients display mild intellectual disability along with metabolic features, including hyperphagia and obesity as well as reduced energy expenditure. Comparisons of patient mutation profiles indicate that PWS does not result from a single causative mutation, but rather from diverse genetic mechanisms that disrupt paternal gene expression at the 15q11-q13 locus. The majority of PWS patients (∼60%) exhibit larger deletions affecting multiple genes and comparison of the mutation profiles point towards a key role for the SNORD116 snoRNA cluster open reading frames in PWS pathogenesis ^13–15^. Curiously though, patients with microdeletions affecting only SNRPN exon 2 and 3 have been reported to exhibit symptoms despite retaining SNORD116 expression ^14–16^. In mice, knock-out (KO) of only *SNRPN*, while leaving other genes within the PWS critical region intact, was reported to have no obvious phenotypic defects. The KO line could be maintained in homozygosity indicating that SNRPB can replace SNRPN for organismal viability, at least under the standard conditions in a mouse facility scored by the authors ^13^. These findings are also in line with SNRPB protein expression detected in brain tissue of PWS patients, likely being responsible for sustained splicing in absence of SNRPN ^8^. Since the late 1990s, these findings have not been further investigated, for example regarding metabolic phenotypes or more subtle cognitive problems observed in PWS. In summary, the reason for the existence of two distinct spliceosomes in mammals containing either SNRPB or SNRPN and their functional distinction in RNA processing remains unresolved.

## Results

### *SNRPN* increases the abundance of transcripts involved in metabolic functions

SNRPB shows expression across a broad range of tissues, whereas SNRPN is strongly expressed in the brain (**Supp Fig. 1a**). Because the two paralogues are expressed in tissues, which differ in many trans-acting factors (e.g., transcription and splicing factors), it is challenging to disentangle intrinsic from extrinsic effects on their functional divergence. To address this, we generated isogenic HEK cell lines using the Flp-In™ T-REx™ 293 system, allowing doxycycline-inducible expression of either *SNRPB* or *SNRPN* from a single-copy cDNA transgene tagged with a C-terminal HA and mNeonGreen epitope (**Fig. 1a**). HEK cells endogenously express SNRPB, but not SNRPN, neither at the transcript nor the protein level (**Extended Data Fig. 1b**). Upon induction of the transgenes, ectopic SNRPB/N proteins could be expressed similar to endogenous levels (**Extended Data Fig. 1c**). Since *SNRPB* is a common essential gene in DepMap ^17^ we used CRISPR/Cas9 to delete the endogenous *SNRPB* gene only after transgene insertion, so the cells relied exclusively on the transgenic copy of either *SNRPB* or *SNRPN* (hereinafter referred to as *SNRPB-* or *SNRPN*-cells, respectively). PCR, Sanger Sequencing as well as immunoblotting confirmed successful deletion of endogenous SNRPB and expression of the transgenes (**Extended Data Fig. 1d-g**). In immunostainings, SNRPB and SNRPN transgenes localized to the nucleus and, as expected ^11^, formed foci that co-localized with Cajal bodies, as indicated by overlap with Coilin staining (**Fig. 1b**).

**Fig. 1.**
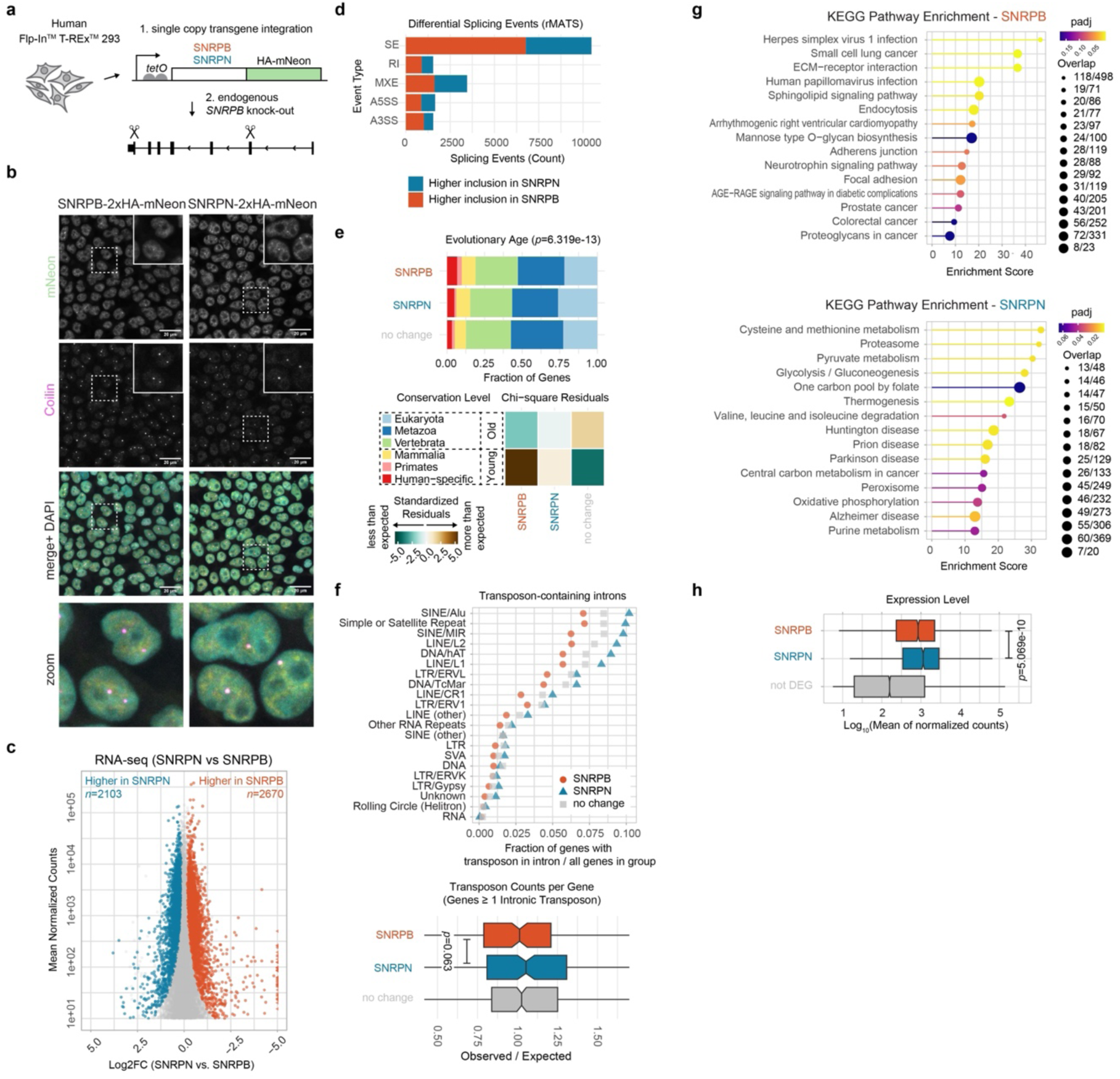
Transcriptome differences in *SNRPB* and *SNRPN*-expressing cells. (a) Scheme showing single-copy, doxycycline-inducible HA-mNeon-tagged *SNRPB* or *SNRPN* transgenes with the Flp-In^TM^ T-REx^TM^ 293 system, followed by CRISPR/Cas9-mediated knockout of endogenous *SNRPB*. (b) Representative immunostaining of Coilin (magenta) with mNeon fluorescence from tagged transgenes (yellow) in *SNRPB*- or *SNRPN*-expressing cells. DAPI in cyan, scale bar 20 μm. Areas marked with a dashed square are magnified in the “zoom” panel. The experiment was conducted two independent times and the images shown are maximum intensity projections of z-stacks. (c) MA-Plots from RNA-seq showing normalized read counts versus log2FC comparing *SNRPB*- versus *SNRPN*-cells. Differentially expressed (DE) genes (FDR≤0.01) are red (higher levels in SNRPB-cells) or blue (higher levels in *SNRPN*-cells). (d) Significant (FDR<0.05) splicing event counts identified by rMATS comparing *SNRPB*- versus *SNRPN*-cells for inclusion level difference and split by type: SE (skipped exon), RI (retained intron), MXE (mutually exclusive exon), A5SS (alternative 5’ splice site), and A3SS (alternative 3’ splice site). (e) Barplot showing the conservation level of DE genes as in (**c**). Genes were classified based on whether they have an orthologue in yeast (Class Eukaryota), fruit flies (Class Metazoa), zebrafish (Class Vertebrata), mouse (class Mammalia), or chimpanzee (Primates). If no orthologue was annotated, the genes were assigned as being specific to humans. As a control group we used non DE genes (FDR>0.01). For statistical testing, the genes conserved in Eukaryota, Metazoa, or Vertebrata were assigned as “Old” and genes conserved in only Mammalia, Primates or Human-specific as “Young”. The distribution of Old versus Young genes across groups was tested using a Pearson chi-square test of independence. The heatmap displays standardized Pearson residuals from the chi-square test for each group × age-bin, where positive residuals indicate enrichment (more genes than expected) and negative residuals indicate depletion (fewer genes than expected). (f) Analysis of transposon content in DE genes identified in (**c**). Transposon annotations from RepeatMasker were intersected with introns from canonical transcripts (defined as the longest transcript per gene) and genes containing ≥1 intronic transposon overlap were assigned to three groups: DE higher in *SNRPB*-cells, DE higher in *SNRPN*-cells or not DE. For each transposon class, the fraction of genes containing ≥1 intronic transposon of that class was calculated relative to the total number of genes in each group. To quantify transposon enrichment accounting for gene length, an expected number of intronic transposons per gene was calculated according to cumulative intron length. The observed/expected ratio was computed for each gene and visualized as boxplots (*p*-value from an unpaired Wilcoxon rank-sum test). (g) KEGG pathway enrichment analysis of DE genes from (**c**). Enrichment scores are plotted on the *x*-axis, with dot size corresponding to gene overlap and color representing adjusted *p*-values. (h) Expression levels (DESeq2 mean normalized read counts) of DE genes from (**c**) or non DE genes (FDR>0.01). *p*-value from an unpaired Wilcoxon rank-sum test.

We performed stranded mRNA sequencing on two independent *SNRPB*-cell clones and four independent *SNRPN*-cell clones, alongside parental HEK cells (Mapping statistics, replicate information and quality control available on ^18^). Clustering analysis revealed that *SNRPB*-expressing cells were similar to the parental HEK cells (**Extended Data Fig. 2a**). In contrast, *SNRPN*-expressing cells exhibited a markedly different transcriptomic profile and clustered separately (**Extended Data Fig. 2a**). Differential expression (DE) analysis using DESeq2 (FDR<0.01) identified 2,103 genes with significantly higher expression in *SNRPN*-cells and 2,670 genes with higher expression in *SNRPB*-cells (**Fig. 1c**) (**Supplementary Data 1**) (Complete results tables available on ^18^). Analysis of the splicing landscape using rMATS ^19^ uncovered several thousand splicing events, with the majority of differential events between *SNRPB* and *SNRPN*-cells being skipped exons (SE) (**Fig. 1d** and ^18^). Several hundred of these genes with differential splicing events were also overall DE at steady-state RNA levels (**Extended Data Fig. 2b**).

We investigated the gene features of (1) DE and non-DE gene groups and (2) differentially spliced transcripts. Neither group showed significant differences in intron numbers (**Extended Data Fig. 2c**). The cumulative intron length and the length of the longest intron were significantly greater in non-DE genes, but no differences were observed between *SNRPB* and *SNRPN* targets (**Extended Data Fig. 2d**). Analysis of 5′ and 3′ splice site strengths using MaxEntScan ^20^ revealed no differences between *SNRPB/N*-regulated and non-DE genes (**Extended Data Fig. 2e**). Due to the evolutionary context of gene paralogue formation, we also investigated the conservation level of the genes differentially stabilized by *SNRPB* and *SNRPN*. Evolutionary young genes (i.e. those with orthologues in mammals, primates or humans, but not other eukaryotes) were significantly overrepresented among genes stabilized by SNRPB (**Fig. 1e**). Over evolutionary time, transposons can accumulate within introns, potentially challenging the splicing machinery ^21^. Therefore, we examined transposon content across gene groups and found that *SNRPB*-stabilized genes tended to contain less transposons, whereas *SNRPN*-targets were enriched for them (**Fig. 1f**).

Lastly, we explored the functional characteristics of differentially regulated genes and conducted KEGG pathway analysis (**Fig. 1g**). *SNRPB*-stabilized genes were enriched in pathways associated with cellular surface functions, including “ECM−receptor interaction,” “Endocytosis,” and “Adherens junction.” Interestingly, SNRPN-*stabilized* genes had very different gene functions and were involved in metabolic processes, with enriched KEGG-pathways like “Cysteine and methionine metabolism,” “Pyruvate metabolism,” and “Glycolysis/Gluconeogenesis.” Examples for genes stabilized in *SNRPN*-cells are prominent metabolic enzymes such as *PGM1* (phosphoglucomutase 1), *PGK1* (phosphoglycerate kinase 1), *PK* (pyruvate kinase), *PGLS* (6-phosphogluconolactonase) or *ASNS* (asparagine synthetase) ^18^. In addition, 17 of the 42 large and 6 of the 35 small mito-ribosomal subunits encoded in the nucleus were more stable in *SNRPN*-cells. In line with these findings, genes stabilized by *SNRPN* had overall significantly higher expression levels, consistent with metabolic functions encoded by these genes (**Fig. 1h**). We conclude that *SNRPB* and *SNRPN* expression induces distinct transcriptomes, indicating that these duplicates fulfil different functions.

### Distinct metabolic states in SNRPB and SNRPN expressing cells

Intrigued by the transcriptome differences pointing towards distinct roles in metabolism, we next compared the *SNRPB/N*-cell metabolomes using Liquid Chromatography coupled to mass spectrometry (LC-MS). After data normalization (see methods) we performed principal component analysis, which showed that *SNRPB*- and *SNRPN*-cells are metabolically distinct (**Fig. 2a**). Accordingly, the samples clustered according to the cells’ genotype in a clustering heatmap, with 31 metabolites exhibiting significant differences (22 with higher abundance in *SNRPN-* and 9 in *SNRPB*-cells, respectively) (**Fig. 2b-d**). In *SNRPN*-cells, the enriched metabolites included glucose, amino acids, and intermediates of amino acid metabolism such as aminoadipate, phenylalanine, lysine, or arginine (**Fig. 2d**) (**Supplemental Data 1**). Additionally, NAD+ levels were significantly increased (FDR=6.99E-5), while NADP+ and Nicotinamide remained unchanged (**Fig. 2d**). KEGG pathway analysis revealed “Arginine and Proline Metabolism” (-log_10_(*p*)=3.3759) as well as “Arginine biosynthesis” (-log_10_(*p*)=2.7184) as the top differentially regulated pathways (**Supplemental Data 1**) (**Extended Data Fig. 3a**). Conversely, *SNRPB-* cells displayed significant elevation of metabolites related to nucleotide metabolism (AMP, GMP, UMP, Hypoxanthine) (**Fig. 2d**).

**Fig. 2.**
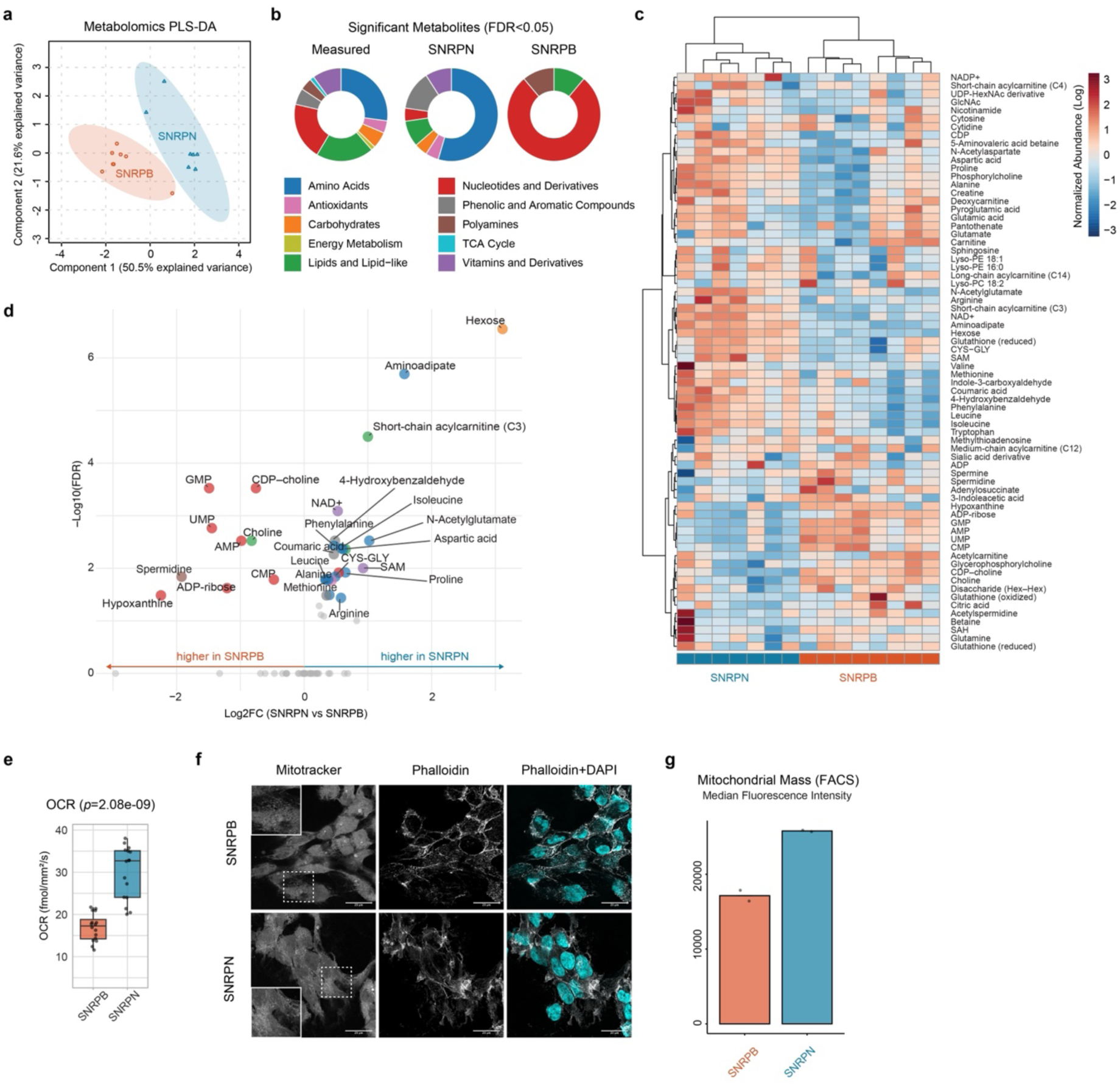
Metabolic differences in *SNRPB* and *SNRPN*-expressing cells. (a) Mass spectrometry-based metabolomics analysis comparing *SNRPB*- (red circles) with *SNRPN*- (blue triangles) expressing cells. In the Partial least squares discriminant analysis (PLS-DA), the separation between groups reflects differences in global metabolomic profiles, with each point representing a biological replicate, and colors indicating experimental groups. See also **Supplemental Data 1.** (b) Metabolomics as in (a); pie charts show metabolites with statistically significant changes (FDR<0.05) between *SNRPB*- and *SNRPN*-cells, in comparison with all measured metabolites/classes. (c) Metabolomics as in (**a**); clustering heatmap of all measured metabolite intensities (rows) across replicates (columns). (d) Metabolomics as in (**a**); dot plot of log2FC (*x*-axis) versus significance (*y*-axis) for all metabolites (non-significant metabolites, FDR>0.05, are shown in grey at -log10FDR=0). Selected metabolites with high fold change and significance are labeled; classes colored as in (**b**). (e) Box plot showing oxygen consumption rate in *SNRPB*- and *SNRPN*-cells. Each data point represents a replicate, calculated as the mean of the recorded values between 16 and 24 hours after the start of the experiment. Three independent experiments were conducted, one representative experiment is shown here, the other two experiments are in **Extended Data** Fig. 3b. Statistical significance was assessed using a linear mixed-effects model across all three experiments. (f) Confocal microscopy of *SNRPB*- and *SNRPN*-cells showing mitotracker and phalloidin staining, along with DAPI (cyan in merged panel), the scale bar is 20 μm. The pictures reflect maximum intensity projections of z-stacks. (g) Flow cytometry analysis of *SNRPB*- and *SNRPN*-cells for mitochondrial mass using mitotracker staining. Overlaid datapoints reflect n=2 biological replicates, with the barplot indicating the median fluorescence intensity of 100,000 measured cells. Gating and raw data in **Supplementary Data 2**.

We next asked whether the observed changes in metabolites (higher glucose, free amino acids, and NAD+ in *SNRPN*-cells) were also reflected by metabolic differences at the cellular level. Because these metabolites are central to cellular energy and redox metabolism and feed into mitochondrial oxidative pathways, we assessed mitochondrial function by measuring the oxygen consumption rate (OCR). *SNRPN*-cells exhibited a significantly increased OCR (**Fig. 2e**; **Extended Data Fig. 3b**; control OCR measurement upon Oligomycin treatment, which blocks mitochondrial ATP-Synthase in **Extended Data Fig. 3c**). The metabolic differences were also reflected in subtle differences observed in confocal microscopy of cells stained with MitoTracker (**Fig. 2f**), which labels mitochondria and provides a readout of mitochondrial abundance within cells. We also noticed changes in cytoskeletal organization as revealed by Phalloidin staining (**Fig. 2f**). Quantification of mitochondrial mass using flow cytometry showed that *SNRPN*-cells indeed contain more mitochondria (**Fig. 2g**) (Gating in **Supplemental Data 2**). Overall, these findings are consistent with a metabolic “reprogramming” induced by *SNRPN* expression.

### Stronger methylosome association and arginine methylation patterns distinguish SNRPN from SNRPB

The finding that SNRPN expression can give rise to changed transcriptomes and metabolic states is particularly striking given that SNRPB/N encode nearly identical proteins, differing only by a few amino acids located in the poorly characterized, intrinsically disordered proline-rich region at their C-termini (**Extended Data Fig. 4a-b**). These few distinctive amino acids are conserved in each paralogue throughout evolution, indicating functional relevance (**Extended Data Fig. 4a-b**). Using immunoprecipitation via the C-terminal HA-tag followed by LC-MS, we next profiled the interaction partners of SNRPB (**Fig. 3a**) and SNRPN (**Fig. 3b**) comparing them to an untagged control cell line (**Extended Data Fig. 4c**). As expected, both paralogues robustly interacted with the Sm/SMN complex and key snRNP components, which were among the most highly enriched interactors in both datasets (**Fig. 2a-b, Extended Data Fig. 4d**). More generally, the two interactomes showed substantial overlap in interactor identity with both high-confidence (padj<0.001, log2FC >2) and low-confidence (padj<0.05, log2FC>4) thresholds (**Fig. 3c, Extended Data Fig. 4e**), Moreover, for the majority of interactors, the magnitude of enrichment was highly similar between the two paralogues, indicating comparable interaction strengths across both interactomes (**Fig. 3d**, Pearson R^2=0.79). However, a few notable exceptions were observed - some that were uniquely associated with SNRPB or SNRPN, and others that showed a substantially stronger enrichment in one paralogue compared to the other (**Fig. 3d**). In particular, SNRPN showed stronger association with multiple components of the methylosome complex, including the arginine methyltransferase PRMT5 and its substrate adaptors CLNS1A (also known as pICln), RIOK1, and WDR77, which were approximately 10-fold more enriched in SNRPN compared to SNRPB pulldowns.

**Fig. 3.**
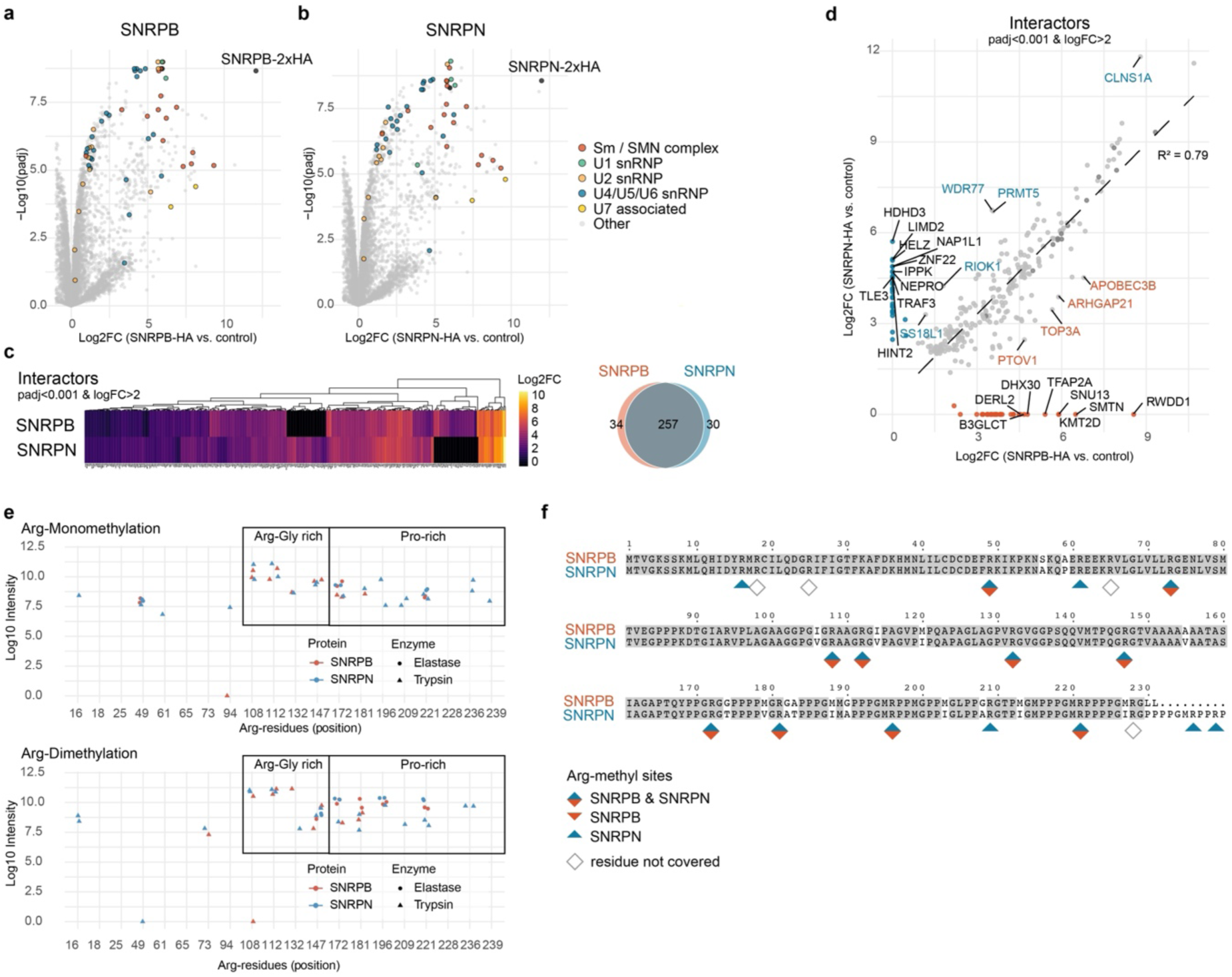
Interactors and PTMs of SNRPB and SNRPN. (a) Volcano plots of HA immunoprecipitations (IP)-mass spectrometry experiment comparing HA-mNeon tagged SNRPB to untagged control cells. Interactors belonging to snRNP complexes are shown in color (see **Extended Data** Fig. 4c for Immunblot of the IP and **Extended Data** Fig. 4d for individual complex subunits and classification). *n*=4 replicates for each SNPRB/N were conducted. Adjusted *p*-values and log2FC were obtained with limma ^66^. (b) as in (**a**) but for SNRPN interactors. (c) as in (**a**) Hierarchical clustering heatmap of interactors (padj<0.001, log2FC>2 in either SNRPB, SNPRN or both) with log2FC of enrichment shown in color. Right: Venn diagram showing the overlap of the identified interactors. (d) as in (**a**) Scatter plot comparing enrichment of interactors (padj < 0.001, log2FC > 2) in SNRPB, SNRPN, or both. The dashed diagonal indicates equal enrichment between the two interactomes, and the Pearson correlation (R²) is shown. Selected proteins displaying strong paralog-specific enrichment are labeled. (e) Quantification of PTMs by mass spectrometry (immunoblot of the IP shown in **Extended Data** Fig. 5h). Two independent high-stringency HA immunoprecipitations (IPs) were performed for each SNRPB and SNRPN. Each replicate IP sample was split and digested with either elastase (circles) or trypsin (triangles) to generate peptides. Each data point represents Arg-mono or dimethylation intensity in single replicate measurement; if a residue was not covered in a specific digest or replicate no symbol is shown. PTM intensities were calculated by summing peptide-level intensities at each modification site and normalized to the total peptide intensity per sample. Raw data and quantification of additional PTMs are provided in **Supplemental Data 1**. (f) PTM analysis as in (**e**) on the linear protein sequences of SNRPB and SNRPN. Colored diamonds indicate methylation detected in both paralogues (blue-orange), exclusively in SNRPB (orange), or exclusively in SNRPN (blue). Open diamonds mark arginine residues not covered by mass spectrometry.

The methylosome complex participates in the regulation of snRNP biogenesis through arginine methylation, where this posttranslation modification (PTM) controls interactions with other splicing factors (e.g. the SMN protein) ^22,23^. In addition, arginine methylation of SNRPB is important for the release of RNAs from chromatin ensuring a productive splicing process ^24,25^. Sm proteins are among the most prominent targets of multiple arginine methyltransferases ^26^, but a differential role with regards to the SNRPB/N paralogues and/or different tissues has not been explored yet.

To understand whether increased methylosome association of SNRPB/N results in differences in PTM levels, we designed a high-stringency immunoprecipitation protocol specifically optimized for the detection of differential PTMs, followed by quantitative MS (Results Table for arginine and lysine methylation, phosphorylation and proline hydroxylation in **Supplemental Data 1**). To achieve high sequence coverage, we employed two enzymatic digestion strategies (Trypsin and Elastase; see Methods). In both SNRPB/N, we found multiple arginines modified by mono- and dimethylation (**Fig. 3e**-**f**). The Arg-Gly-rich region (positions 90-170) showed overall similar methylation status between the paralogues, but differences emerged in the Sm-domain and Pro-rich region (position 170 onwards) where several methylation sites were detected exclusively in SNRPN. In addition, extra methylation sites were detected for the Arg-residues in the C-terminal extension exclusive to SNPRN (Arg236, Arg239).

Taken together, SNRPN displays stronger association with the methylosome complex and harbors additional, paralogue-specific arginine methylation sites that are absent in SNRPB

### Translation inhibition and amino acid restriction modulates SNRPN through arginine methylation

Protein arginine methylation is tightly linked to methyl donor availability and thus, one-carbon metabolism. It is also particularly abundant in the brain ^27^ - the tissue where SNRPN is expressed. We reasoned that cellular pathways responding to environmental or metabolic cues (e.g. nutrient availability), could dynamically regulate SNRPN’s arginine methylation state. Such a mechanism could provide a potential link between the metabolic differences observed in *SNRPB*- and *SNRPN*-cells (**Figs. 1 and 2**) and the distinctive PTM patterns in their C-termini (**Fig. 3**). Inspired by the fact that well known Sm-antibodies like anti-Smith Antigen Antibody Y12 are PTM-sensitive for recognition ^28,29^, we screened multiple commercially available antibodies for detection of the C-terminal fragment of SNRPB/N. We identified the monoclonal antibody 12F5, which successfully recognized the C-terminal fragments 190-231 (SNRPB) and 190-240 (SNRPN), but failed to detect a truncated SNRPB (1-190) protein in immunoblots (**Fig. 4a**-**b**) (12F5 labelled as “SmB/B’/N”). Following treatment with two well-established arginine methyltransferase inhibitors ^26,30^, SNRPN, but to a substantially lesser extent, SNRPB detection, decreased with 12F5 (**Fig. 4c**). Immunoblotting for the HA epitope verified consistent overall protein expression levels across all conditions. (**Fig. 4a-c**). This indicates that 12F5 detects an epitope in the Pro-rich region, which is sensitive to arginine methylation.

**Fig. 4.**
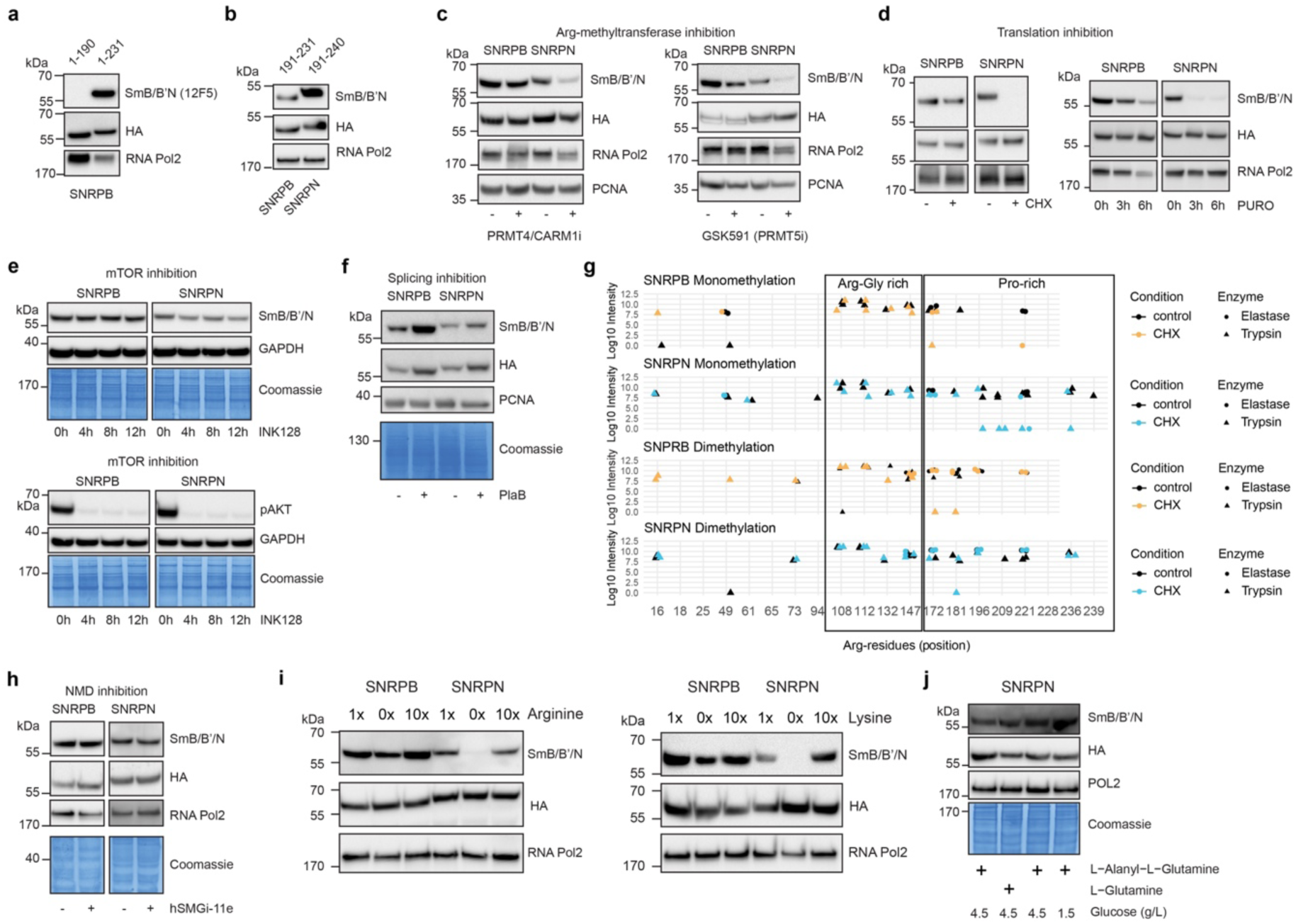
PTM alterations of SNRPB/N in different conditions. (a, b, c, d, e, f, h, i, j) Cropped immunoblots of *SNRPB*- and *SNRPN*-cells under various conditions. RNA Pol2, PCNA, GAPDH and Coomassie serve as loading controls. HA-antibody shows overall protein stability differences. Monoclonal SmB/B’/N antibody (clone 12F5) shows detection differences for SNRPN under certain conditions. All experiments were performed at least twice with similar results. For control conditions, the solvent (DMSO, Ethanol, Water) in which the inhibitors are dissolved were added to the media. (a) Truncation mutant of SNRPB - amino acids 1-190 versus 1-231 (full-length) (b) SNRPB/N mutants encoding only amino acids 190 until C-terminal end. (c) *SNRPB/N*-cells treated 27 hours with 5 μM of PRMT4/CARM1 (left) or 6 days with 10 μM of PRMT5 inhibitor (right). (d) *SNRPB/N*-cells treated with translation inhibitors Cycloheximide (CHX, 4 hours, 100 μg/mL) or Puromycin (PURO, 20 μM) (e) *SNRPB/N*-cells treated with mTOR inhibitor INK128 (320 μM). The pAKT blot serves as a positive control for effective treatment. (f) *SNRPB/N*-cells treated with splicing inhibitor Pladienolide B (PlaB, 16 hours, 100 nM). Effective treatment was evaluated by RT-qPCR, see **Extended Data** Fig. 5c. (g) Quantification of post-translational modifications (PTMs) by mass spectrometry, as in Fig. 3e, including CHX-treated conditions. The experiment was performed together, so the same datapoints from Fig. 3e are also shown here (control condition). Each replicate IP sample was split and digested with either elastase (circles) or trypsin (triangles) to generate peptides for mass spectrometry. (h) *SNRPB/N*-cells treated with NMD inhibitor hSMGi-11e (24 hours, 0.6 μM). Effective treatment was evaluated by RT-qPCR, see **Extended Data** Fig. 5i. (i) *SNRPB/N*-cells grown in regular media (1x), media containing no arginine (left, 0x) or no Lysine (right, 0x) or 10-fold higher arginine (left, 10x) or Lysine (right, 10x) concentrations. Treatment was conducted for 16 hours. (j) *SNRPB/N*-cells grown in regular media (lane 1), media containing L-Glutamine instead of L-Alanyl-L-Glutamine (lane 2), regular media (lane 3) or regular media with reduced glucose content (lane 4). Treatment was conducted for 16 hours.

Due to this property of 12F5, we used it as an experimental tool to screen for differential PTM regulation of SNRPB/N. We observed a strong reduction in SNRPN detection by 12F5 following treatment with Cycloheximide (CHX) and Puromycin (both translation inhibitors), Actinomycin D (transcription inhibitor) (**Fig. 4d**, **Extended Data Fig. 5a-b**), as well as a moderate reduction after heat shock and INK128 treatment (inhibitor of the mTOR kinase ^31^) (**Fig. 4e**, **Extended Data Fig. 5a**). Treatments with Camptothecin (DNA Topoisomerase I inhibitor), STM2457 (m6A-RNA methylation inhibitor), Bortezomib (proteasome inhibitor), and Deferoxamine (iron chelator) did not result in any detection differences (**Extended Data Fig. 5a**), supporting that the observed effects are specific and not caused by general cellular stress or perturbations in transcriptome or proteome homeostasis. No signal decrease was observed after splicing inhibition with pladienolide B (PlaB) (**Fig. 4f**; RT-qPCR confirming effective inhibition by PlaB in **Supp. Fig 5c**). Instead, PlaB stabilized overall SNRPB and SNRPN proteins similarly, indicated by concordant increases in HA and 12F5 signals (**Fig. 4f**). The effects of CHX were not suppressed by PlaB, indicating that productive splicing is not required for PTM changes elicited by translational inhibition (**Extended Data Fig. 5d**).

**Fig. 5.**
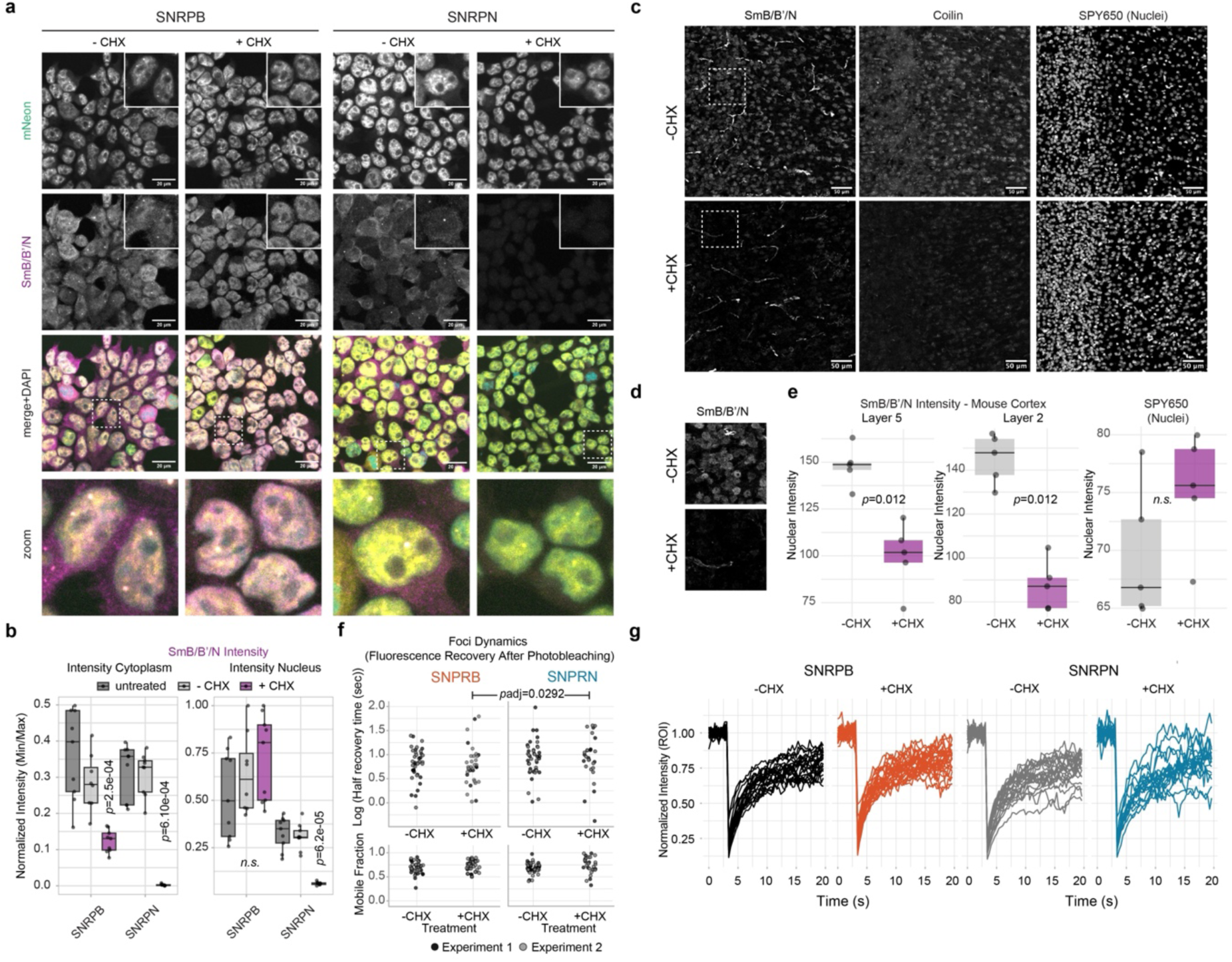
Cellular behaviour of SNRPB/N upon translational inhibition. (a) Representative confocal microscopy images of mNeon-tagged *SNRPB/N*-cells (yellow in merged panel), 12F5-immunostained SNRPB/N (magenta in merged panel) and DAPI (cyan in merged panel) with Ethanol (-CHX; Control) or CHX treatment (4 hours). The areas marked with a dashed square are magnified in the “zoom” panel. Pictures reflect maximum intensity projections of a z-stack. Scale bar 20 μm. The experiment was conducted two independent times. (b) Quantification of (**a**) for 12F5 immunostaining (SmB/B′/N) channel intensity. Boxplots show normalized cytoplasmic (left) and nuclear (right) intensities (see **Methods**), with each dot representing a field of view containing multiple quantified cells. Total cells quantified (*n*=): HEK - untreated 578, EtOH (−CHX) 541, CHX (+CHX) 359; SNRPB - untreated 580, EtOH 737, CHX 579; SNRPN - untreated 918, EtOH 876, CHX 692. Adjusted *p*-values were obtained using a Wilcoxon rank-sum exact test with Benjamini-Hochberg correction comparing fields of view. Comparisons between EtOH (-CHX; control) and untreated were not significant (all *p*adj>0.05). (c) Representative confocal microscopy images of mouse cortex stained with 12F5 (SmB/B′/N) and Coilin antibodies. Cycloheximide (CHX) or Ethanol Control (-CHX) was administered to P7 mouse pups by bilateral intracerebroventricular stereotaxic injection and allowed to act for 5 hours prior to tissue collection. Pictures reflect maximum intensity projections of a z-stack. Scale bar 50 μm. The staining was conducted three independent times. (d) Magnified area delineated with a dashed square in (**c**) (e) Quantification of mouse cortex 12F5 immunostaining (SmB/B′/N) or SPY650 dye intensity (z-projection). The intensity sum in the nuclear area was quantified and mean values per field of view calculated. In the box plot, each dot represents the mean of all nuclei within a single field of view. *p*-values comparing field-of-view means were determined using an unpaired Wilcoxon rank-sum test. Total number of quantified cells: Layer 2, *n*=455 (-CHX), *n*=417 (+CHX); Layer 5, *n*=229 (-CHX), *n*=208 (+CHX). (f) Dot plots showing the dynamics of nuclear foci in *SNRPB*- or *SNRPN*-cells, respectively, as measured by fluorescence recovery after photobleaching (FRAP). Each dot represents the recovery parameters (top: recovery time; bottom: mobile fraction) fitted to a given FRAP curve recorded after bleaching a given mNeon focus in the nucleus (normalized intensity curves in (**g**); details on normalization and data fitting in methods). Statistical significance was evaluated with a nonlinear mixed effects model. The experiment was conducted two times independently. (g) FRAP as in (**f**) Normalized intensity curves, where each line represents the recorded data of one bleached focus in the nucleus. See also **Extended Data** Fig. 7d and **Extended Data** Fig. 7e (intensity curves of second experiment).

To ensure that these results are not confounded by overall protein stability differences we employed (1) an independent protein extraction method where cells were directly lysed in trichloracetic acid (**Extended Data Fig. 5e**), (2) another polyclonal antibody (**Extended Data Fig. 5f** and methods), and (3) SNRPB/N protein quantification by flow cytometry measuring mNeon-tag fluorescence (**Extended Data Fig. 5g**). We also used an alternative construct where the 2xHA-mNeon tag is positioned at the N-terminus (**Extended Data Fig. 5b**). All approaches substantiated the aforementioned results - CHX sensitivity was observed for *SNRPN*- but not *SNRPB*-transgenes. To confirm arginine methylation changes for SNRPN in the context of translational inhibition, we again performed immunoprecipitation using the C-terminal HA-tag followed by PTM analysis by MS (**Extended Data Fig. 5i**). Methylation in the Arg-Gly–rich region was unaffected by CHX treatment, but for several arginine sites in the Pro-rich region of SNRPN methylation levels became undetectable (**Fig. 4g**). Other PTMs like lysine mono- and dimethylation or proline hydroxylation were unchanged (**Supplemental Data 1**). This MS data, together with the methyltransferase enzyme inhibitor (**Fig. 4c**), supports that arginine methylation is part of the axis leading to the distinct behaviour of SNRPB/N.

mRNA translation is a requirement for nonsense mediated RNA decay (NMD), a mechanism that recognizes and degrades transcripts with premature stop codons, such as by-products of splicing errors. However, different from translation inhibition with CHX, SNPRB/N remained unaffected after NMD inhibition with the SMG1 inhibitor 11e ^32^ (**Fig. 4h**; RT-qPCR confirming inhibition in **Extended Data Fig. 5j**). This indicates that the alterations triggered by translation inhibition are not linked to NMD pathway perturbation, but - taking into account the observed changes upon mTOR inhibition with INK128 - rather connect to metabolism.

The mTOR kinase complex is a central regulator of anabolic and catabolic processes for organismal growth. It integrates environmental inputs - primarily amino acid and growth factor availability - and, in response, exerts its regulatory effects through the stimulation of protein translation, while concomitantly suppressing autophagy and lysosome-mediated lipid degradation ^33^. We thus investigated the impact of modulating nutrients (arginine, lysine, glucose and L-glutamine) to SNRPN’s PTM status. Intriguingly, arginine and lysine depletion, but not providing them in excess, phenocopied the effect of CHX and also induced a loss of the 12F5 band for SNRPN, while SNRPB was unaffected (**Fig. 4i**). Lowering glucose or changing the Glutamine source (L-Glutamine versus L-Alanyl-L-Glutamine, also known as Glutamax) had no effect on either paralogue (**Fig. 4j**).

Taken together, these results show that translational inhibition and amino acid restriction lead to a loss of arginine methylation in SNRPN’s C-terminus, indicating that this PTM unique to SNRPN is dynamically regulated by cellular conditions.

### Cycloheximide-dependent changes in SNRPB/N localization and dynamics

To better understand the cellular context of dynamic PTMs in SNRPN’s Pro-rich region, we focused on exploring the following non–mutually exclusive ideas: (1) change in subcellular localization ^23,34^, (2) PTM-dependent interaction partners ^35^ or (3) altered dynamics in Cajal bodies (also see **Fig. 1b**) ^36–38^. Using immunofluorescence microscopy, we found that the SNRPB/N-mNeon intensity was predominantly nuclear (indicated by a nuclear-to-cytoplasm ratio of around 9 for both paralogues) and remained unchanged in the nucleus upon CHX treatment (**Fig. 5a, Extended Data Fig. 6a-c**). After CHX treatment, both Coilin as well as SNRPB/N-mNeon foci numbers decreased significantly, but remained colocalized (**Extended Data Fig. 6d-f**). In the 12F5 staining, we detected a signal similar to mNeonGreen (homogenous nuclear with foci overlapping with Coilin), but in addition, notable cytoplasmic signal. As expected, differences appeared after CHX treatment: the nuclear 12F5 intensity remained unaffected in *SNRPB*-cells, but was fully lost in *SNRPN*-cells, while the cytoplasmic signal disappeared in both (**Fig. 5b**).

We next wanted to assess whether these changes also occur *in vivo* and examined SNRPN in the mouse cortex, where SNRPN begins to become strongly expressed at postnatal day 7 (P7) ^39^. CHX was delivered by stereotaxic injection and allowed to act for 5 hours before tissue collection and immunostaining. In control conditions, 12F5 staining was readily detected in the cortex and hippocampus, consistent with a high density of neurons (visualized by NeuN staining) (**Fig. 5c, Extended Data Fig. 6g**). Interestingly, the signal was non-uniform across different areas of the brain: only the neocortex (layers 2-5) and hippocampus (stratum pyramidale) exhibited the characteristic localization with foci in Cajal bodies (**Extended Data Fig. 6g-h**). The same was observed for Coilin staining, which also showed a nonuniform distribution, with discrete foci restricted to select areas. Quantification of SNRPN intensity (12F5 signal) revealed a marked decrease in CHX-treated cortex (**Fig. 5d-e**). This shows that CHX treatment changes endogenous SNRPN and that this pattern is observed not only in HEK293T cells but also *in vivo*.

We next analyzed SNRPB/N interaction partners after CHX treatment by performing IP-MS analysis in the transgenic HEK cells (**Supplemental Data 1**). The interaction with the major spliceosome components, methylosome complex, putative Arg-demethylases as well as major Tudor-domain proteins (Arg-methyl readers) remained largely unaffected for both paralogues (**Extended Data Fig. 7a-b**). In line with what has been reported by ^25^, this supports that Sm-RNP assembly and nuclear translocation is unchanged in CHX and thus, does not *a priori* require arginine methylation. A previously reported interactor of the SNRPB C-terminus, RBM5 ^40^, showed only weak and non-significant enrichment in our pulldown experiments under all conditions (based on log2FC, 885 proteins (SNRPB) or 1144 protein (SNRPN) were more strongly enriched in the respective pulldowns).

Focusing on valid SNRPB/N interactors that are localizing in the nucleus, the majority showed concordant changes after CHX in both SNRPB and SNRPN (diagonal / yellow dots in (**Extended Data Fig. 7c**), but some exceptions appeared that could potentially link to SNRPN’s ability to “sense” nutrients. For example, SNRPN in CHX gained interaction with PHIP/BRWD2, which functions in insulin and insulin-like growth factor signalling pathways ^41^. Besides binding the insulin receptor substrate-1, PHIP also contains a cryptic tudor domain and two bromodomains that function in transcription regulation and PTM recognition ^42–44^. Given these features, PHIP/BRWD2 could thus be a co-factor that connects changes in nutrient signals to arginine methylation dynamics in SNRPN.

Since the major spliceosome interaction partners did not change, we speculated that PTM changes elicited by CHX could impact splicing by alterations of the intrinsic behaviour of the SNRPN protein - for example through cation-phi interactions and altered dynamics in subnuclear bodies ^34,45^. To investigate this, we profiled the *in vivo* dynamics of nuclear SNRPB/N foci (Cajal bodies, **Fig. 1b**) using fluorescence recovery after photobleaching (FRAP). In control conditions, the SNRPB/N dynamics in Cajal foci were similar (**Fig. 5f**, **Extended Data Fig. 7d**). After CHX treatment we detected a significant difference, with SNRPN association with foci becoming slower compared to SNRPB (mean half recovery time SNRPB 1.83 sec, SNRPN 2.22 sec; *padj*=0.0292), while there were no differences in the mobile fraction (around 70% in all conditions) (**Fig. 5f-g**, **Extended Data Fig. 7d-e**).

In summary, CHX-induced changes distinguish SNRPN from SNRPB in the nucleus, occur *in vivo* and are accompanied by distinct interaction partners and Cajal-body dynamics, supporting a role for SNRPN as a nutrient-responsive regulator of splicing.

### SNRPN enables transcriptional and metabolic rewiring upon nutrient restriction

To determine whether these altered nuclear dynamics are accompanied by downstream changes in gene expression, we next compared the global transcriptional response to translation inhibition. We performed RNA-seq after 4-hour CHX treatment in *SNRPB-* and *SNRPN*-cells. Principal component analysis (PCA) revealed distinct profiles between the two cell types that were further amplified after treatment (**Extended Data Fig. 8a**). As expected, CHX induced widespread gene expression changes: 3,348 genes were upregulated and 2,809 downregulated in SNRPB-cells, while 4,887 were upregulated and 3,982 downregulated in *SNRPN*-cells (**Fig. 6a**). To identify cell-type–specific CHX responses, we compared differentially expressed genes across conditions. (**Fig. 6b**). A substantial number of genes overlapped between the two cell types (3,131 upregulated and 2,510 downregulated), but we also found unique responses: 1,752 genes were uniquely upregulated and 1,471 downregulated in *SNRPN*-cells, compared to only 216 uniquely upregulated and 299 downregulated in *SNRPB*-cells (**Fig. 6b**).

**Fig. 6.**
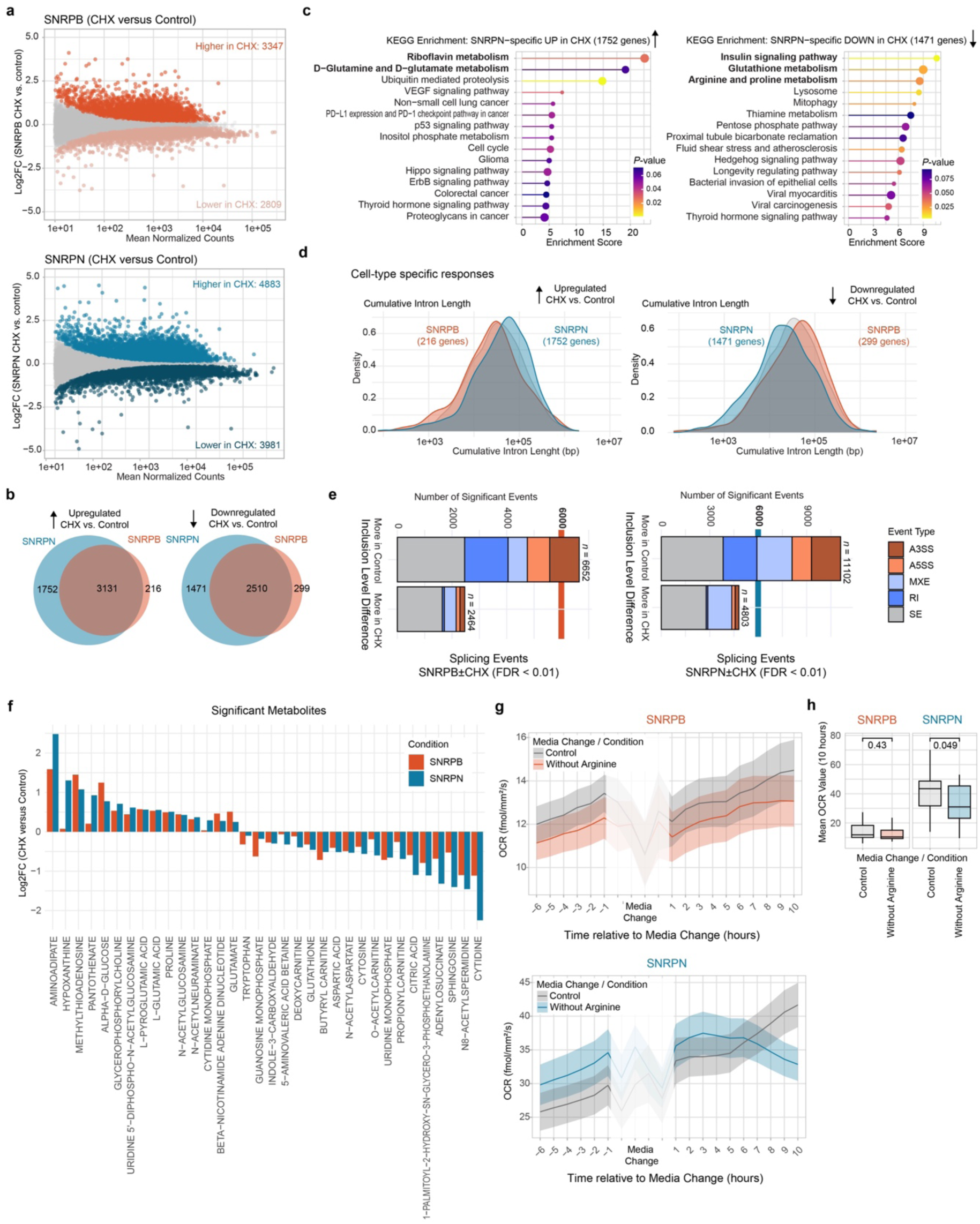
Transcriptome alterations after CHX treatment. (a) MA-Plots from RNA-seq showing normalized read counts versus log2FC comparing *SNRPB*- (top) and *SNRPN-*(bottom) cells in CHX versus Control (Ethanol) condition. DE genes (FDR≤0.01) are colored in red or blue, respectively. Grey = no significant change. (b) Venn diagram showing overlap of DE genes in CHX between *SNRPB*- and *SNRPN*-cells. (c) KEGG pathway enrichment analysis of uniquely DE genes in *SNRPN*-cells, see (**b**). Enrichment scores are plotted on the *x*-axis, with dot size corresponding to gene overlap and color representing adjusted *p*-values. (d) Density plot showing the cumulative intron lengths of DE genes that are cell-type specific, i.e. occur only in *SNRPB-* (red) or *SNRPN-* (blue) cells. Up- (left) or downregulated (right) genes after CHX treatment were analyzed separately. The control group is a list of gene subsampled from non-DE genes with comparable expression levels to *SNRPB* and *SNRPN* DE genes. (e) Significant (FDR<0.01) splicing event counts identified by rMATS comparing CHX versus Control condition in *SNRPB*- or *SNRPN*-cells, respectively. The events were split by type: SE (skipped exon), RI (retained intron), MXE (mutually exclusive exon), A5SS (alternative 5’ splice site), and A3SS (alternative 3’ splice site). The vertical line is shown to visually depict the absolute number of differential splicing events in *SNRPB/N*-cells. (f) Metabolomics as in Fig. 2; Control and CHX-treated samples were collected and analyzed together. Bar charts show log2FC (CHX versus Control) of statistically significant changing metabolites (FDR<0.05) in *SNRPB-* (red) and *SNRPN-* (blue) cells. (g) Ribbon plot showing OCR before and after arginine deprivation. Five independent experiments were conducted, with two or three replicate wells recorded in each experiment. In the ribbon plot, the *x*-axis indicates time (in hourly bins relative to the treatment) and the *y*-axis shows the mean OCR for each hour. Shaded areas correspond to SEM, and lines represent mean values across all experiments and replicates. Before treatment, cells were grown in full media; during treatment, the media was replaced with regular medium (Control) or medium lacking arginine. (h) Box plot showing mean oxygen consumption rates as in (**g**) in *SNRPB*- and *SNRPN*-cells at 10 hours after the medium switch. *P*-values were determined using an unpaired Wilcoxon rank-sum test.

KEGG analysis showed that *SNRPN*-specific CHX-responsive genes were enriched in pathways including glutamine metabolism, insulin signaling pathway and arginine/proline metabolism (**Fig. 6c**). In *SNRPB*-cells, some enrichment was observed e.g. in selenocompound and sulfur metabolism, but did not reach statistical significance (**Extended Data Fig. 8b**). The uniquely CHX-responsive genes in *SNRPN*-cells differed from non-DE and the ones found in *SNRPB*-cells: The 1,752 uniquely upregulated genes were generally longer (**Extended Data Fig. 8c**), with greater cumulative intron length (**Fig. 6d, left**) and longer lengths of the longest introns (**Extended Data Fig. 8d**). In contrast, uniquely downregulated genes were shorter, with fewer and shorter introns (**Fig. 6d, right**; **Extended Data Fig. 8c-e).** Given the central role of RNA processing in determining mature mRNA abundance, these findings suggest that, under translation-inhibiting conditions, SNRPN more efficiently processes transcripts containing long introns compared to SNRPB.

We next used rMATS (FDR<0.01) to quantify splicing changes between CHX-treated and control samples within each cell type (results tables available on ^18^). Consistent with the changes in overall expression levels, *SNRPN*-cells exhibited nearly twice as many CHX-induced splicing changes (11,102 events) as SNRPB-cells (6,652 events) (**Fig. 6e)**. Most alternative splicing types decreased substantially after CHX, but exon skipping (SE) occurred at comparable frequency in treated and control samples (**Fig. 6e)**. These patterns indicate that CHX leads to a broad reduction in alternative splicing, favouring constitutive isoform usage and reducing isoform diversity.

These transcriptome signatures (greater CHX-induced changes in *SNRPN*-cells) were also mirrored at the metabolite level: In *SNRPN*-cells, most metabolites with significant CHX-induced changes (e.g., aminoadipate, cytidine) showed greater fold changes than in *SNRPB*-cells (**Fig. 6f**). This was also reflected in altered cellular metabolic outputs, assessed by measurements of the oxygen consumption rate (OCR). Because CHX elicits strong effects common to both cell lines (**Fig. 6a**), these assays were performed under arginine-restriction rather than CHX treatment / translation inhibition. Comparing *SNRPB-* and *SNRPN*-cells before and after removal of the amino acid arginine from the growth medium, we found that arginine deprivation led to a marked reduction in OCR specifically in SNRPN-cells within a few hours, whereas *SNRPB*-cells remained largely unresponsive (**Fig. 6g-h**). Overall, these findings show that *SNRPN*-cells can more flexibly adapt their transcriptional and cellular programs in response to nutrient status and translation inhibition compared to *SNRPB*-cells.

### Mutation of SNRPN arginine residues reverts cells towards the *SNRPB*-program

Lastly, we were interested in linking the arginine methylation status to the observed transcriptional and cellular changes and generated a *SNRPN-R6K* mutant, in which arginines in the proline-rich region were replaced by lysines (**Fig. 7a**).

**Fig. 7.**
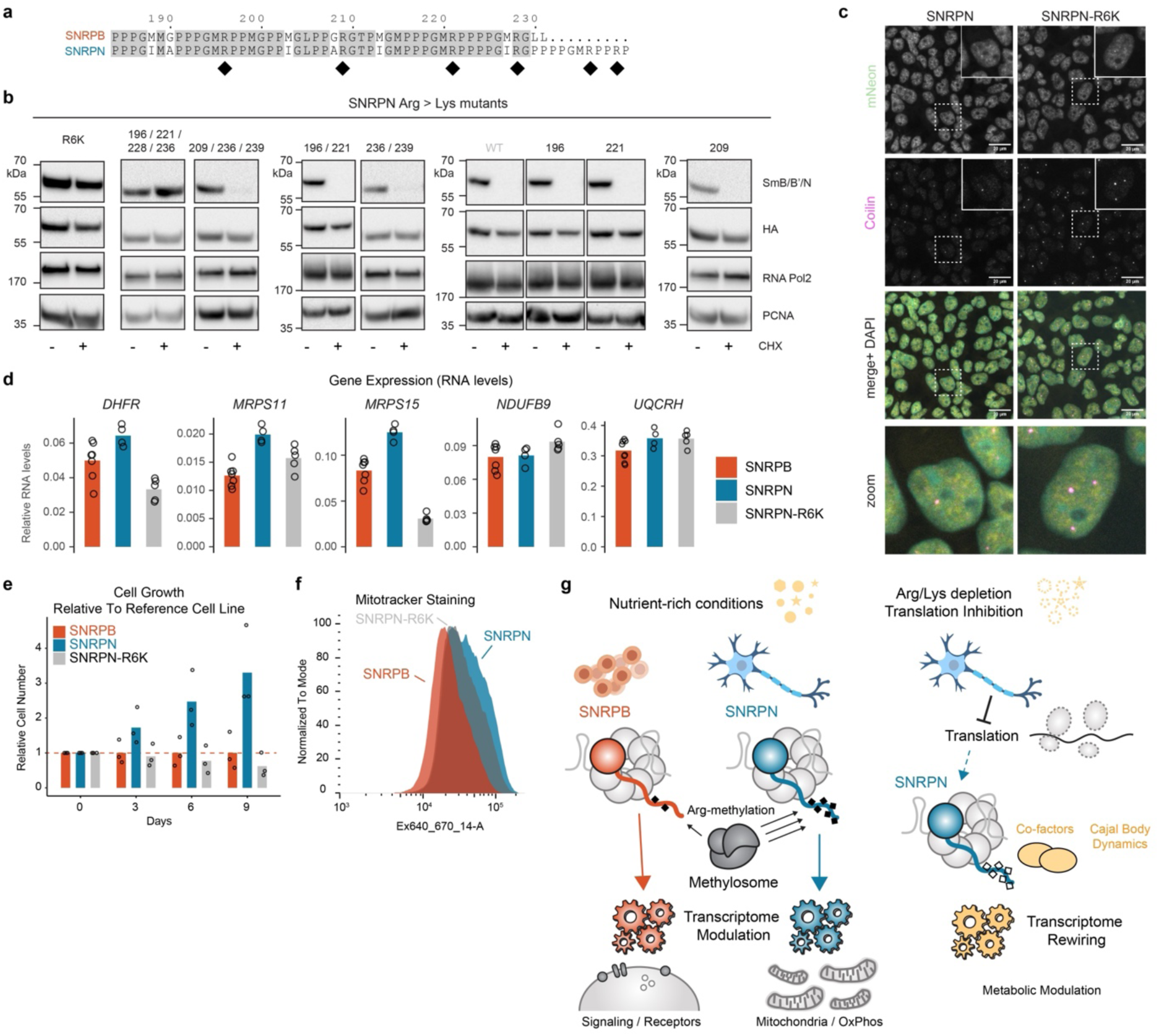
CHX resistance upon arginine mutation in SNRPN. (a) Sequence Alignment of the SNRPB/N C-termini with Arginine-residues highlighted with a black diamond. (b) Cropped immunoblots of cells stably expressing Arginine-to-Lysine SNRPN point mutants (endogenous *SNRPB* present). Cells were treated with CHX or Ethanol Control (- CHX). In R6K, all arginines marked with a diamond in (**a**) are mutated. RNA Pol2 and PCNA serve as loading controls. HA-antibody shows overall protein stability differences. The R6K experiment was replicated twice, all other mutants were assessed once. Endogenous *SNRPB* was present. (c) Representative immunostaining of Coilin (magenta) with mNeon fluorescence from tagged transgenes (yellow) in *SNRPN*- or *SNRPN-R6K*-expressing cells (endogenous *SNRPB* is knocked out). DAPI in cyan, scale bar 20 μm. Areas marked with a dashed square are magnified in the “zoom” panel. The experiment was conducted two independent times and the images shown are maximum projections of z-stacks. (d) RT-qPCR quantification of RNA levels of the indicated genes identified as DE between *SNRPB-* and *SNRPN*-cells in RNA-seq (Fig. 1). The barplot represents the mean levels normalized to *RPLP0*. Overlaid data points represent biologically independent replicates. (e) Bar plot showing long-term cell growth differences between *SNRPB-*, *SNRPN-*, and *SNRPN-R6K* cells (endogenous *SNRPB* knocked out). Because cells needed to be split during the course of the experiment a co-culture approach was chosen. For this, equal numbers of mNeon-tagged *SNRPB/N/R6K*-cells and non-fluorescent HEK293T reference cells were mixed, and their relative abundance was measured by flow cytometry at the indicated time points. Abundance values were normalized to the mean growth of *SNRPB*-cells at each time point, and overlaid data points represent biological replicates. (f) Representative flow cytometry histogram showing mitotracker staining intensities in *SNRPB/N/R6K*-cells. Unstained controls, second replicate and gating is shown in **Supplementary Data 2**. (g) Graphical abstract summarizing the findings. Methylated arginines are shown as black diamonds.

After transgene integration, we again knocked-out the endogenous *SNRPB* gene, so that cells relied exclusively on the mutant. Using Western blot, we found that, different from wild-type SNRPN, the R6K mutant protein no longer responded to CHX treatment (**Fig. 7b**). Overall protein stability was unaffected as detected by the HA-tag. Immunofluorescence microscopy showed that the R6K-mutant, like the wild-type protein, localized mostly to the nucleus and showed foci overlapping with Cajal bodies (**Fig. 7c**). CHX-resistance was also observed upon mutation of four specific arginines (R196, R221, R228, R236), but single (R196, R221, R209), double (R196/R221, R236/R239), or triple (R209/R236/R239) mutants behaved like wild-type SNRPN (**Fig. 7b**). This indicates that the two C-terminal arginines unique to SNRPN (R236/R239) that are absent in SNRPB due to its shorter C-terminus, are not solely responsible for the observed responses (**Fig. 7b**). To analyze gene expression in the SNRPN-R6K mutant, we quantified RNA levels of key SNRPN targets (**Fig. 1**) by RT-qPCR and found that gene-expression patterns in the mutant now resembled those of the SNRPB-cells (**Fig. 7d**). We also observed that, during prolonged culturing (one week or more), *SNRPN*-cells began to outnumber *SNRPB*-cells, a phenotype that was reverted in the *SNRPN-R6K*-cells (**Fig. 7e**). Lastly, we investigated mitochondria by MitoTracker staining followed by quantification with flow cytometry, and found that the R6K mutant displayed an intermediate phenotype, showing mitochondrial numbers that lie between *SNRPB*-cells (fewer mitochondria) and *SNRPN*-cells (more mitochondria) (**Fig. 7f**). Taken together, these findings indicate that the arginine residues in the C-terminus of SNRPN are important for its unique ability to modulate cellular metabolism (**Fig. 7g**).

## Discussion

When gene duplicates are highly similar, they are often viewed as being redundant. However, our work on the SNRPB/SNRPN paralogues demonstrates that in an essential spliceosomal subunit, minimal sequence divergence can be the basis of metabolic subfunctionalization, rather than merely providing backup. Despite near-complete overlap in canonical Sm/snRNP interactions, which provide essential splicing functions, expression of SNRPN was sufficient to “reprogram” cells. This is associated with a different transcriptome and metabolome along with increased mitochondrial mass, and enhanced respiratory capacity. More specifically, SNRPN expression led to higher steady-state levels of long, transposon-rich transcripts encoding metabolic enzymes and nuclear-encoded mitochondrial ribosomal subunits. This metabolically focused gene expression program renders cells more responsive to translation inhibition and amino acid restriction.

These findings are interesting in the context of the tissue-specific expression of SNRPN, which is highly expressed in the brain ^9–11^ (**Extended Data Fig. 1a**). The brain has long been a major focus of splicing research because many neuronal genes undergo extensive alternative splicing. The neuroligin-neurexin complex, which functions in cell-cell-adhesion, is one well-known example where different isoform usage specifies neuronal subtypes by shaping trans-synaptic contacts ^46^. On this rationale, it initially seemed appealing to expect a specialized role for SNRPN in brain-specific alternative splicing ^39,47^. At the same time, this idea is difficult to reconcile with SNRPN being a core Sm subunit of the spliceosome, which acts on essentially every intron-containing transcript. Over the last years, however, mitochondria and metabolic programs have moved from being classified as “housekeeping” to drivers in neuroscience. Mitochondrial activity and metabolic processes connect to proliferation, differentiation, and neuronal identity during development well as the adult brain ^48,49^. Our data give a new perspective on how RNA processing of metabolic and growth-associated programs in a context-dependent manner could shape the development of the brain.

Mechanistically, these growth and metabolic effects can be traced to a surprisingly small set of arginine residues in the proline-rich C-terminus of the SNRPB/N proteins. SNRPN shows stronger engagement with the methylosome complex compared to SNRPB and a distinctive arginine methylation pattern in this region. Arginine mutation renders SNRPN resistant to CHX treatment and reverts SNRPN to a SNRPB-like molecule. These results indicate that this short, methylation-sensitive module is an important determinant of nutrient responsiveness in an otherwise “redundant” Sm protein, which is largely identical with regards to the major interaction partners.

A key question is how such distinct methylation patterns arise at the molecular level, if four of the six substrate arginine residues are conserved between the two paralogues. One possibility lies in the immediately surrounding residues, which differ between SNRPB and SNRPN - for example, SNRPN Ile227 and Ala208 - as well as the extended C-terminal tail unique to SNRPN that contains two of the six arginines. The SNRPN tail as a whole, rather than individual sites, could thus be a better substrate for the methylosome complex. Because arginine methylation can exist in multiple states (mono-, di-symmetric, di-asymmetric) that are catalyzed by different enzymes, crosstalk between methylation pathways may also influence the differential modification patterns observed by us. Proline-Glycine-Methionine (PGM) motifs - which occur in the SNRPN C-terminus - can be methylated by two different enzymes, PRMT5 and CARM1, and there is extensive crosstalk between different arginine methylation states, particularly when one methylation axis is perturbed ^50^. This idea is supported by our observation that arginine residues in the glycine-rich regions upstream of amino acid 190 lack PGM motifs, remain unaffected by translational inhibition, and behave similar between SNRPB and SNRPN.

The upstream regulatory mechanism that makes SNRPN change its PTM pattern upon translational inhibition likely revolves around the mTOR kinase. This complex is well known for sensing availability of amino acids like arginine or leucine and being responsive to translational status ^51–53^. Previous studies have investigated connections between mTOR activity and the regulation of pre-mRNA splicing. In particular, subsets of mTOR-responsive genes, including those involved in lipogenesis, are regulated when mTOR activates the SRPK2 kinase. SRPK2 then phosphorylates SR family splicing factors, which can then promote intron excision of metabolic genes ^54–56^. Our work identifies arginine methylation of

SNRPN as an additional post-translational modification within the mTOR-responsive axis, one that appears to exert rather global effects on splicing efficiency and impacts metabolic pathways. The adaptor PHIP/BRWD2 could be a key player in that axis as it is nutrient-responsive ^41^, while at the same time gaining interactions with SNRPN upon translational inhibition.

How can the differential methylation status of SNRPB/N change splicing and/or the transcriptome? Recent work demonstrated that PRMT5 inhibition or key arginine mutations alter SNRPB interactions with chromatin and nascent RNA, thereby regulating the chromatin release of slowly spliced, intron-rich transcripts through arginine methylation ^25^. In that study, residues R108 and R112 were identified as critical for this function. However, we find that these residues (i) do not differ in their methylation state between SNRPB and SNRPN (**Fig. 3**), (ii) are not altered upon CHX treatment (**Fig. 4**), and (iii) do not overlap with the region to which our observed effects map, which instead localize to the C-terminal region starting at residue 190. Together, our data and those of ^25^ indicate that distinct arginine modules within SNRPB/N confer mechanistically independent layers of splicing control.

Using FRAP, we show that translation inhibition alters SNRPN dynamics within Cajal bodies and Cajal body numbers. This is consistent with recent work showing that ribosomal proteins are important for Cajal body formation ^57^ and suggests that reciprocal feedback between different layers of gene expression (transcription, splicing, translation) could be orchestrated through nuclear bodies. Signalling pathways like phosphorylation or arginine methylation are ideal candidates that could link such processes, as they occur more rapidly than gene expression changes. Future experiments could, for example, investigate whether translation inhibition and altered dynamics result in changed RNA-binding preferences of SNRPN and SNRPB. This could be addressed using iCLIP, which would also help distinguish direct from indirect effects on gene expression after e.g. CHX treatment. In addition, it will be important to directly measure differential RNA processing for SNRPN and SNRPB targets using e.g. RNA half-life and decay measurements in cells and *in vivo* as we have only captured steady-state RNA levels in cells.

Lastly, it is worth considering the evolutionary context of our findings as only mammals possess two distinct spliceosomal paralogs, while other eukaryotes rely on a single *SNRPB/N* protein. Mammalian *SNRPN* is exclusively expressed from the paternal, but not maternal allele (i.e. imprinted - a phenomenon that in animals is found only in mammals). Genes with paternal-only expression are often involved in growth-promoting functions, while those expressed from only the maternal allele tend to have opposing roles - a pattern explained by the parental conflict theory in placental mammals ^58,59^. Our findings suggest that *SNRPN* may be imprinted because of its growth- and metabolism-promoting functions and that its dysregulation could potentially contribute to PWS, an imprinting disorder.

What is special about mammals that they need two paralogues? The brain is a prominent organ that needs constant energy supply independent of continuous nutrient availability and food intake. Endothermic, placental mammals spend a comparatively small fraction of their time feeding and tend to sleep longer than most non-mammalian vertebrates and invertebrates, increasing the temporal separation between nutrient intake and energy demand. Beyond *SNRPN* ensuring faithful brain development and growth, we therefore speculate that the duplication and functional specialization of *SNRPB/N* may support enhanced buffering of nutrient and metabolic fluctuations in the brain and/or contribute to thermogenic control after birth, when maternal nutrient supply and thermal protection cease. Such pressures may have favored the retention of two spliceosomal variants with partially distinct regulatory capacities in mammals.

Taken together, our findings argue that SNRPN functions as a brain-enriched, nutrient-responsive modulator of splicing that functions to tune metabolism rather than being a simple SNRPB substitute. We think that it will be important to revisit how SNRPN dosage and arginine methylation state might contribute to the neurodevelopmental and metabolic features of PWS and related 15q11-q13 disorders and carefully re-investigate mice lacking *SNRPN*, without affecting other genes in the PWS critical region.

## Methods

### Cloning and plasmids

Total RNA from mouse brain tissue (embryonic for *Snrpb*, adult for *Snrpn*) was converted into complementary DNA (cDNA). Full length *Snrpb* (231 amino acids) and *Snrpn* (240 amino acids) were amplified from cDNA by using primer pairs containing a 2xHA tag. The mNEON tag was amplified from a plasmid available in the Keller-Valsecchi lab. Amplified sequences were cloned into pcDNA™FRT⁄TO plasmid, generating C-terminally tagged plasmids. Note that both SNRPB and SNRPN protein sequences are 100% identical between human and mouse, and thus, using the mouse cDNA for the transgenes allowed us to distinguish the human (endogenous) SNRPB/N from the transgenes at DNA and RNA level. Point mutations of Snrpn were generated with the QuikChange Lightning Kit (Agilent Technologies, 200518), following the manufacturer’s protocol, with reaction volumes halved. N-terminally tagged full length Snrpb and Snrpn plasmids were cloned via Gibson Assembly. To achieve this, Snrpb, Snrpn, mCherry and HA-mNeon tags were amplified by PCR and a T2A sequence was added directly to the primer sequences. To generate HEK cell lines lacking endogenous SNRPB, two single guide RNAs (gRNAs) were designed to target exon2 (gRNA1, CATTTCTCCTTGCAGACGGT) and downstream of exon 7 (gRNA2, GTGCTTAATAGAGTCAAGAA) of the SNRPB locus. The corresponding gRNA oligonucleotides were cloned into SpCas9(BB)-2A-Puro (PX459) via BbsI restriction sites. This approach was intended to induce a large deletion spanning exon 2 to 7, thereby disrupting *SNRPB* expression.

### Genotyping of SNRPB knock-out

Genomic DNA was extracted using QuickExtract DNA Extraction Solution (Biozym 101094) by incubating cell pellets for 55 °C for 1 h followed by 15 min at 75°C. The *SNRPB* gene was amplified with La Taq DNA polymerase (Takara, RR042A) using the 10x Mg^2+^ buffer and primers q181 (GTATCAGAGCCATCAGAACCG) and s303 (CAAGTGCCTTCCCAAGAGGATA) with an extension of 9 min 30 s (the WT band is 9.3 kb-long). A control gene, *MSL2*, was amplified using an in-house made Taq polymerase using primers p199 (TGGCAGTTCTGTTATCAATGGTT) and p200 (GCTATCTTCGGAGTTGCTGCT) with an extension of 30 s (the WT band is 454 bp-long).

### Cell culture and inhibitor treatments

All cell culture experiments were performed in a humidified incubator at 37 °C and 5% CO2. Flp-In™ T-REx™ 293 (herein referred to as HEK293T) and their derivatives were maintained in 4.5 g/L glucose DMEM (Gibco, 11960044) supplemented with 9% fetal bovine serum (FBS) (Gibco, A5256701), 1% Penicillin/Streptomycin (Gibco, 15140122), and 1% GlutaMAX™ Supplement (Gibco, 35050061).

For western blot, qPCR, and flow cytometry, 2x10^6^ cells per well were seeded in a 6-well plate. For immunoprecipitation followed by mass spectrometry (IP-MS), 8x10^6^ cells per 10-cm plate were used. For immunofluorescence, cells were seeded on Poly-L-Lysine (Sigma-Aldrich, P1274) coated coverslips. Twenty-four hours after plating, the medium was aspirated, and fresh medium containing the following inhibitors or their respective controls were added. A 100 mg/mL stock solution of cycloheximide (CHX) (Sigma Aldrich, C7698-1G) was freshly prepared in 100% EtOH (Thermo Fisher, 10342652) for each use, and diluted in complete DMEM to a 0.1 mg/mL working concentration for 4-hour treatments. A 10 mM stock of PRMT4/CARM1 inhibitor (Cayman, Cay11033-500) in DMSO (Sigma Aldrich, D8418-100ML) was diluted to 5 μM for 27-hour treatments. A 5 mM stock of GSK591 (Cayman, Cay18354-1) in DMSO was diluted to 10 μM for 6-day treatments. A 1 mM stock of Pladienolide B (Santa Cruz Biotechnology, sc-391691) in DMSO was diluted to 100 nM for 16-hours. A 10 mM stock of hSMG1-inhibitor 11e (ProbeChem, PC-35788) in DMSO was diluted to 0.6 μM for 24-hour treatments. After the indicated timeframes, the medium was removed, followed by immediate addition of TRIZol^TM^ Reagent (Invitrogen, 15596026) to the cell culture dish, fixation for immunofluorescence staining or collection of cells for other experiments.

### Generation of transgenic HEK293T-cell lines

For transfections, the desired plasmids and the commercial Flp-recombinase expression vector pOG44 were co-transfected into Flp-In™ T-REx™ 293 cells using the transfection reagent ScreenFect®A (ScreenFect, S-3001) in Opti-MEM™ (Gibco, 31985062), following the manufacturer’s protocol for a 6-well format. After 6 hours, Opti-MEM™ was replaced with complete DMEM supplemented with 9% FBS, 1% Penicillin/Streptomycin, and 1% GlutaMAX™ Supplement. The following day, the transfected cells were collected and seeded onto 15-cm dishes containing 0.3 mg/mL hygromycin (Sigma Aldrich, H3274) in complete DMEM. Selection continued with 0.3 mg/mL hygromycin for two days, followed by maintenance in 0.15 mg/mL hygromycin. Once colonies appeared, they were picked and seeded into 96-well plates with 0.15 mg/mL hygromycin. Cells were then expanded sequentially into 48-well, 24-well, 12-well, and 6-well plates maintaining 0.15 mg/mL hygromycin throughout. When cells reached confluency in 6-well plates, hygromycin was removed, and cells were maintained in 4.5 g/L glucose DMEM supplemented with 9% FBS, 1% Penicillin/Streptomycin, and 1% GlutaMAX™ Supplement. Following colony expansion, positive clones were initially selected based on X-Gal staining and protein expression validation by Western Blot. To induce gene of interest expressions, doxycycline (Sigma Aldrich, D9891) at 100 ng/mL final concentration was added to the medium.

To generate the endogenous *SNRPB* knock-out clones, C-terminally tagged constructs (SNRPB-2XHA-mNeon or SNRPN-2XHA-mNeon) were transfected with the gRNA-containing PX459-3HA plasmids (pCK670 and pCK671) using ScreenFect®A, following the same transfection protocol as described above. Following transfection, cells were selected with 1 μg/mL puromycin. Surviving colonies were expanded, and successful knock-out clones were identified by PCR genotyping (see methods section cloning and plasmids) and confirmed by Western blot. Interestingly, we were unable to delete the endogenous *SNRPB* gene in cells expressing N-terminally tagged constructs, suggesting that N-terminal tagging compromises protein function, at least partially. Consequently, we used C-terminally tagged proteins for all subsequent functional experiments.

### X-Gal staining

Picked colonies were seeded into a 96-well plate along with HEK293T cells as a control. The next day, two different mixtures were prepared: “Mixture 1” contained 2% paraformaldehyde (Thermo Scientific, 28906) and 0.2% glutaraldehyde (Sigma Aldrich, G5882-50ML) in PBS (without Ca++ and Mg++). “Mixture 2” contained 5 μM Potassium hexacyanoferrate(II) trihydrate (Sigma Aldrich, P3289-5G), 5 uM Potassium ferricyanide(III) (Sigma Aldrich, 702587-50G), 2 μM MgCl_2_, and 1 mg/mL X-Gal in PBS (without Ca++ and Mg++). Cells were washed with PBS (without Ca++ and Mg++). After removing the PBS (without Ca++ and Mg++), 50 μL of “Mixture 1” was added to each well and incubated for 10 min at room temperature. Following two washes with PBS (without Ca++ and Mg++), 50 μL of “Mixture 2” was added. The plate was covered with aluminum foil and incubated at 37°C until a blue color developed in HEK293T cells. Colorless colonies were selected for further experiments, while blue colonies were discarded.

### Protein isolation

Cells were harvested using either Accutase (Sigma Aldrich, A6964-100ML) or a cell scraper (Starstedt, 83.3950). The cells were then centrifuged at 700g for 5 minutes at room temperature. The resulting pellet was washed with ice-cold Dulbecco’s Phosphate Buffered Saline (PBS with Ca++ and Mg++, Sigma, D8662) and centrifuged again at 700g for 5 minutes at 4°C. The pellet was resuspended in “Henikoff buffer” (0.1% SDS, 3 mM CaCl_2_, 20 mM Tris pH 8, 100 mM NaCl, 0.9% Triton X-100), supplemented with 1 μL Sm nuclease per 1 mL of buffer, 1X cOmplete™ EDTA-free Protease Inhibitor Cocktail (Roche, 11836170001), and 1X PhosSTOP™ (Roche, 4906845001). The mixture was vortexed briefly and incubated on ice for 15 minutes. EDTA was then added to a final concentration of 20 mM. The samples were sonicated using a Bioruptor Pico (Diagenode) for 5 cycles of 30 seconds ON and 30 seconds OFF at 4°C. Following sonication, the samples were centrifuged at 12,000g for 10 minutes, and the supernatant was transferred to a fresh tube.

Protein concentrations were determined using a colorimetric Bradford assay (Panreac AppliChem ITW, A6932). A 20 mg/mL stock solution of recombinant albumin (New England BioLabs, B9200S) was diluted to 1 mg/mL in the buffer used for protein isolation. Serial dilutions were prepared by adding 0, 3, 6, 9, 12, and 15 µL of the 1 mg/mL albumin stock to cuvettes containing 1 mL of Bradford reagent. Additionally, 1 µL of each isolated protein sample was added to 1 mL of Bradford reagent in separate cuvettes. A standard curve was generated by measuring the absorbance of the albumin dilutions using a DS-11 FX+ Spectrophotometer/Fluorometer (DeNovix). The concentrations of the isolated protein samples were then determined based on the standard curve. Protein extracts were diluted to the required concentrations using nuclease-free water (Qiagen, 129115) and NuPAGE™ LDS Sample Buffer 4X (Novex, NP0007) supplemented with 50 mM DL-Dithiothreitol (DTT) (Sigma, D0632-25G). The final mixture was boiled at 70°C on a shaking heat block (Eppendorf® ThermoStat C) for 10 min, snap-frozen on dry ice, and stored at -20°C.

### Immunoprecipitation

For PTM analysis cells were harvested using a cell scraper and centrifuged at 700g for 5 minutes at room temperature. The pellet was washed with ice-cold PBS (with Ca++ and Mg++), centrifuged again at 700g for 5 minutes at 4°C, and resuspended in “Henikoff buffer” (0.1% SDS, 3 mM CaCl_2_, 20 mM Tris pH 8.0, 100 mM NaCl, 0.9% Triton X-100), supplemented with 1 µL/mL Sm nuclease, 1X cOmplete™ EDTA-free Protease Inhibitor Cocktail (Roche, 11836170001), and 1X PhosSTOP™ (Roche, 4906845001). After vortexing and incubating on ice for 15 minutes, samples were sonicated (Bioruptor Pico-Diagenode, 2 cycles: 30 seconds ON, 30 seconds OFF, 4°C), then centrifuged at 16,000g for 20 minutes. The supernatant was collected, and protein concentration was measured.

For each sample, 1 mg of protein in 0.5 mL Henikoff buffer was prepared, with 5% set aside as input. Pierce™ Anti-HA Magnetic Beads (Thermo Fisher, 88836) were pre-washed in Henikoff buffer, then added to protein samples and incubated on a rotator at 4°C for 1 hour. Beads were washed three times with wash buffer (0.1% SDS, 20 mM Tris pH 8.0, 1 mM EDTA, 400 mM NaCl, 0.9% Triton-X 100) supplemented with 1X cOmplete™ EDTA-free Protease Inhibitor Cocktail, and 1X PhosSTOP™, then resuspended in 50 uL elution buffer (25 mM Tris-HCl pH 8.0, 1% [wt/vol] SDS), and boiled at 60°C for 15 min at 800 rpm. After centrifugation at full speed for 1 min at 4°C, the supernatant was collected, and beads were re-eluted with 10 μL elution buffer. The combined eluate (60 uL) was snap-frozen and submitted for Mass Spectrometry. For interaction partner analysis, the same protocol was followed except that “Henikoff buffer” and wash buffer were replaced with cell lysis buffer (250 mM NaCl, 50 mM HEPES pH 7.6, 0.1% IGEPAL, 10 mM MgCl_2_, 10% glycerol).

### Western blot

35 μg of proteins extracts were mixed with 4X loading dye, incubated at 70 °C for 10 min and then separated by regular SDS–PAGE, transferred at 300 mA for 1 h to PVDF membranes using a Bio-Rad Wet Tank Blotting System in Tris-Glycine Transfer Buffer with 10% methanol. The membranes were blocked for 30 min in 5% milk in PBS (without Ca++ and Mg++)-0.1% Tween before incubation with primary antibodies in 5% milk PBS(without Ca++ and Mg++)-0.1% Tween. Primary antibodies anti-Sm B/B’/N (mouse, Santa Cruz Biotechnology, sc-130670, 1:2000), anti-HA (mouse, Biozol, BLD-901502, 1:5000), anti-SNRPB (rabbit, Proteintech, 16807-1-AP, 1:5000), anti-RNA Pol2 (mouse, Active Motif, 39097 and 39047, 1:6000), anti-GAPDH–HRP (Thermo Scientific MA5-15738-HRP, 1:6,000), anti-PCNA (mouse, Santa Cruz Biotechnology, sc-56, 1:5000), anti-pAKT(Ser473) (rabbit, Cell Signaling Technology, 4060T, 1:5000), anti-H3 (mouse, Active Motif 39763, 1:5,000) were used overnight. Secondary HRP-coupled antibodies anti-mouse IgG HRP (Abcam, ab46540) and anti-rabbit IgG HRP (Invitrogen, A18917) were used at 1:10,000 for 1 h. Blots were developed using SuperSignal West Pico PLUS Chemiluminescent Substrate (Thermo Scientific, 34577) and imaged on a ChemiDoc XRS+ (Bio-Rad).

### Protein digestion for mass spectrometry

All samples were processed using the SP3 approach ^60^. Proteins were reduced in 5 mM DTT, alkylated in 15 mM iodoacetamide (Sigma-Aldrich, I6125) in the dark and quenched in 5 mM DTT. Enzymatic protein digestion was performed using trypsin (Sigma-Aldrich, T6567) or porcine pancreatic elastase (Promega, V189A) overnight at 37°C. The resultant peptide solution was purified by solid phase extraction in C18 StageTips ^61^.

### Proteome analysis by liquid chromatography tandem mass spectrometry

For the PTM analysis of HEK cells, peptides were analyzed using an Orbitrap Exploris 480 mass spectrometer (Thermo Fisher Scientific) coupled to an EASY-nLC 1200 UHPLC system (Thermo Fisher Scientific). Peptides were separated in an in-house packed 60-cm analytical column (inner diameter: 75 μm; ReproSil-Pur 120 C18-AQ 1.9-μm silica particles, Dr. Maisch GmbH) by online reverse-phase chromatography through a 90-min gradient of 2.4-32% acetonitrile with 0.1% formic acid at a nanoflow rate of 250 nl/min. The eluted peptides were sprayed directly by electrospray ionization into the mass spectrometer. Mass spectrometry was conducted in data-dependent acquisition mode using a top15 method with one full scan (resolution: 60,000, scan range: 300-1650 m/z, target value: 3 × 106, maximum injection time: 28 ms) followed by 15 fragment scans via higher energy collision dissociation (HCD; normalized collision energy 30%; resolution: 15,000, target value: 1 × 105, maximum injection time: 40 ms, isolation window: 1.4 m/z). Only precursor ions of +2 to +6 charge state were selected for fragment scans. Additionally, precursor ions already isolated for fragmentation were dynamically excluded for 25 s.

For all other samples, the peptides were separated via an in-house packed 45-cm analytical column (inner diameter: 75 μm; ReproSil-Pur 120 C_18_-AQ 1.9-μm silica particles, Dr. Maisch GmbH) on a Vanquish Neo UHPLC system (Thermo Fisher Scientific). Online reverse-phase chromatography was performed through a 70-min non-linear gradient of 1.6-32% acetonitrile with 0.1% formic acid at a nanoflow rate of 300 nl/min. The eluted peptides were sprayed directly by electrospray ionization into an Orbitrap Astral mass spectrometer (Thermo Fisher Scientific). Mass spectrometry was conducted in data-dependent acquisition mode using a top50 method with one full scan in the Orbitrap analyzer (scan range: 325 to 1,300 m/z; resolution: 120,000, target value: 3 × 10^6^, maximum injection time: 25 ms) followed by 50 fragment scans in the Astral analyzer via higher energy collision dissociation (HCD; normalized collision energy: 26%, scan range: 150 to 2,000 m/z, target value: 1 × 10^4^, maximum injection time: 10 ms, isolation window: 1.4 m/z). Precursor ions of unassigned, +1 or higher than +6 charge state were rejected. Additionally, precursor ions already isolated for fragmentation were dynamically excluded for 15 s.

### Proteomics mass spectrometry data analysis

Mass spectrometry raw data were processed by MaxQuant software (version 2.1.3.0) ^62^ using its built-in Andromeda search engine ^63^. MS/MS spectra were searched against a target-decoy database containing the forward and reverse protein sequences of UniProt H. sapiens reference proteome (release 2022_03; 101,732 entries), mCherry, HA::mNEON::SNRPB, HA::mNEON::SNRPN and a default list of common contaminants. Carbamidomethylation of cysteine was considered a fixed modification. The variable modification search included methionine oxidation, protein N-terminal acetylation, mono- and di-methylation on KR residues, and hydroxyproline. A maximum of 2 missed cleavages were allowed. The “second peptides” option was switched on. Minimum peptide length was set to be 7 amino acids. False discovery rate (FDR) was set to 1% at both peptide and protein levels.

For samples digested using trypsin, the trypsin/P specificity was chosen. For samples digested using elastase, the search was performed in two steps. Semi-specific cleavage at AGILSTV residues ^64^ was assigned initially without considering the variable modifications of mono-, di-methylation and hydroxyproline. To limit the search space of multiple variable modifications in a semi-specific setting, the detected proteins from the first search served as a reduced sequence database for a second search in which mono-, di-methylation on KR residues and hydroxyproline were considered. Another potential modification of arginine in consideration is citrullination, which leads to de-imination of the arginine side chain. Extensive efforts, including manual spectral validation, were made to search for it in our spectral data. The mass shift (+0.9840 Da) caused by citrullination on arginine is exactly the same as that by de-amidation of asparagine/glutamine, which could be endogenous of origin or introduced during sample preparation. This presented a challenge to identify citrullination on SNRPB/SNRPN unequivocally in our data.

To quantify the detected PTMs, the intensities of peptide evidences associated with each detected PTM site were summed and then normalized according to the overall detected peptide intensities in each sample, assuming the overall peptide intensities were similar across the test conditions.

Protein quantification was performed using the MaxLFQ algorithm ^65^ skipping its default normalization option. Minimum LFQ ratio count was set to one. Both the unique and razor peptides were used for quantification. Differential expression analysis was performed in R. Reverse hits, potential contaminants and “only identified by site” protein groups were first filtered out. Data normalization was performed on the log-transformed data using median-centering. Proteins were further filtered to retain only those detected in at least 3 out of the 4 replicates in either group of each comparison. Following imputation of the missing LFQ intensity values, a linear model was fitted using the limma package in R ^66^ to assess the difference between the two groups for each protein, with adjustment for multiple testing using the Benjamini-Hochberg approach ^67^. The visualization of the results were performed in R. To classify proteins based on subcellular localization, the list of nuclear proteins was obtained from Uniprot (accessed on 4.4.2025, uniprotkb_nucleus_AND_reviewed_true). To avoid artificially amplifying differences between control and CHX treatment conditions, log2FC values <0 (derived from comparisons of Control versus untagged or CHX versus untagged cell lines) were set to zero prior to calculating treatment-specific differences.

### Metabolomics Sample Preparation and Reagents

The metabolomics experiment was conducted with two independent *SNRPB*-cell clones (*n*=8 replicates in total, 4 for each clone) and two independent *SNRPN*-cell clones (*n*=8 replicates in total, 4 for each clone). Cells were harvested from culture dishes by first removing the culture medium and washing once with 1 mL room temperature (RT) DPBS (1X) (14190-094, LOT 2902925) to eliminate residual medium. Cells were detached by gentle pipetting. The cell suspension was collected and cells were counted in this 1 mL DPBS using Cell counter (Denovix, S-05952) brightfield imaging; trypan blue exclusion was used only for samples with low cell numbers. A total of 1 × 10^6^ cells was transferred to a new 1.5 mL DNA LoBind Eppendorf tube (Eppendorf, 022431021) and centrifuged at 3,500 × g for 3 min at RT. After carefully removing the supernatant, the resulting cell pellet was snap-frozen on dry ice and stored at −80 °C until further processing.

The pellet was resuspended in lysis buffer (80% methanol / 20% water) pre-cooled to −20 °C and vortexed vigorously. The samples were incubated at −20 °C for 2 h and then centrifuged at maximum speed for 15 min at 4 °C. The supernatant was carefully transferred to a new tube without disturbing the pellet, snap-frozen in liquid nitrogen, and stored at −80 °C until analysis. All solvents used for this study were LC-MS grade. Water (Cat.No. 701074), acetonitrile (Cat.No. 701881), methanol (Cat.No. 701091) were bought from PanReac AppliChem GmbH (Darmstadt, Germany) and formic acid was bought from Merck KGaA (Darmstadt, Germany). We acquired NIST® SRM® 1950 from Merck KGaA (Darmstadt, Germany). The Kinetex 2.6 μm F5 100 Å, LC Column 150 x 2.1 mm was bought from Phenomenex (Aschaffenburg, Germany). Eppendorf Safe-lock tubes 1.5 ml were bought from Eppendorf (Hamburg, Germany). Sapphire pipette tips were bought from Greiner Bio-One GmbH (Frickenhausen, Germany).

### Metabolomics analysis by LC-MS/MS

Supernatants were transferred to a new tube and evaporated under constant nitrogen flow. Dried metabolites were resuspended in 10μl water. The equivalent of 75 000 cells were analyzed by liquid chromatography-mass spectrometry (LC-MS) using an Agilent 1290 Infinity II coupled to a SCIEX ZenoTOF 7600. Metabolite separation was performed on a Phenomenex Kinetex F5 column (2.6 μm, 2.1 mm × 150 mm) using a linear gradient elution. The mobile phase initially consisted of 0.1% formic acid (FA) in water at a flow rate of 0.2 mL/min. Over 2.1 minutes, the gradient transitioned to 0.1% FA in 95% acetonitrile (ACN)/5% water, reaching this composition at 6.5 minutes and maintained until 8.15 minutes. Subsequently, the flow rate was increased to 1.0 mL/min while maintaining the same solvent composition for one minute. At 9.3 minutes, the solvent was switched back to 0.1% FA in water (100%) and held for 1 minute. The flow rate was then reduced to 0.2 mL/min at 10.45 minutes, followed by a column wash until 11.0 minutes. Mass spectrometric detection was conducted in positive ion mode, with full scan MS spectra acquired over an m/z range of 70–1000. Data-dependent acquisition (DDA) was employed to fragment the top 11 precursor ions per cycle. Each precursor was selected up to two times before being dynamically excluded for 6 seconds. MS/MS spectra were recorded across a mass range of 40–1000 m/z, with Zeno pulsing enabled. LC-MS data was processed in MS-DIAL (ver. 5.5.250221) for metabolite annotation using untargeted spectral matching against the public ESI(+)-MS/MS library from authentic standards and an in-house spectral library ^68^. Reference matched metabolites were manually curated and used for further statistical analysis.

### Metabolomics data analysis

The analysis was performed using MetaboAnalyst 6.0 platform^69^ available at https://www.metaboanalyst.ca. Metabolite intensities were normalized by median, log10-transformed, and mean-centered. The two independent *SNRPB/N*-cell clones in this experiment were treated as replicates. One replicate of *SNRPN*-cells (sample CKV27) showed a marked deviation from all other samples in the PCA analysis and was therefore excluded from downstream analyses. Statistically significant metabolites were defined using an FDR < 0.05. Pathway analysis of significantly altered metabolites was conducted in the MetaboAnalyst 6.0 Pathway Analysis module using metabolite names and default parameters with the Homo sapiens KEGG pathway library. The results were visualized in R.

### RNA isolation and RT-qPCR

RNA was extracted using the Direct-zol RNA MicroPrep Kit (Zymo Research, R2062) with 300 µL TRIzol (Fisher Scientific, 15-596-026). Quantification was carried out using a nanodrop. For qPCR quantification, cDNA was synthesized with oligo(dT) primers with a reverse transcriptase enzyme produced by the IMB Protein Production Core Facility. qPCR was done in a 7 µL reaction at 300 nM final primer concentration with the FastStart Universal SYBR Green Master (ROX) mix (Roche, 04913850001). The reaction was run in the LightCycler 480 2 (Roche, 05015243001) with the cycling conditions as recommended by the manufacturer. Serial dilutions were used to test primers and all showed results in the range of 90-105% for their efficiency without amplification in water control.

### qPCR primers

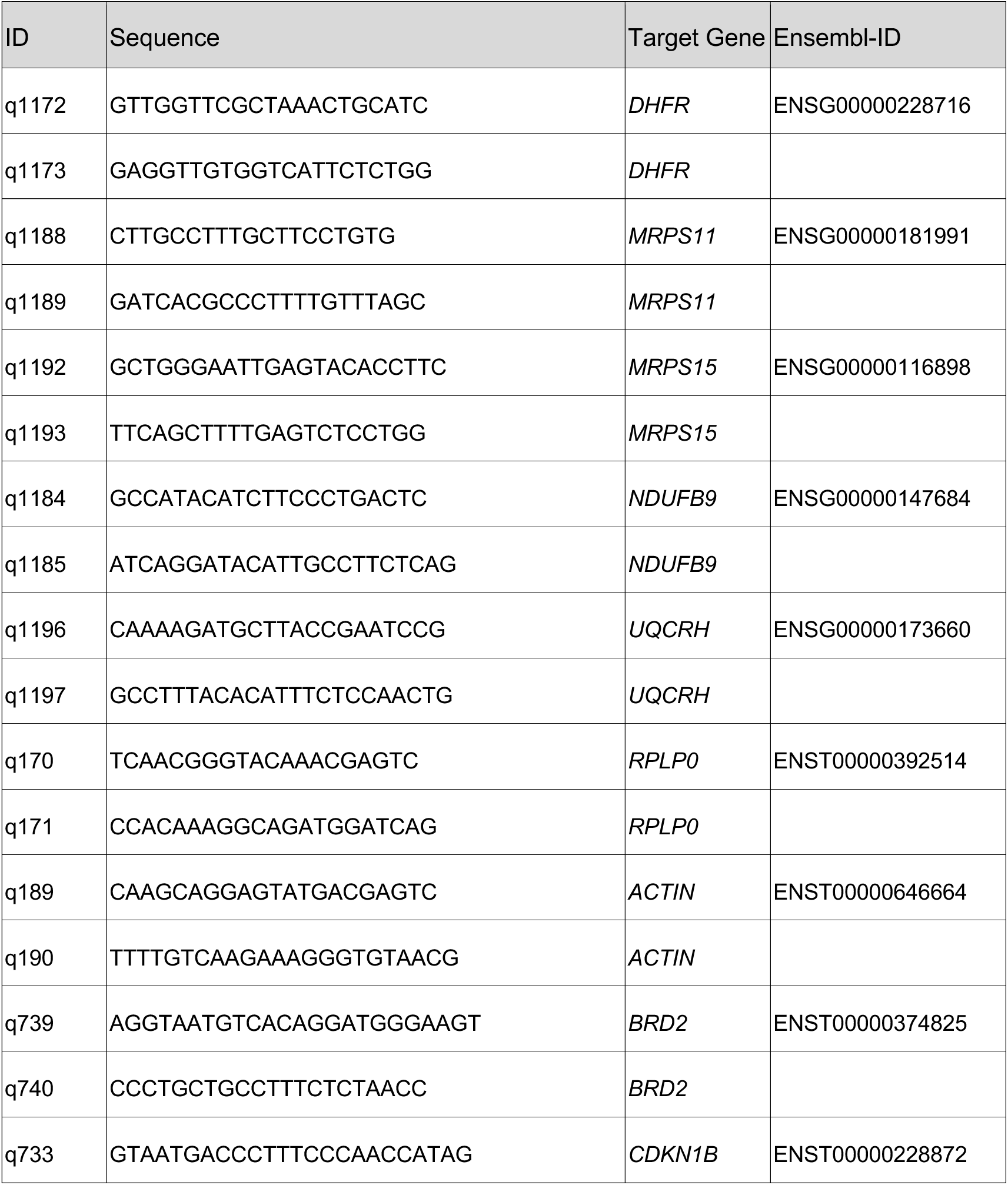
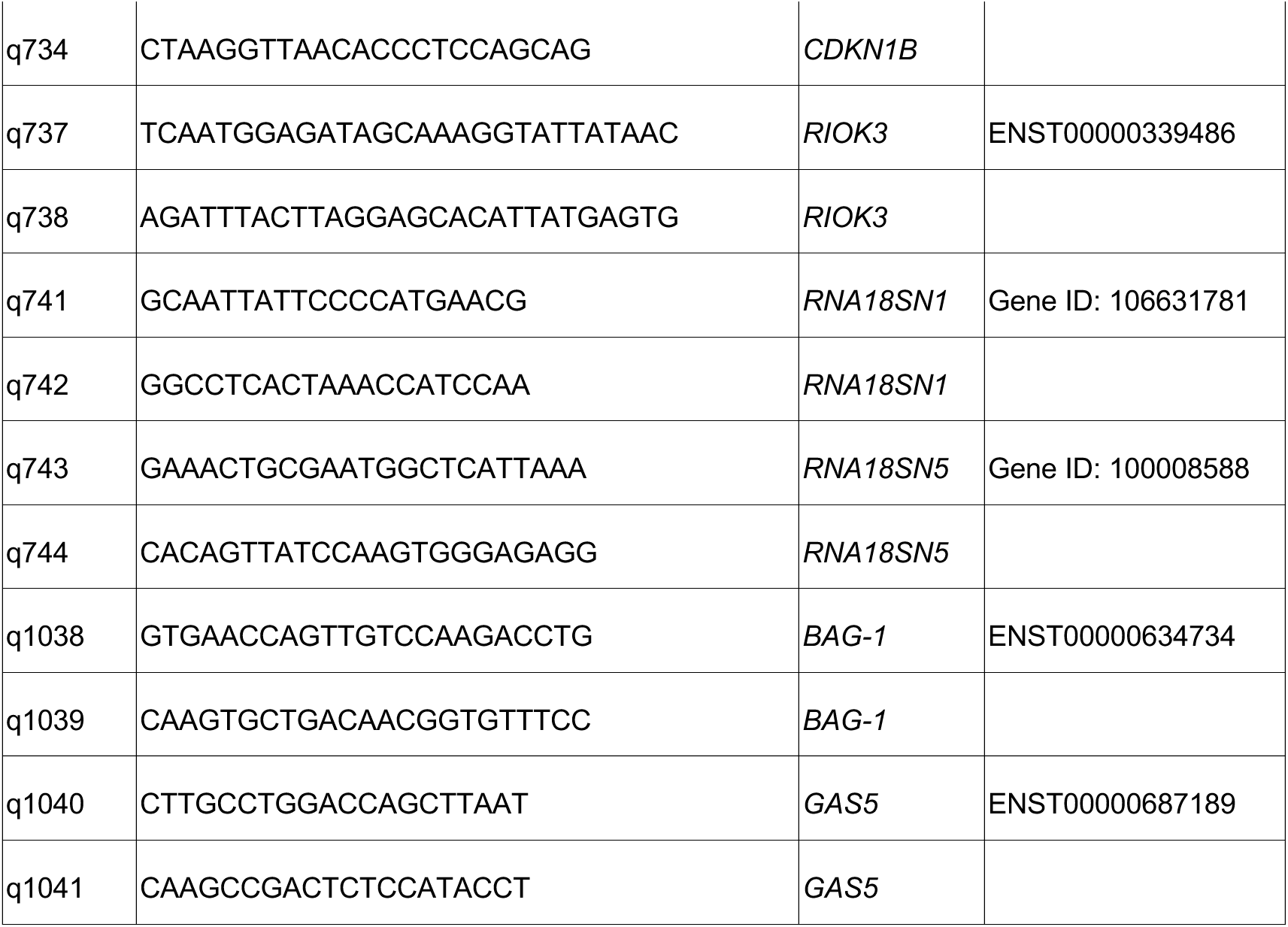

### poly(A) RNA-seq library generation

NGS library preparation was performed with Illumina’s Stranded mRNA Prep Ligation Kit following Stranded mRNA Prep Ligation ReferenceGuide (April 2021) (Document 1000000124518 v02). Libraries were prepared with a starting amount of 200ng and amplified in 10 PCR cycles. Two post PCR purification steps were performed to exclude residual primer and adapter dimers. The libraries were pooled in equimolar ratio and sequenced on a NextSeq 2000 P3 flowcell (200 cycles) in paired end mode yielding an average of 30.8 million read pairs per sample.

### RNA-seq data analysis

Raw data demultiplexing has been performed using Illumina’s bcl-convert software v.4.1.5 and overall sequence quality and rRNA content is assessed with FastQC v0.11.9 (https://www.bioinformatics.babraham.ac.uk/projects/fastqc/) and FastQ Screen v0.15.2 ^70^. Reads were trimmed for Illumina adapter sequences using cutadapt v.4.0 ^71^ and subsequently mapped to Encode reference genome GRCh38.p13 (release 43) with STAR v2.7.10b ^72^ with default parameters (except --outFilterMismatchNoverLmax 0.04 and --outFilterMismatchNmax 999). Only uniquely mapped reads were used for downstream analysis. CPM-normalized coverage signal tracks (bigWigs) of primary alignments were generated using DeepTools v3.5.1 ^73^. Read summarization at gene level (exon reads counted) is performed using Subread featureCounts v.2.0 ^74^ with default parameters.

The rMATS tool v.4.1.2 ^19^ was used for pairwise differential splicing analysis considering exon skipping (SE), alternative 5’ splice sites (A5SS), alternative 3’ splice sites (A3SS), mutually exclusive exons (MXE) and retained introns (RI) events. Low coverage events with an average coverage per condition < 5 reads were removed. Events were tested based on junction and exonic reads and were considered significant if showing a minimum PSI (percentage spliced in) change of 10% with an FDR cutoff of 0.01.

The pairwise differential expression analysis between the experimental groups testing for a log2FC different from zero was performed in R v.4.3.2 with R-package DESeq2 v.1.42.0 ^75^. DE genes with padj <0.01 were considered significant. The GOTerm analysis was conducted in R using the enrichR ^76^ package and the KEGG_2021_Human database. To analyze the gene features, the GenomicFeatures ^77^ package in Bioconductor with the TxDb.Hsapiens.UCSC.hg38.knownGene database was used. A control group of genes was generated by using the summary function and sampling genes that are padj>0.01 (not significantly changing) and exhibit a mean expression count between 233.1 (1st quartile of SNRPB DE genes) and 2883.6 (3rd quartile SNRPN DE genes). Statistical significance for the gene features (exon / intron numbers, lengths, expression levels) was evaluated with an unpaired, two-sided Wilcoxon Rank Sum test. For the information on repeats, we used the hg38 file downloaded from Repeatmasker Database 24. May 2025 at 17:50 // https://www.repeatmasker.org/species/hg.html. Their presence in the introns of the SNRPB-DE, SNRPN-DE or control gene (not DE) group were analyzed with the GRanges ^78^ packages. The enrichment of given repeats / transposons in introns was obtained by calculating the expected number of transposons based on the cumulative intron length of interesting gene groups (i.e. DE genes or differentially spliced genes). To analyze the evolutionary conservation, we used ENSEMBL biomart to capture the scerevisiae_homolog_ensembl_gene (Eukaryota), dmelanogaster_homolog_ensembl_gene (Metazoa), drerio_homolog_ensembl_gene (Vertebrata), mmusculus_homolog_ensembl_gene (Mammalia), ptroglodytes_homolog_ensembl_gene groups (Primates) from the gene groups of interest. Statistical significance was evaluated with a chi-square test. To evaluate the 5’ and 3’ splicing strengths, we used MaxEntScan ^20^ with the scripts available on http://hollywood.mit.edu/burgelab/software.html.

### Ribosome profiling data analysis

Ribosome profiling (GSM7430249, GSM7430250) and the corresponding RNA-seq (GSM7430259, GSM7430260) data generated by ^79^ was used for analysis with default settings unless specified otherwise. The datasets were downloaded using sratoolkit v2.11.0. Adapter trimming and quality control was done with fastp v0.23.2. Next, reads were mapped to human genome (ensembl v110, GRCh38.p14) with STAR v2.7.3a --outSAMtype BAM SortedByCoordinate, bam files were indexed with samtools index v1.10 and coverage tracks were generated with deepTools v3.5.5 bamCoverage. The tracks were visualized with IGV (v. 2.17.4). For the data across mouse tissues, the normalized TPM values were downloaded from GSE94982 ^80^.

### Long term cell growth measurements by flow cytometry

To evaluate subtle differences in cell growth rate over a time frame of several days, we co-cultured cells-of-interest (mNeon-tagged SNRPB/N/R6K) together with a reference cell line (parental HEK293T, non-fluorescent). For this, on day 0, 400,000 Trypan blue (Invitrogen T10282) negative (live) cells of HEK293T cells were co-cultured together with 400,000 live SNRPN, SNRPB or SNRPN-6KR cells. On day 3, cells were detached, counted (DeNovix CellDrop FL) and 1 million live cells were re-seeded. On day 6, cells were detached, counted and 1 million live cells were re-seeded. Lastly, on day 9, cells were detached, counted and 1 million live cells were re-seeded. The number of mNEON+ and non-fluorescent cells were determined on day 0, 3, 6 or 9 by flow cytometry. For this, after detaching, cells were spun down (1000 rpm, 3 min), washed once with PBS containing Ca++ and Mg++ and centrifuged again. Pellets were resuspended in PBS without Ca++ and Mg++. The cell suspension was analyzed with BD LSRFortessa SORP cytometer. Side scatter (SSC) and forward scatter (FSC) were used to exclude debris, comparing FSC-area against FSC-height was used to exclude doublets and Helix NP NIR at a final concentration of 5 nM was used for live-dead cell exclusion. Gates for mNeon+ signal were set up on a negative control (HEK293T) baseline fluorescence. Flowjo v10.10 was used for analysis (Gatings in **Supplementary Data 2**).

### Oxygen consumption rate

Live-cell respiration was measured in real-time using a Resipher instrument (Lucid Scientific). 30’000 cells were cultured in 96-well plates (Corning® 96-well Clear Flat Bottom Polystyrene TC-treated plates). The 32 needle resipher sensor lid (10.1 mm probe length, Lucid Scientific, NS32-101A) was pre-equilibrated with complete media in a cell-free 96-well microplate at 37 °C and 5% CO2 for at least 3 hours. After the equilibration the lid was moved to the plate containing the cells in regular media. The oxygen concentration in the media was measured with an operating height between 1,500 and 1000 µm for 24 hours. During media changes (1 uM oligomycin treatment, arginine depletion), the sensor lid was temporarily moved to an empty equilibrated plate while compounds were added to the cell plate. The lid was then immediately repositioned back onto the cell plate, allowing continuous measurement without interruption. This approach enabled uninterrupted oxygen consumption monitoring over 56-80 hours total (24 hours initial measurement, followed by 24-32 hours after each media change). O2 flux values (fmol/mm^2^/s) were extracted from O2 µM values with normalization every 15 min. For plotting, flux data were imported into R studio (Version 2025.09.1+401 (2025.09.1+401)) from CSV files.

For comparison of SNRPB- with SNRPN-cells, three independent experiments were performed, where the measurements were filtered to a defined steady-state window (16 - 24 hours post-plating) and the mean of each well was calculated. After excluding wells with negative values, the distributions of mean values were displayed as boxplots using *ggplot2* stratified by cell line and experiment. Statistical significance was evaluated using a linear mixed-effects model implemented in the *lme4* and *lmerTest* packages in R. Mean values were modeled as a function of cell line class (SNRPB versus SNRPN) as a fixed effect, with experiment included as a random intercept to account for inter-experiment variability (MeanValue ∼ Class + (1 | Experiment)). Model parameters were estimated using restricted maximum likelihood (REML), and *p*-values for fixed effects were computed using Satterthwaite’s approximation for degrees of freedom.

For the Arg-switch experiments, time-resolved flux data were aligned relative to the treatment period (media switch) and an analysis window spanning -8 to +10 h relative to the treatment midpoint was extracted. Data were aggregated into hourly bins by averaging measurements within each bin for each well/sample. An additional treatment bin (“T”) was computed as the mean value during the treatment interval. Outlier samples with extremely low baseline values (<5 units in the - 8 hour bin), consistent with insufficient cell numbers, were excluded. To visualize the flux data across wells and experiments ribbon plots were generated for *SNRPB* and *SNRPN* cell lines by computing the mean response across samples within each condition at each time bin. Variability was represented as the standard error of the mean (SEM), displayed as a shaded ribbon around the mean. Lastly, we also performed endpoint comparisons with the 10 hour post-treatment timepoint where the values were extracted and visualized using boxplots with overlaid individual data points. Comparisons between conditions were performed using Wilcoxon rank-sum tests.

### Immunofluorescence stainings of HEK293T cells

Coverslips (Starstedt, 83.1840.002) were coated with Poly-L-Lysine (Sigma-Aldrich, P1274) according to the manufacturer’ instructions. Cells were seeded onto cell culture dishes containing the coated coverslips. Following the inhibitor treatment, cells were washed twice with PBS with Ca++ and Mg++ (Gibco, 14040117). The cells were then fixed with 4% paraformaldehyde (Thermo Scientific, 28906) in PBS (without Ca++ and Mg++) for 15 minutes at room temperature. After three washes with PBS (without Ca++ and Mg++), they were incubated in 0.25% Triton X in water for 3 minutes, followed by two PBS (without Ca++ and Mg++) washes. Blocking was performed using 1% BSA (Pan Biotech, P06-1391500) in PBS (without Ca++ and Mg++) for 30 minutes at room temperature. After two additional washes, samples were incubated overnight at 4°C with the respective primary antibody: anti-Sm B/B’/N (mouse, Santa Cruz Biotechnology, sc-130670, 1:500) or anti-Coilin (mouse, Santa Cruz Biotechnology, sc-55594, 1:300), diluted in 0.1% BSA in PBS (without Ca++ and Mg++). Following three washes with PBS (without Ca++ and Mg++), samples incubated with the secondary antibody (Donkey anti-Mouse IgG (H+L) Highly Cross-Adsorbed Secondary Antibody, Alexa Fluor™ 555, Invitrogen, A-31570, 1:400), diluted in 0.1% BSA in PBS (without Ca++ and Mg++), for 1 hour at room temperature. They were then washed twice with PBST (1X PBS (without Ca++ and Mg++), 0.2% Tween® 20, Merck, P1379-1L) and stained with DAPI (1:2000 dilution in PBS (without Ca++ and Mg++), Invitrogen, 10184322) for 10 minutes. After a brief wash with water, the coverslips were mounted using ProLong^TM^ Diamond Antifade Mountant (Invitrogen, P36965).

### MitoTracker and Phalloidin staining for confocal microscopy

Cells were plated at 80% confluency in a Ibidi 3 well chamber (Ibidi 80381, no coating) with 0.025 μg/mL doxycycline-containing media per well. The next day, the cells were incubated with MitoTracker™ Red CMXRos Dye (Thermo Fisher, M46752) at 500 nM in media for 1 hour. Media containing dye was then removed, washed and cells were allowed to destain in fresh media for at least 1h. Cells were then fixed in 4% Formaldehyde in complete media at 37°C for 20 min. After fixation slides were washed twice with PBS without Ca++ and Mg++ and stored at 4°C in PBS until further processing. Cells were treated with DAPI and 1:1000 Phalloidin-647(Thermo Fisher, A30107) for 20 min, washed once in PBS and mounted in Ibidi mount (Ibidi, 50001).

### MitoTracker staining and analysis by flow cytometry

Mitotracker staining was performed following the same protocol as for microscopy, using a 60 mm plate containing 2 million cells per genotype. After the 1-hour destain period, cells were detached using Accutase (5 min incubation at 37°C, 5% CO₂). The detached cells were carefully resuspended and transferred to a falcon tube, then centrifuged at 1000 rpm for 3 min. The supernatant was removed and the cell pellet was washed once with PBS. The cells were then transferred to an Eppendorf tube to ensure complete media removal and centrifuged again at 1000 rpm for 3 min. The final cell pellet was gently resuspended in 4% Formaldehyde-PBS solution and fixed for 20 min on a nutating shaker. The fixed cell pellet was washed twice in PBS, and cells were kept in PBS in the dark until measurement. The cell suspension was analyzed with BD LSRFortessa SORP cytometer. Side scatter (SSC) and forward scatter (FSC) were used to exclude debris and comparing FSC-area against FSC-height was used to exclude doublets. Gates for Mitotracker signal were set up on a negative control (unstained *SNRPN*-cells) baseline fluorescence. Flowjo v10.10 was used for analysis (Gating in **Supplementary Data 2**).

### Mouse tissue sectioning and stainings

Tissues were sliced coronally at a thickness of 20 µm using a Leica cryostat and post-fixed in 4% paraformaldehyde (PFA) for 15 minutes. To ensure maximal antibody penetration, immunostainings were performed on free-floating sections. Blocking of nonspecific binding sites and tissue permeabilization were carried out simultaneously by incubating the sections with filtered 3% bovine serum albumin (BSA; Roth No. 8076.5), 5% normal donkey serum (NDS, Millipore, S-30), 0.1% TritonX-100 (Sigma-Aldrich, T8787) 1.5 hours at room temperature on a rocking shaker. Sections were then incubated with antibodies against Coilin (Genetex, GTX55575), Sm B/B’/N 12F5 (Santa Cruz, SC-130670), NeuN (Synaptic Systems, 266004), each at a dilution 1:200 in 5% NDS, 0.1% TritonX-100 for 60 - 72 hours at 4°C on a shaker. Sections were subsequently washed three times for 15 minutes with 0.1% TritronX-100 PBS, followed by incubation with 1:400 Alexa Fluor-555, Alexa Fluor-488 and CyF405S-conjugated secondary antibodies (1:400 dilution) at room temperature. Sections were then incubated with SPY 650-DNA (Spirochrome, 1:1000) for 20 minutes prior to mounting. Sections were mounted using ProLong Diamond Antifade (Invitrogen, No. P36965) and allowed to cure for at least 48 hours at room temperature prior to imaging.

### Mouse tissue image acquisition

Images were acquired using a Leica Stellaris 8 FALCON point-scanning confocal microscope. The 16-bit images were collected in point-scanning mode with a z-step of ∼ 0.3 μm using 63x/1.4 NA oil-immersion HC PL APO CS2 oil objective and 1.5-2x digital zoom. Fluorophores were excited sequentially using line scanning at the following wavelengths 405 nm (direct modulation 405-nm diode laser), and 488nm, 555 nm, 650nm (white light laser). Emitted signals were collected in four spectral detection windows: 417-472 nm, 500-556 nm, 567-620 nm, and 662-834 nm. Images were acquired at a scan speed of 600 Hz with a voxel size of 0.1806 × 0.1806 × 0.2985 µm³ and a frame size of 512 × 512 pixels. For intensity quantification, nuclei were segmented from z-maximum projections of the SPY 650 channel using Cellpose with the pretrained CYTO2 model in Fiji. An approximate nuclear diameter of 12 µm was provided to guide segmentation. For each nucleus, the channel of interest was extracted and collapsed using a sum-intensity projection, the integrated signal was quantified per nucleus.

### Confocal Microscopy (HEK293T cells)

The MitoTracker / Phalloidin stained cells were acquired using a Leica Stellaris 8 FALCON point-scanning confocal microscope. The 16-bit images were collected in point-scanning mode with a z-step of 0.2985 µm using a 63×/1.4 NA oil-immersion HC PL APO CS2 objective and 2× digital zoom. Fluorophores were excited sequentially using line scanning at 405 nm (diode laser) and 492 nm, 593 nm, 652 nm (white light laser, WLL). Emitted signals were collected in four spectral detection windows: 425–501 nm, 504–595 nm, 605–638 nm, and 666–834 nm. Images were acquired at a scan speed of 600 Hz, with a voxel size of 0.1806 × 0.1806 × 0.2985 µm³ and a frame size of 512 × 512 pixels. Images were deconvolved using the Leica Lightning algorithm with default settings.

All other images were taken with a HC PL APO CS2 100x/1.4 NA oil-immersion objective. An area of 116 x 116 µm was imagedwith an image format of 1160 x 1160 pixels with bidirectional scanning. Confocal z-stacks were acquired with 40 z-slices spanning a total thickness of 8 µm, resulting in a voxel size of 0.1 x 0.1 x 0.2 µm. DAPI, mNeonGreen (mNg) and Alexa Fluor 555 were excited sequentially (line-switching) using 405 nm (diode), 488 nm (OPSL) and 550 nm (80 MHz pulsed White Light Laser) excitation lasers, respectively. Their fluorescence was detected using HyD hybrid detectors in photon-counting mode over wavelength ranges 415-485 nm, 500-538 nm and 560-725 nm, respectively, generating 3-channel digital images with a bit-depth of 8. 4-8 fields of view were chosen for imaging per sample, where the cells were growing in a monolayer.

### FRAP (Fluorescence Recovery After Photobleaching)

FRAP measurements were performed on live cells in ibidi 8-well microwell slides using the Leica Stellaris 8 Falcon confocal microscope, equipped with an Okolab incubation cage for maintaining the cells at 37°C and 5% CO2 during the measurements. For each FRAP measurement, a single focal plane 18.45 x 18.45 µm in size was imaged using the HC PL APO CS2 63x/1.4 NA oil-immersion objective from Leica with an imaging format of 256 x 256 pixels, resulting in a pixel size of 72 x 72 nm. Fluorescence of SNRPB/N-mNg was excited using the 80 MHz pulsed White Light Laser at 488 nm and the emission was detected in the wavelength range 500-550 nm using a HyD-S hybrid detector in photon counting mode. A 10-frame pre-bleach timelapse was recorded with a frame interval of 0.342s. Following this, a SNRPB/N-mNg focus was bleached using a high intensity of the same laser and a 50-frame post-bleach timelapse was recorded with a frame rate of 0.331s. The bleached SNRPB/N-mNg focus was then tracked over time and the mean intensity of mNg fluorescence inside the focus was used to obtain the FRAP recovery curve as described below.

Fiji v1.54p was used for all image visualization and analysis 81. All segmentations were performed on filtered 2D z-projections of the 3D images, where the filters and thresholding algorithms to be applied were determined empirically for each segmented object of interest and then applied consistently across samples.

### Image Processing and analysis

#### Detection and counting of SNRPB/N and coilin foci

Maximum z-projections were used for the detection of coilin and SNRPB/N-mNg foci. Max z-projections of the coilin channel (gaussian filter, radius = 2 pixels) and SNRPB/N-mNg channel (variance filter, r = 2 px followed by gaussian filter, r = 3 px) were thresholded using the Maximum Entropy algorithm to obtain binary masks of the foci in the respective channels ^82^. The foci in each channel were counted, and these counts were normalized to the number of cells in the field of view. In control cells not expressing SNRPB/N-mNg, only coilin foci were segmented and counted.

#### Distance-based colocalization of SNRPB/N and coilin foci

The ‘Find maxima’ function in Fiji was applied to the binary masks of coilin and SNRPB/N foci with prominence 1 to detect the pixel closest to the center of each identified focus in both channels and a Euclidean Distance Map (EDM) was calculated for the centers of the coilin foci. The value on the EDM was then read out at the center of each SNRPB/N-mNg focus to obtain the distance to the center of the nearest coilin focus. SNRPB/N-mNg foci categorized as their distance to a coilin focus (0-200 nm, 200-1000 nm and > 1000 nm).

#### Quantification of nuclear and cytoplasmic SNRPB/N

Sum z-projections of the 3D images were used for quantifying nuclear and cytoplasmic distribution of SNRPB/N signal in samples immunostained for SNRPB/B’/N. The sum z-projection of the DAPI channel (median filter, r = 2 px followed by gaussian filter, r = 5 px) was used to generate a nuclear mask using the Huang thresholding algorithm ^83^. The edges of the resulting thresholded nuclei were further eroded by two pixels to obtain an accurate binary mask of the nuclei. Similarly, the sum z-projection of the SNRPB/N-mNg channel (gaussian filter, r = 10 px) was used to obtain a binary mask of cells using the Triangle thresholding algorithm ^84^, followed by the erosion of 5 pixels. For control cells not expressing SNRPB/N-mNg, autofluorescence of the cells could be picked up in this channel and used for segmenting the cells in the same way but using the iterative intermeans thresholding algorithm instead.

A binary mask of the cytoplasm was obtained by subtracting the nuclear mask from the cell mask, and the cell mask was inverted to obtain a mask for the background. The binary masks for nuclei, cytoplasm and background were then applied to the sum z-projections of SNRPB/B’/N and SNRPB/N-mNg channels to obtain mean intensity in each of these regions. The mean background intensity was then subtracted from both the mean intensities obtained for cytoplasmic and nuclear signal.

To account for variability in fluorescence intensities between experimental replicates prepared and acquired on different days, background-subtracted mean intensities (see previous paragraph) were normalized using min–max scaling. Normalization was performed separately for each fluorescent channel and experimental replicate, with intensity values pooled across all treatment conditions and subcellular compartments. Normalized values were calculated according to the formula *x*_norm_= (*x*−*x*_min_) / (*x*_max_−*x*_min_) where *x* is the background-subtracted mean intensity of an individual measurement, and *x*_min_ and *x*_max_ represent the minimum and maximum values observed within the corresponding channel and experimental day. This transformation scales values to the range of 0-1, preserving relative differences while minimizing technical variability across experiments. Normalized intensities were used for downstream visualization and statistical analyses, including comparisons across treatments, cellular compartments (nuclear versus cytoplasmic), and experimental conditions.

#### Analysis of FRAP data

Fluorescence recovery curves for the bleached SNRPB/N-mNg foci were obtained from the recorded timelapses using Fiji, where the TrackMate plugin ^85^ was used to track the bleached SNRPB/N-mNg focus over the timelapse using the Linear Assignment Problem tracker ^86^. The mean intensity inside a circle of diameter 0.7µm centered at the focus was estimated at each frame of the timelapse (I_bleach_ (t)). To track mean intensity in the nucleus over time, a binary nuclear mask was generated from the first frame of the timelapse (gaussian filter, r = 2px) using the Huang thresholding algorithm ^83^. The binary mask was then applied to each frame of the timelapse to determine the mean intensity of SNRPB/N-mNg signal in the nucleus over time (I_nuc_ (t)). Finally, a representative region in the background was chosen to determine the mean background intensity in each frame (I_bg_ (t)).

Background correction and normalization were performed as described in the easyFRAP-web Manual ^87^. For statistical analysis, we employed a nonlinear mixed effects model with autocorrelation, implemented using the R packages *nlme* ^88,89^ and *emmeans* ^90^ (v. 1.11.1). The model was based on the exponential recovery function from easyFRAP, reparameterized to enable direct inference on biologically meaningful parameters.

Fluorescence intensities of the bleached spot and the nucleus were corrected by subtracting the background intensity,

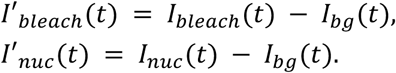

Let 𝐼′*_bleach,pre_* and 𝐼′*_nuc,pre_* denote the average pre-bleach intensities of the bleached region and the nucleus, respectively. The data were normalized in two steps (“double normalization” as described in *easyFRAP*): first, each signal was divided by its corresponding pre-bleach average, then the normalized bleach signal was divided by the normalized nuclear signal to correct for the global decrease in fluorescence due to bleaching:

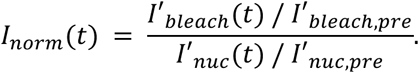

The exponential recovery model for the post-bleach dynamics is

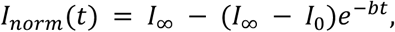

with asymptote 0 < 𝐼∞ ≤ 1, initial value 0 < 𝐼_0_ ≤ 𝐼∞, and rate 𝑏 > 0, where 𝑡 = 0 corresponds to the first post-bleach measurement. The quantities of interest are mobile fraction

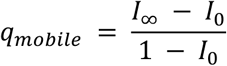

and half-value time

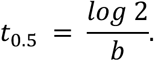

To improve interpretability and statistical inference, we reparametrized the model in terms of

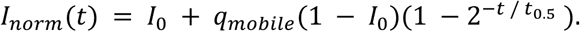

Note that, for meaningful FRAP curves, the parameters are expected to lie within the ranges 0 < 𝐼_0_, 𝑞*_mobile_* < 1 and 0 < 𝑡_0.5_ < ∞. Time series were excluded from analysis if these constraints were violated or the *nlme*-optimizer failed to converge.

The nonlinear mixed effects model included four predictive factors -– *cell line*, *treatment*, *replicate*, and individual *curve* – to analyze their influence on the model parameters 𝐼_0_, 𝑞*_mobile_* and 𝑡_0.5_. The primary factors of interest – cell line and treatment – were included as fixed effects with interaction. To capture and control for unwanted sources of variability, the nested factors *replicate* and *curve* were modeled as random effects. A diagonal covariance structure was used for *replicate* (i.e., we assumed independent effects on model coefficients), while an unstructured covariance matrix was used for *curve* (allowing correlated effects). Given the longitudinal nature of the data, an autoregressive correlation structure of order 1 (corAR1) was applied within curves to model serial correlation.

Model selection was guided by the Akaike Information Criterion (AIC) and likelihood ratio tests.

### Mice

Mice were kept according to the protocols approved by Institutional Animal Care and Use Committee, Institutional Ethics Committee and the Preclinical Core Facility at Institute of Science and Technology of Austria (ISTA). Experimental protocols were performed under the approval of the Austrian Federal Ministry of Science and Research in accordance with the Austrian and European Union animal laws (license number: BMWFW-66.018/0015-WF/V/3b/2017; GZ: 2020-0.579.989; GZ: 2024-0.698.056).

All mice used showed pathogen free status according to the Federation of European Laboratory Animal Science Associations (FELASA) recommendations. Animals were maintained and bred in the Animal Facilities of the Institute of Science and Technology of Austria (ISTA). Animals were kept on a 12:12 hours light:dark cycle, room temperature of 21 ± 1°C and relative humidity of 40% to 55%, in accordance to European Union regulations. Food (V1126, Ssniff Spezialiteaten GmbH, Soest, Germany) and tap water available ad libitum. Wild type mice in a C57BL/6J background were used.

### Intracerebroventricular injections of mice

Pups of 7 days postnatal (P7) were deeply anesthetized with Vetflurane (induced at 3% and maintained at 1-2%). After fixing the pup to the stereotaxic, the head was cleaned with 70% EtOH, Xylocain Gel 2% was applied and the skull was exposed by a minimal cut along the anterior-posterior axis and other perpendicularly to it at the level of lambda. The injected location was established by the coordinates: -0.3 to -0.5mm from Bregma in the anterior-posterior axis, ±1.00 to 1.5mm in the medial-lateral axis and 2.0 to 2.5mm in the ventral-lateral axis. For the injections, pulled glass micropipettes were used together with a Nanoject III (Drummond Scientific Company). 500nL with a 200nL/s rate for 10 cycles was injected (a total of 5µL) per hemisphere. Cycloheximide (CHX, Sigma, REF C7698-1G) at 4µg/µL concentration in NaCl 0.9% was administered in the treatment condition, while controls received a solution of NaCl 0.9%. CHX working solution was prepared fresh the day of the injections. Both conditions included 0.02% Fast Green (Sigma, REF F7252-5G). After injecting both hemispheres, tissue adhesive (SURGIBOND, REF ZG2) was applied in the edges of the wound. The pup was maintained over a warm pad during the whole procedure and until it woke up. Pups were placed back with the mother for 5 hours. Then, the pups were sacrificed by cervical dislocation and immediately processed. The brain was extracted and fixed in PFA 4% by immersion over night at 4°C. Then, embedded in OCT (Tissue-Tek OCT, Sakura, REF 4583) and frozen with dry ice. Brains were kept in -20°C until processed. Male and female sex was assigned based on Y-chromosome genotyping at P4 with primers Y-chromosome forward AGATGAAGATGCTGGTGGCACAGC and Y-chromosome reverse GACTAGACATGTCTTAACATCTGTCC GoTaq® G2 DNA Polymerase in a Ready-to-Use Master Mix (REF M782B) was used according to the manufacturer protocol and the PCR reaction (60°C).

### Evolutionary Analyses, Sequence Analyses, Alignments and Visualizations

DNA and protein sequences were retrieved from Ensembl and Uniprot, respectively. Alignments were created on Clustal Omega. Alignments were visualized with ESPript. The analysis of the amino acid sequence (structured versus disordered) was visualized with (MobiDB lite). Amino Acid Evolutionary Constrained Analysis was performed with Aminode (http://www.aminode.org/). Figures were assembled with Adobe Illustrator (2026).

### Statistics and reproducibility

All plots and statistical analyses were generated in RStudio 2024.04.1+748 (2024.04.1+748) with R version 4.4.0 (2024-04-24). Boxplots were generated in ggplot2, with the following parameters: lower whisker = smallest observation greater than or equal to lower hinge - 1.5 * IQR; lower = lower hinge, 25% quantile; notchlower = lower edge of notch = median - 1.58 * IQR/sqrt(n); middle = median, 50% quantile; notchupper = upper edge of notch = median + 1.58 * IQR/sqrt(n); upper = upper hinge, 75% quantile; ymax = upper whisker = largest observation less than or equal to upper hinge + 1.5 * IQR. Bar plots represent the mean ± s.e.m. with overlaid data points representing independent experiments or replicates. Results were considered significant at a FDR or *p*-value below 0.05. Unless otherwise indicated, statistical tests were two-sided. Immunofluorescence was independently repeated at least 4 times with similar results. The immunofluorescence and FRAP experiments shown in this paper were independently performed and quantified twice. Western blots and Co-immunoprecipitation experiments were independently replicated at least twice with similar results. Semi-quantitative RT–PCRs were performed once. Genotyping PCRs for positive CRISPR clones were repeated twice, followed by validation with complementary experimental approaches (qPCR and Western blot). Additional information and test results of statistical analysis are provided in the figure legends and Supplementary Data.

## Data availability

The RNA-seq data has been deposited at GEO under the accession GSE300638. The DESeq2 and rMATS results tables have been deposited ^18^. The mass spectrometry proteomics data have been deposited to the ProteomeXchange Consortium via the PRIDE ^91^ partner repository with the dataset identifier PXD065522 and PXD065532. The metabolomics data have been deposited to MetaboLights^92^ repository with the study identifier MTBLS13877.

## Acknowledgements

We thank Oliver Mühlemann and Alex Hofer (University of Bern) for sharing SMG inhibitors and for their expertise in nonsense-mediated mRNA decay and Maria Hondele for critical reading of the manuscript draft. We also thank the IMB Genomics Core Facility for assistance with library preparation and sequencing, Martin Möckel and the IMB Protein Production Core Facility for providing enzymes used in this work, Marton Gelleri together with the IMB Microscopy Core Facility for support with microscopy and FRAP experiments, Jasmin Cartano for proteomics sample processing and the IMB Flow Cytometry Core Facility for support. In addition, we thank the Imaging Core Facility (IMCF) and the FACS Core Facility at the Biozentrum, University of Basel, for technical assistance. CIKV acknowledges funding by the Deutsche Forschungsgemeinschaft (DFG, German Research Foundation) - Individual Grant Project no. 513744403, Scientific Network Grant Project no. 531902894, GRK2526 “Genevo” - Project no. 407023052”, GRK2859 (“4R”) - Project no. 491145305, Forschungsinitiative Rheinland-Pfalz (ReALity), the EMBO Young Investigator Program (5795), institutional funding from the Institute of Molecular Biology and funds from the Kanton Basel-Stadt and Basel-Land provided to the Biozentrum of the University Basel. J.H.G.F.G. was part of the ‘Science of Healthy Ageing Research Programme’ (SHARP) initiative funded by Rhineland-Palatinate’s Ministry of Science, Education and Culture. PR is funded by the Biozentrum PhD Fellowships Program. MFB received financial support from the intramural High Potentials Grant program of the University Medical Center Mainz, Forschungsinitiative Rheinland-Pfalz (ReALity) and Stiftungen zugunsten der Medizinischen Fakultät der LMU Klinikum (26069). Instruments in the IMB core facilities were supported by funds from the DFG: Laser Scanning Confocal (Leica Stellaris 8 Falcon, funded by the DFG - Project #497669232), Orbitrap Astral system (funded by the DFG - Project #524805621) and BD LSRFortessa SOPR is funded by the DFG - Project #210253511.

## Author Contributions

The project was conceptualized by CIKV. AVR performed the mouse experiments and isolated the tissues. PR performed cryosectioning and immunostainings/microscopy of mouse tissues. AG conducted microscopy and FRAP experiments and analyzed the data, with FRAP data analysis supported by FK. JHGFG provided genotyping PCR, Sanger Sequencing and one Western blot for *SNRPB* knock-out in cell clones pre-validated by FPH. JHGFG performed the long-term cell growth assay by flow cytometry and provided one replicate of Arg depletion Western blot experiment. JHGFG performed flow cytometry acquisition and analysis of the MitoTracker staining experiment. AE and JHGFG assisted with various experiments. FR and CIKV analyzed the RNA-seq data and performed all bioinformatic analysis, except for ribosome profiling, which was processed by AS. JXC conducted mass spectrometry and analyzed the data, with visualization performed by FPH. AA, SN and TS performed the metabolomics experiments under supervision of TS. FPH conducted all other experiments. SH shared expertise and supervised AVR. MFB designed, conducted and interpreted various experiments, was responsible for supervision and provided funding. CIKV performed bioinformatic analyses, interpreted the data, secured funding, supervised and coordinated the experiments. CIKV wrote the manuscript with support from FPH. All authors edited the manuscript.

## Ethics declarations

Mice were kept according to the protocols approved by Institutional Animal Care and Use Committee, Institutional Ethics Committee and the Preclinical Core Facility at Institute of Science and Technology of Austria (ISTA). Experimental protocols were performed under the approval of the Austrian Federal Ministry of Science and Research in accordance with the Austrian and European Union animal laws (license number: BMWFW-66.018/0015-WF/V/3b/2017; GZ: 2020-0.579.989; GZ: 2024-0.698.056). All mice used showed pathogen free status according to the Federation of European Laboratory Animal Science Associations (FELASA) recommendations.

