## Extended Data Figures for "The splicing paralogues *SNRPB* and *SNRPN* control differential metabolic states"

### Extended Data Figures and Legends

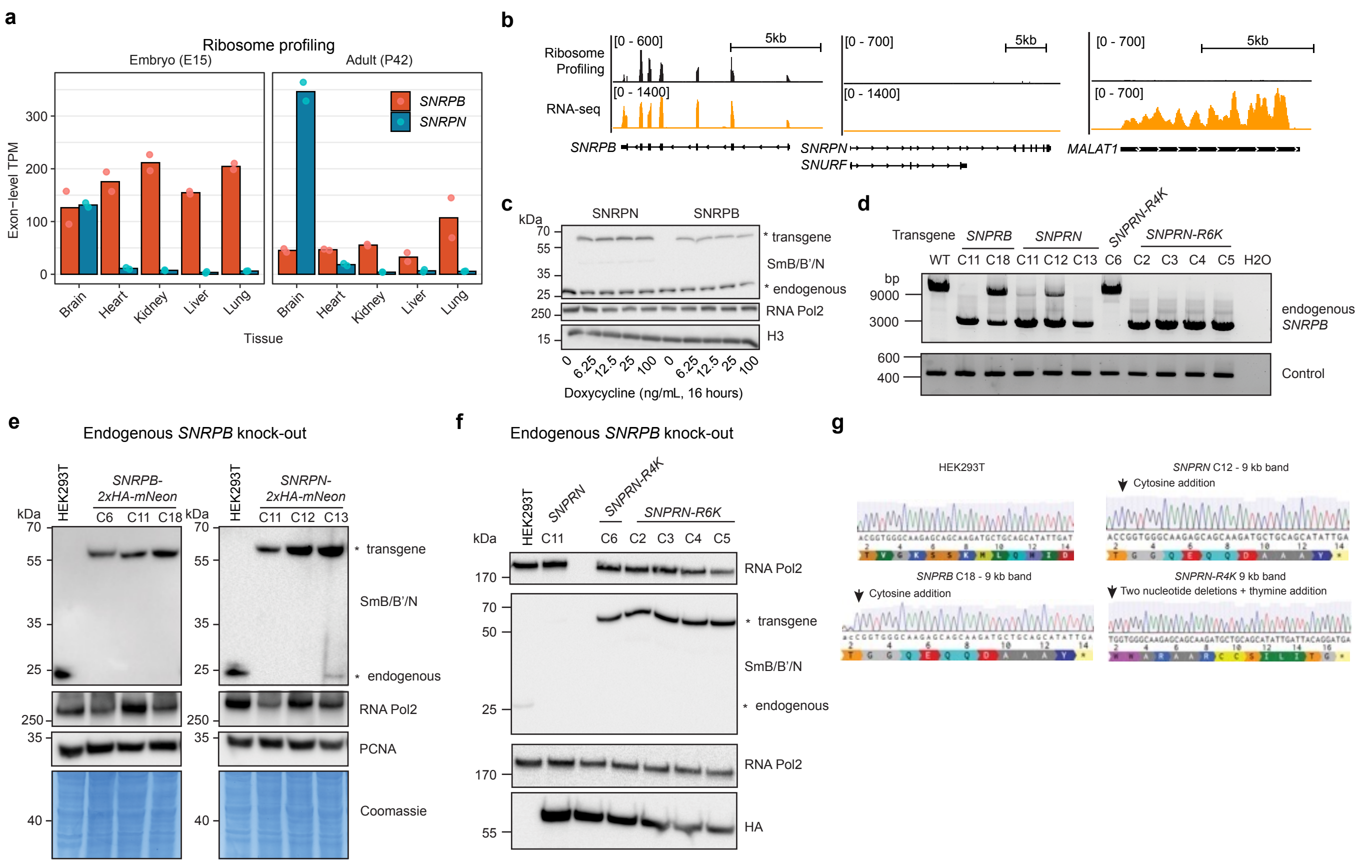
a
Ribosome profiling
Embryo (E15)
Adult (P42)
SNRPB
SNRPN
Exon−level TPM
0
100
200
300
Brain
Heart
Kidney
Liver
Lung
Tissue
b
Ribosome Profiling
RNA-seq
[0 - 600]
[0 - 1400]
[0 - 700]
[0 - 1400]
[0 - 700]
[0 - 700]
5kb
SNRPB
SNRPN
SNURF
MALAT1
c
kDa
SNRPN
SNRPB
* transgene
SmB/B'/N
* endogenous
RNA Pol2
H3
0
6.25
12.5
25
100
Doxycycline (ng/mL, 16 hours)
d
Transgene
SNPRB
SNPRN
SNPRN-R4K
SNPRN-R6K
WT
C11
C18
C11
C12
C13
C6
C2
C3
C4
C5
H2O
bp
9000
3000
600
400
endogenous SNRPB
Control
e
Endogenous SNRPB knock-out
HEK293T
SNRPB-2xHA-mNeon
SNRPN-2xHA-mNeon
C6
C11
C18
C11
C12
C13
kDa
* transgene
SmB/B'/N
* endogenous
RNA Pol2
PCNA
Coomassie
f
Endogenous SNRPB knock-out
HEK293T
SNPRN
SNPRN-R4K
SNPRN-R6K
C11
C6
C2
C3
C4
C5
kDa
RNA Pol2
* transgene
SmB/B'/N
* endogenous
RNA Pol2
HA
g
HEK293T
SNPRN C12 - 9 kb band
Cytosine addition
SNPRB C18 - 9 kb band
Cytosine addition
SNPRN-R4K 9 kb band
Two nucleotide deletions + thymine addition

ED Fig. 1

**Extended Data Fig. 1 - Characterization of transgenic lines with endogenous *SNRPB* knock-out.**

(a) Normalized exon-level transcript per million (TPM) from published [80] ribosome profiling data showing *SNRPB* and *SNRPN* translation across tissues in mouse E15 embryos and P42 adults.

(b) Ribosome profiling and RNA-seq coverage from [79] showing endogenous expression of *SNRPB*, but not *SNRPN*, in HEK293T cells. Tracks show ribosome occupancy (black) and transcript abundance (orange) across *SNRPB*, *SNRPN/SNURF*, and *MALAT1* loci. *SNRPB* displays transcript expression and ribosome occupancy, whereas *SNRPN* shows no ribosome engagement and minimal transcription. *MALAT1* is shown as a positive control long non-coding RNA that is not translated.

(c) Cropped immunoblots showing expression of HA-mNeon-tagged SNRPB or SNRPN transgenes under varying Doxycycline concentrations (endogenous SNRPB is present). Transgene-derived and endogenous SNRPB proteins are distinguished by molecular weight. RNA Pol2 and histone H3 serve as loading controls.

(d) Agarose gel showing PCR products across the endogenous *SNRPB* gene in parental HEK as well as different transgenic *SNRPB* knock-out cells (also see Sanger Sequencing results in (**g**)). The experiment was conducted once, results were confirmed with complementary methodology.

(e) Cropped immunoblots showing knock-out of endogenous *SNRPB* compared to parental HEK293T cell line. Transgene-derived and endogenous SNRPB proteins are distinguished by molecular weight. RNA Pol2, PCNA and Coomassie serve as loading controls. The results were validated two independent times.

(f) as in (e) but for SNRPN Arginine-to-Lysine mutant transgenes.

(g) Sanger Sequencing results for PCR products shown in (**d**), which confirm loss-of-function mutations in endogenous *SNRPB* gene.

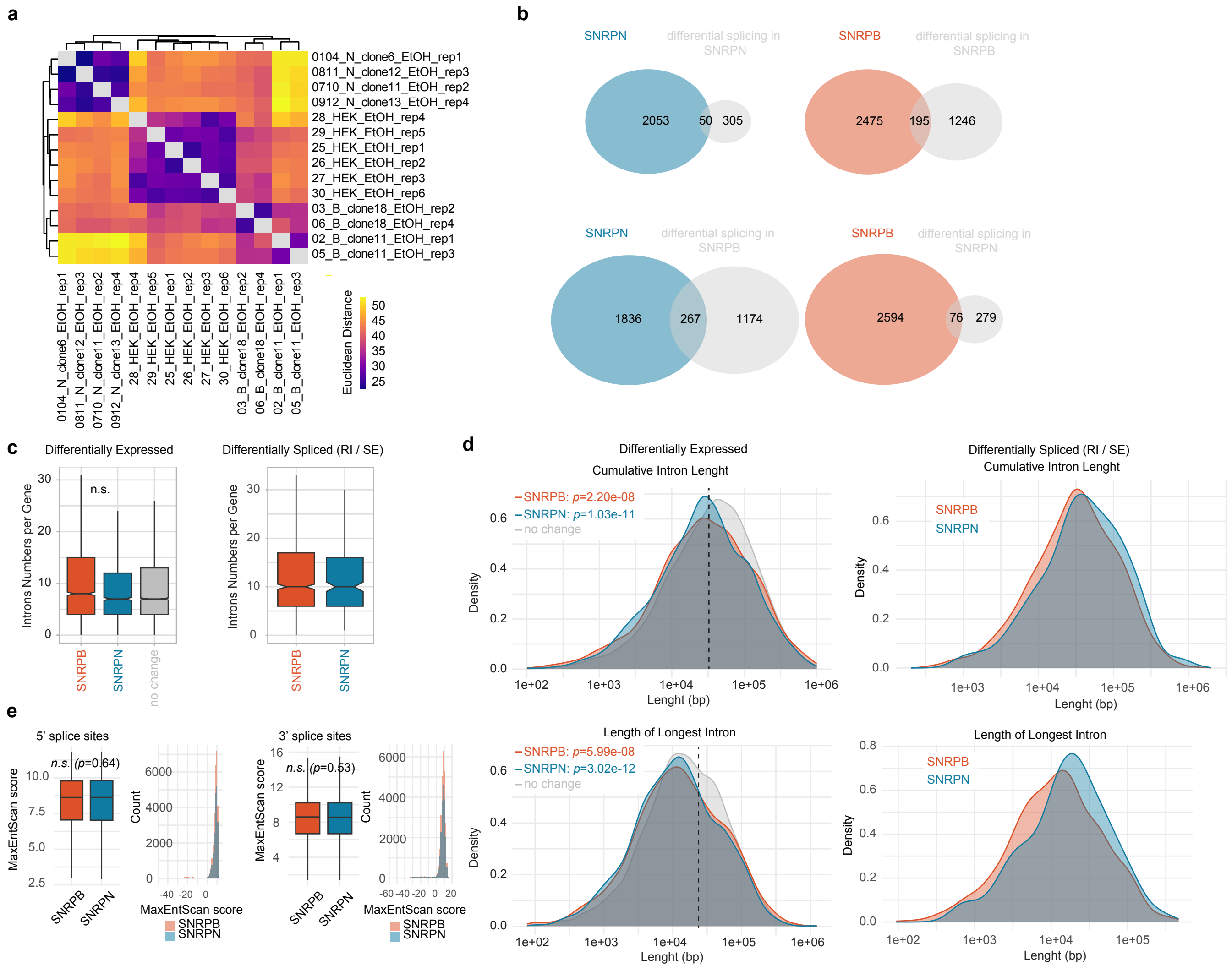
a
0104_N_clone6_EtOH_rep1
0811_N_clone12_EtOH_rep3
0710_N_clone11_EtOH_rep2
0912_N_clone13_EtOH_rep4
28_HEK_EtOH_rep4
29_HEK_EtOH_rep5
25_HEK_EtOH_rep1
26_HEK_EtOH_rep2
27_HEK_EtOH_rep3
30_HEK_EtOH_rep6
03_B_clone18_EtOH_rep2
06_B_clone18_EtOH_rep4
02_B_clone11_EtOH_rep1
05_B_clone11_EtOH_rep3
Euclidean Distance
b
SNRPN
differential splicing in SNRPN
2053
50
305
SNRPB
differential splicing in SNRPB
2475
195
1246
SNRPN
differential splicing in SNRPB
1836
267
1174
SNRPB
differential splicing in SNRPN
2594
76
279
c
Differentially Expressed
n.s.
Introns Numbers per Gene
SNRPB
SNRPN
no change
Differentially Spliced (RI / SE)
d
Differentially Expressed
Cumulative Intron Lenght
SNRPB: p=2.20e-08
SNRPN: p=1.03e-11
no change
Density
Lenght (bp)
Differentially Spliced (RI / SE)
Cumulative Intron Lenght
Length of Longest Intron
SNRPB: p=5.99e-08
SNRPN: p=3.02e-12
e
5' splice sites
n.s. (p=0.64)
MaxEntScan score
Count
3' splice sites
n.s. (p=0.53)

ED Fig. 2

**Extended Data Fig. 2 - Comparison of *SNRPB*- with *SNRPN*-expressing cells by RNA-seq.**

(a) Sample-to-sample distance matrix based on regularized log-transformed RNA-seq data from parental HEK cells, *SNRPB*-cells and SNRPN-cells (endogenous *SNRPB* knock-out).
(b) Venn diagram showing the overlap between DE genes (higher in SNRPN-cells: blue circles; higher in *SNRPB*-cells: red circles) and genes with differential splicing events (grey circles).
(c) Boxplots showing (left:) the Intron Numbers in DE genes (higher in *SNRPB*-cells: blue circles; higher in *SNRPB*-cells: red circles) compared to a control group (genes subsampled from non-DE genes with similar expression levels than *SNRPB* and *SNRPN*-DE). (right:) same for differentially spliced genes. Comparisons with unpaired Wilcoxon rank-sum tests did not reveal any significant *p*-values ($p<0.05$).
(d) Density plots comparing cumulative intron lengths and length of longest introns among genes that are DE (left) or differentially spliced (right) in *SNRPB*- and *SNRPN*- cells. Control group labelled as “no change” as in (c).
(e) Boxplots and histograms showing 5’ and 3’ splice site strengths of *SNRPB/N*-DE genes obtained by MaxEntScan [20]. Comparisons with unpaired Wilcoxon rank-sum tests did not reveal any significant *p*-values ($p<0.05$).

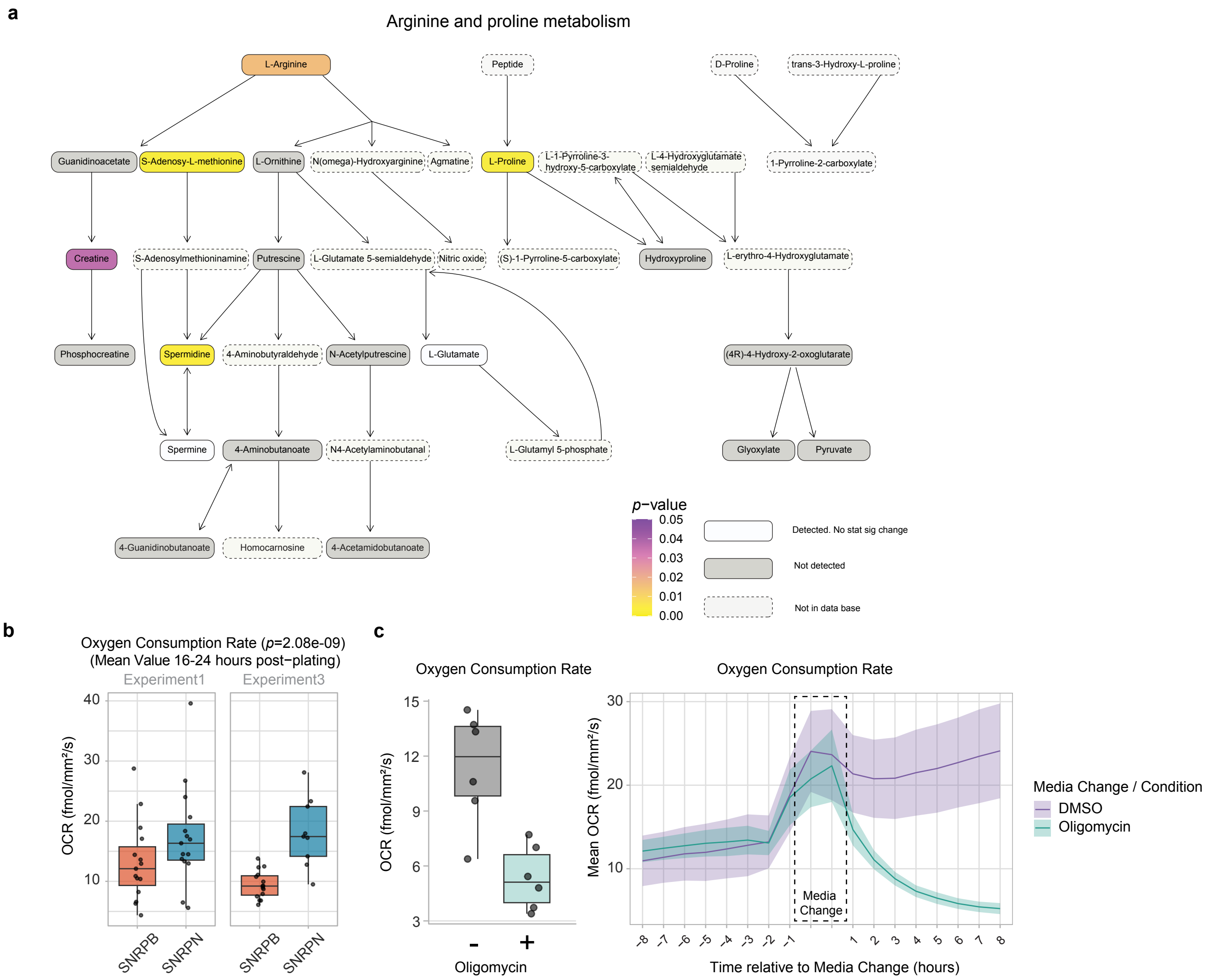
a
Arginine and proline metabolism
L-Arginine
Peptide
D-Proline
trans-3-Hydroxy-L-proline
Guanidinoacetate
S-Adenosy-L-methionine
L-Ornithine
N(omega)-Hydroxyarginine
Agmatine
L-Proline
L-1-Pyrroline-3-hydroxy-5-carboxylate
L-4-Hydroxyglutamate semialdehyde
1-Pyrroline-2-carboxylate
Creatine
S-Adenosylmethioninamine
Putrescine
L-Glutamate 5-semialdehyde
Nitric oxide
(S)-1-Pyrroline-5-carboxylate
Hydroxyproline
L-erythro-4-Hydroxyglutamate
Phosphocreatine
Spermidine
4-Aminobutyraldehyde
N-Acetylputrescine
L-Glutamate
(4R)-4-Hydroxy-2-oxoglutarate
Spermine
4-Aminobutanoate
N4-Acetylaminobutanal
L-Glutamyl 5-phosphate
Glyoxylate
Pyruvate
4-Guanidinobutanoate
Homocarnosine
4-Acetamidobutanoate
p−value
0.05
0.04
0.03
0.02
0.01
0.00
Detected. No stat sig change
Not detected
Not in data base
b
Oxygen Consumption Rate (p=2.08e-09)
(Mean Value 16-24 hours post−plating)
Experiment1
Experiment3
OCR (fmol/mm²/s)
40
30
20
10
SNRPB
SNRPN
SNRPB
SNRPN
c
Oxygen Consumption Rate
OCR (fmol/mm²/s)
15
12
9
6
3
-
+
Oligomycin
Oxygen Consumption Rate
Mean OCR (fmol/mm²/s)
30
20
10
Media Change
-8
-7
-6
-5
-4
-3
-2
-1
1
2
3
4
5
6
7
8
Time relative to Media Change (hours)
Media Change / Condition
DMSO
Oligomycin

ED Fig. 3

**Extended Data Fig. 3 - Further characterization of metabolomic changes in *SNRPB/N* cells.**

(a) KEGG pathway analysis of metabolomics data. The “Arginine and Proline Metabolism” pathway is colored to indicate metabolite changes, presence, or absence in metabolomics dataset.
(b) Box plots showing oxygen consumption rate (OCR) in *SNRPB*- and *SNRPN*-cells. Each data point represents a replicate, calculated as the mean of the recorded values between 16 and 24 hours after the start of the experiment. Three independent experiments were conducted on different days, two representative experiments are shown here, the other is in **Fig. 2e**. Statistical significance was assessed using a linear mixed-effects model across all three experiments.
(c) Box plots (left) and ribbon plot (right) showing OCR before and after treatment with oligomycin. Three independent experiments were conducted, with two replicates (data points) in each experiment. Each data point represents the mean OCR recorded over the 8 hours before (- oligomycin) or after (+ oligomycin) treatment, initiated by the media change. In the ribbon plot, the x-axis indicates time (in hourly bins relative to the treatment) and the y-axis shows the mean OCR for each hour. Shaded areas correspond to SEM, and lines represent mean values across all experiments and replicates. Before treatment, cells were grown in full media; during treatment, the media was replaced with DMSO (control) or oligomycin-containing medium.

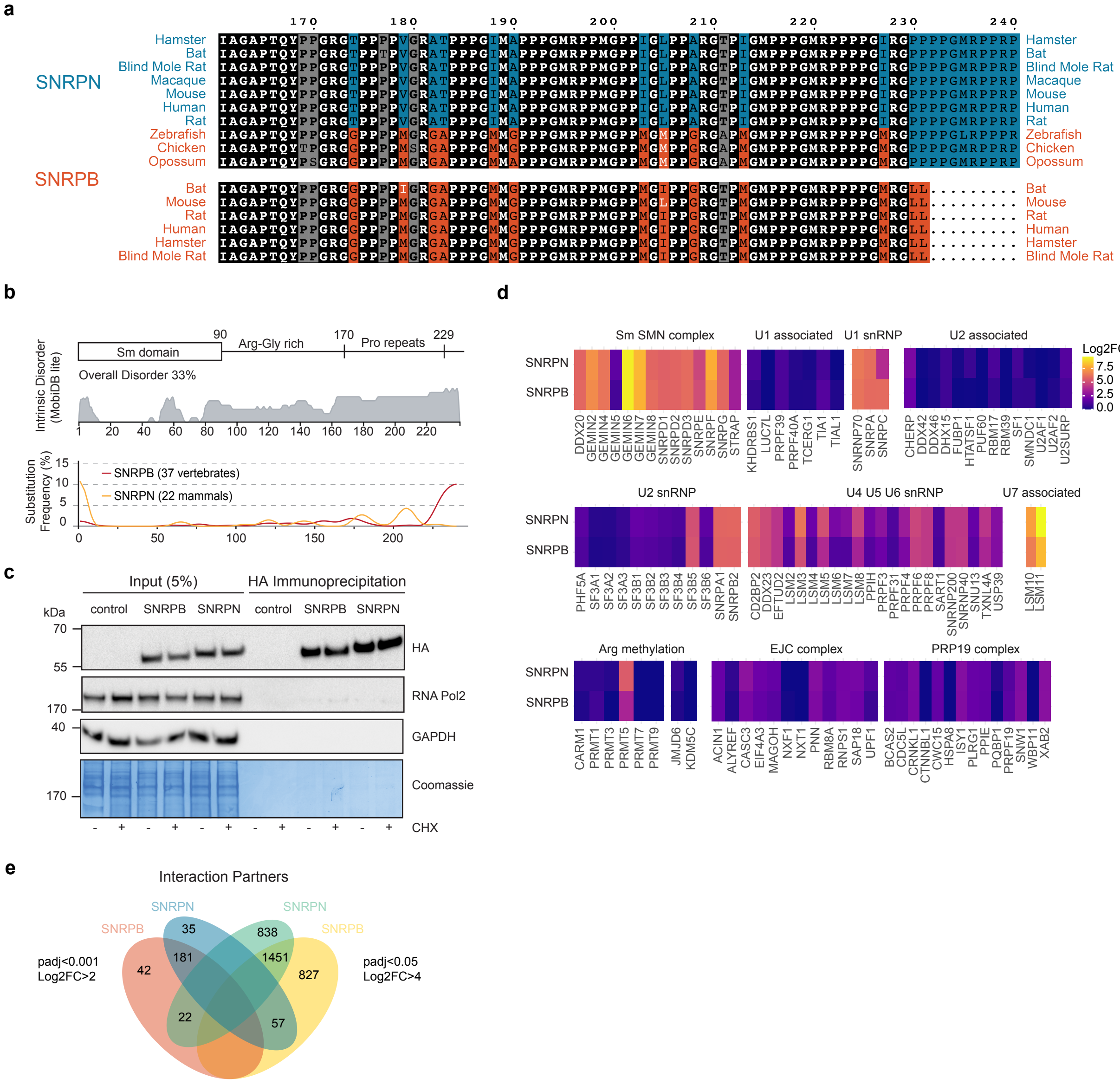
a
SNRPN
SNRPB
Hamster
Bat
Blind Mole Rat
Macaque
Mouse
Human
Rat
Zebrafish
Chicken
Opossum
b
Sm domain
90
Arg-Gly rich
170
Pro repeats
229
Overall Disorder 33%
Intrinsic Disorder (MobiDB lite)
Substitution Frequency (%)
SNRPB (37 vertebrates)
SNRPN (22 mammals)
c
Input (5%)
HA Immunoprecipitation
kDa
control
SNRPB
SNRPN
HA
RNA Pol2
GAPDH
Coomassie
CHX
d
Sm SMN complex
U1 associated
U1 snRNP
U2 associated
Log2FC
U2 snRNP
U4 U5 U6 snRNP
U7 associated
Arg methylation
EJC complex
PRP19 complex
e
Interaction Partners
padj<0.001
Log2FC>2
padj<0.05
Log2FC>4
35
838
42
181
1451
827
22
57

ED Fig. 4

**Extended Data Fig. 4 - Protein-level differences between SNRPB and SNRPN.**

(a) Protein sequence alignment of SNRPB and SNRPN in different species.

(b) Scheme showing the protein domain architecture and disorder content of SNRPB/N (top) and amino acid substitution frequency of the respective regions across vertebrates (SNRPB) or mammals (SNRPN).

(c) Cropped immunoblots of HA IPs in untagged control cells, and cells expressing HA-mNeon tagged SNRPB or SNRPN, with and without cycloheximide (CHX) treatment (100 µg/mL, 4 hours). HA confirms tagged protein expression and immunoprecipitation efficiency. RNA POL2, GAPDH, and Coomassie staining serve as loading, as well as negative IP controls.

(d) Heatmaps showing enrichment (Log2FC HA-IP versus control IP) of proteins in selected snRNP and RNA-processing complexes, grouped by function.

(e) Venn diagram showing overlap of interactors for SNRPB and SNRPN at two thresholds: high-confidence (padj<0.001, Log2FC>2) and low-confidence (padj<0.05, Log2FC>1).

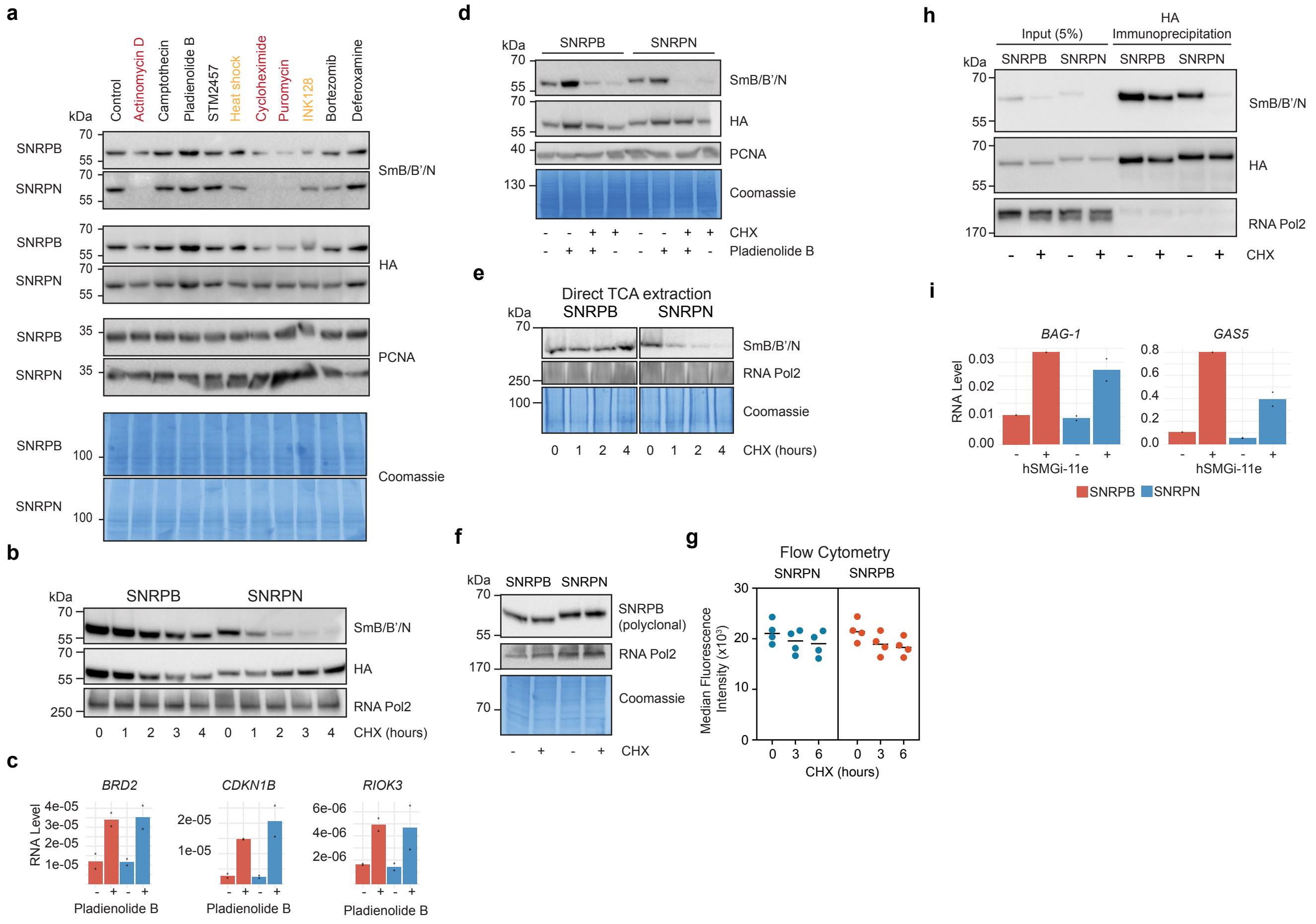

a
Control
Actinomycin D
Camptothecin
Pladienolide B
STM2457
Heat shock
Cycloheximide
Puromycin
INK128
Bortezomib
Deferoxamine
kDa
SNRPB
SNRPN
SmB/B'/N
HA
PCNA
Coomassie
b
SNRPB
SNRPN
SmB/B'/N
HA
RNA Pol2
CHX (hours)
c
BRD2
CDKN1B
RIOK3
RNA Level
Pladienolide B
d
SNRPB
SNRPN
SmB/B'/N
HA
PCNA
Coomassie
CHX
Pladienolide B
e
Direct TCA extraction
SNRPB
SNRPN
SmB/B'/N
RNA Pol2
Coomassie
CHX (hours)
f
SNRPB
SNRPN
SNRPB (polyclonal)
RNA Pol2
Coomassie
CHX
g
Flow Cytometry
SNRPN
SNRPB
Median Fluorescence Intensity (x10^3)
CHX (hours)
h
Input (5%)
HA Immunoprecipitation
SNRPB
SNRPN
SmB/B'/N
HA
RNA Pol2
CHX
i
BAG-1
GAS5
RNA Level
hSMGi-11e
SNRPB
SNRPN


ED Fig. 5

**Extended Data Fig. 5 - Characterization of 12F5 antibody detection under different conditions.**

(a) Cropped immunoblots of *SNRPB*- and *SNRPN*-cells treated with various compounds. PCNA and Coomassie serve as loading controls. HA-antibody shows overall protein stability differences. Monoclonal SmB/B'/N antibody (clone 12F5) shows detection differences for SNRPN under certain conditions. The experiment was performed twice with similar results. Treatments: Actinomycin D (transcription), heat shock, CHX (translation), puromycin (translation), Camptothecin (DNA Topoisomerase I), STM2457 (m6A-RNA methylation), Bortezomib (proteasome activity), Deferoxamine (iron homeostasis).
(b) Cropped immunoblots of *SNRPB*- and *SNRPN*-cells treated with CHX for the indicated hours. RNA POL2 serves as a loading control,  HA-antibody shows overall protein stability differences. The experiment was performed twice with similar results.
(c) RT-qPCR quantification validating splicing inhibition with Pladienoloide B (PlaB) (**Fig. 4f**). Intron RNA levels [93] were normalized to the geometric mean of 18S r*RNA N1* and *N5*. The bar indicates the mean, with overlaid data points representing biologically independent replicates.
(d) Cropped immunoblots of *SNRPB*- and *SNRPN*-cells co-treated with Pladienoloide B (PlaB, 16 hours, 100 nM) and CHX (4 hours, 100 µg/mL). RNA POL2 serves as a loading control, HA-antibody shows overall protein stability differences. The experiment was performed twice with similar results.
(e) Cropped immunoblots of *SNRPB*- and *SNRPN*-cells treated with CHX. After treatment, the cellular proteins were directly extracted with trichloroacetic acid. RNA POL2 and Coomassie serve as loading control The experiment was performed once.
(f) Cropped immunoblots of *SNRPB*- and *SNRPN*-cells treated with CHX. SNRPB/N was detected with a polyclonal antibody raised in *E. coli* and is insensitive to Arg-methylation. RNA POL2 and Coomassie serve as loading control The experiment was performed once.
(g) Barplot showing the median intensity of HA-mNeon-tagged SNRPB/N protein levels measured by flow cytometry after CHX treatment (Gatings in **Supplementary Data 2**).
(h) Cropped immunoblots of HA IPs of HA-mNeon tagged SNRPB or SNRPN, with and without cycloheximide (CHX) treatment (100 µg/mL, 4 hours). HA confirms tagged protein expression and immunoprecipitation efficiency. RNA POL2 serves as loading, as well as negative IP controls. The IP was conducted under high stringency conditions for PTM analysis.
(i) RT-qPCR quantification of selected NMD target gene [94] RNA levels validating SMGi inhibitor treatment. Expression levels were normalized to *RPLP0* expression. The bar indicates the mean, with overlaid data points representing biologically independent replicates.

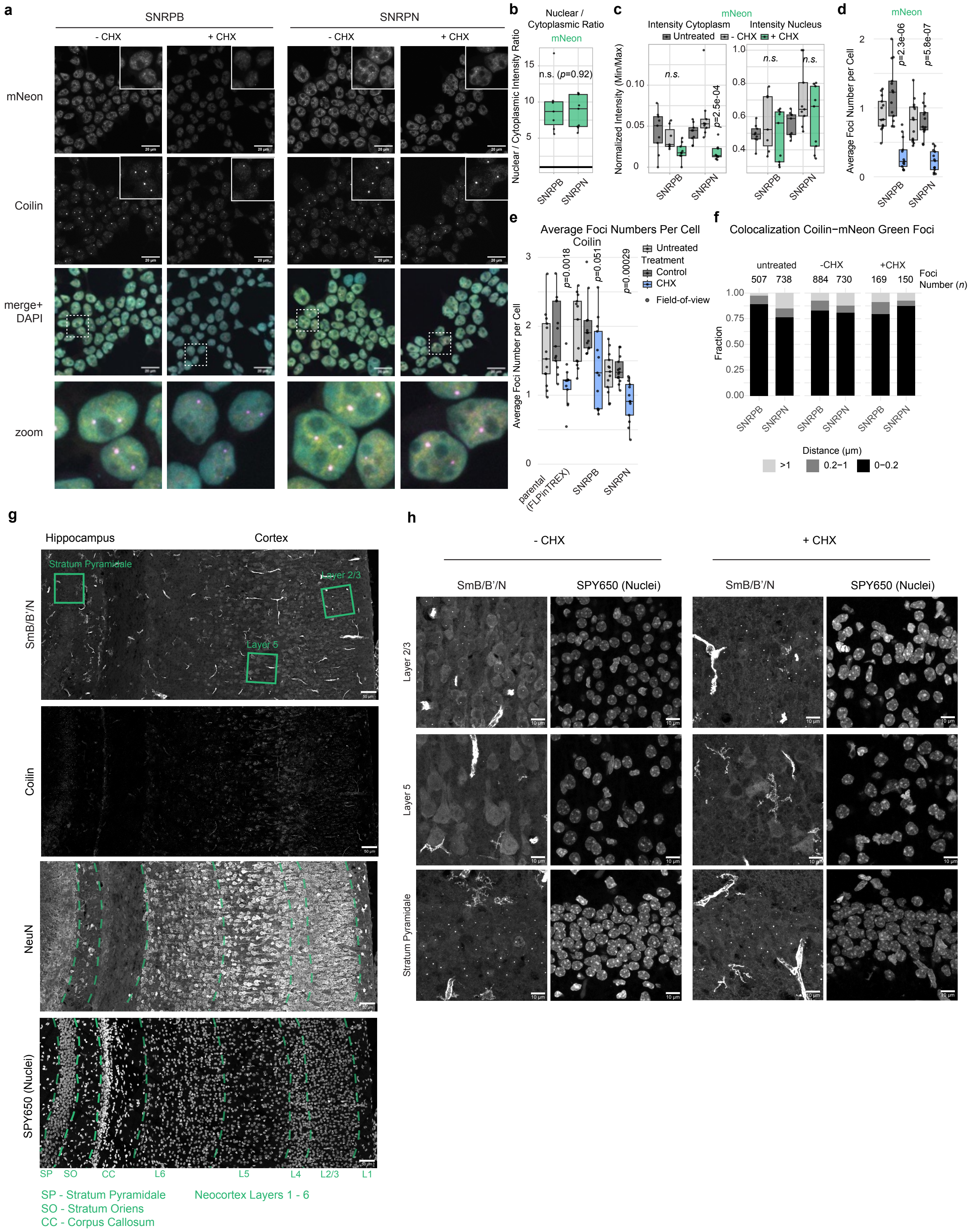
a
SNRPB
SNRPN
- CHX
+ CHX
mNeon
Coilin
merge+ DAPI
zoom
b
Nuclear / Cytoplasmic Ratio
mNeon
n.s. (p=0.92)
Nuclear / Cytoplasmic Intensity Ratio
c
mNeon
Intensity Cytoplasm
Intensity Nucleus
Untreated
- CHX
+ CHX
Normalized Intensity (Min/Max)
n.s.
p=2.5e-04
d
mNeon
p=2.3e-06
p=5.8e-07
Average Foci Number per Cell
e
Average Foci Numbers Per Cell
Coilin
Untreated
Treatment
Control
CHX
Field-of-view
p=0.0018
p=0.051
p=0.00029
parental (FLPinTREX)
SNRPB
SNRPN
f
Colocalization Coilin−mNeon Green Foci
untreated
-CHX
+CHX
Foci Number (n)
507
738
884
730
169
150
Fraction
Distance (µm)
>1
0.2−1
0−0.2
g
Hippocampus
Cortex
SmB/B'/N
Stratum Pyramidale
Layer 2/3
Layer 5
Coilin
NeuN
SPY650 (Nuclei)
SP
SO
CC
L6
L5
L4
L2/3
L1
SP - Stratum Pyramidale
SO - Stratum Oriens
CC - Corpus Callosum
Neocortex Layers 1 - 6
h
- CHX
+ CHX
SmB/B'/N
SPY650 (Nuclei)
Layer 2/3
Layer 5
Stratum Pyramidale

**Extended Data Fig. 6 - Further investigation of SNRPB/N behaviour in cells and mouse brain.**

(a) Representative fluorescence microscopy images of *SNRPB/N*-cells employing the mNeon-tag (yellow in merged panel), immunostained Coilin (magenta in merged panel) and DAPI (cyan in merged panel) in Ethanol (-CHX; Control) or CHX treatment (4 hours), scale bar = 20µm. The areas marked with a dashed square are magnified in the "zoom" panel. The experiment was conducted two independent times. The images shown are maximum projections of z-stacks.
(b)-(e) The total quantified cell numbers were: HEK293T untreated (*n*=578), HEK293T-CHX (*n*=541), HEK293T+CHX (*n*=359), SNRPB untreated (*n*=580), SNRPB-CHX (*n*=737), SNRPB+CHX (*n*=579), SNRPN untreated (*n*=918), SNRPN-CHX (*n*=876), SNRPN+CHX (*n*=692).
(b) Quantification of (**a**). The intensity of mNeonGreen (-CHX condition) was quantified in the nucleus and cytoplasm (for details see methods) and nuclear / cytoplasmic ratio was determined for each field of view. Each dot represents the result of one field of view and the boxplot shows the distribution across fields of view. The *p*-value was obtained with an unpaired Wilcoxon rank sum test.
(c) Quantification of (**a**) as described in (**b**) - for the mNeonGreen intensity in cytoplasm and nucleus. The adjusted p-values were obtained with a Pairwise Wilcoxon rank sum exact test with Benjamini-Hochberg correction.
(d)-(e) Boxplot showing the average foci numbers per cell in a given field of view. Foci were identified from the mNeonGreen (**d**) and Coilin (**e**) channels (see methods) and divided by the total number of cells (manually counted) in a given field of view. Each dot represents the result of one field of view and the boxplot shows the distribution across fields of view. The adjusted p-values were obtained with a Pairwise Wilcoxon rank sum exact test with Benjamini-Hochberg correction and shown for the Ethanol (-CHX) versus +CHX condition. None of the comparisons between Ethanol (-CHX; control) and untreated were significant (padj<0.05). The untreated control experiment was stained and acquired simultaneously and is shown in **Fig. 1b**.
(f) Quantification of (**a**). Stacked Barplot showing the distance of each mNeonGreen focus to the nearest coilin focus as a fraction in 3 groups (>1 µm,0.2-1 µm and 0-0.2 µm). The number of identified mNeonGreen foci is given in the figure.
(g) Representative confocal microscopy pictures of a mouse brain section stained with 12F5 (SmB/B'/N), Coilin, NeuN or SPY650, the latter of which stains nuclei akin to DAPI. The different areas are delineated with green squares / dashed lines. Scale bar 50 µm.
(h) Representative confocal microscopy pictures mouse brain tissue in control and CHX-treatment condition and stained with SmB/B'/N (12F5) and SPY650, the latter of which stains nuclei akin to DAPI. Areas from the mouse cortex (Layers2/3 and Layer5) and hippocampus are shown. The pictures represent a maximum intensity projection of a z-stackScale bar 10 µm. Quantification in **Fig. 5e**.

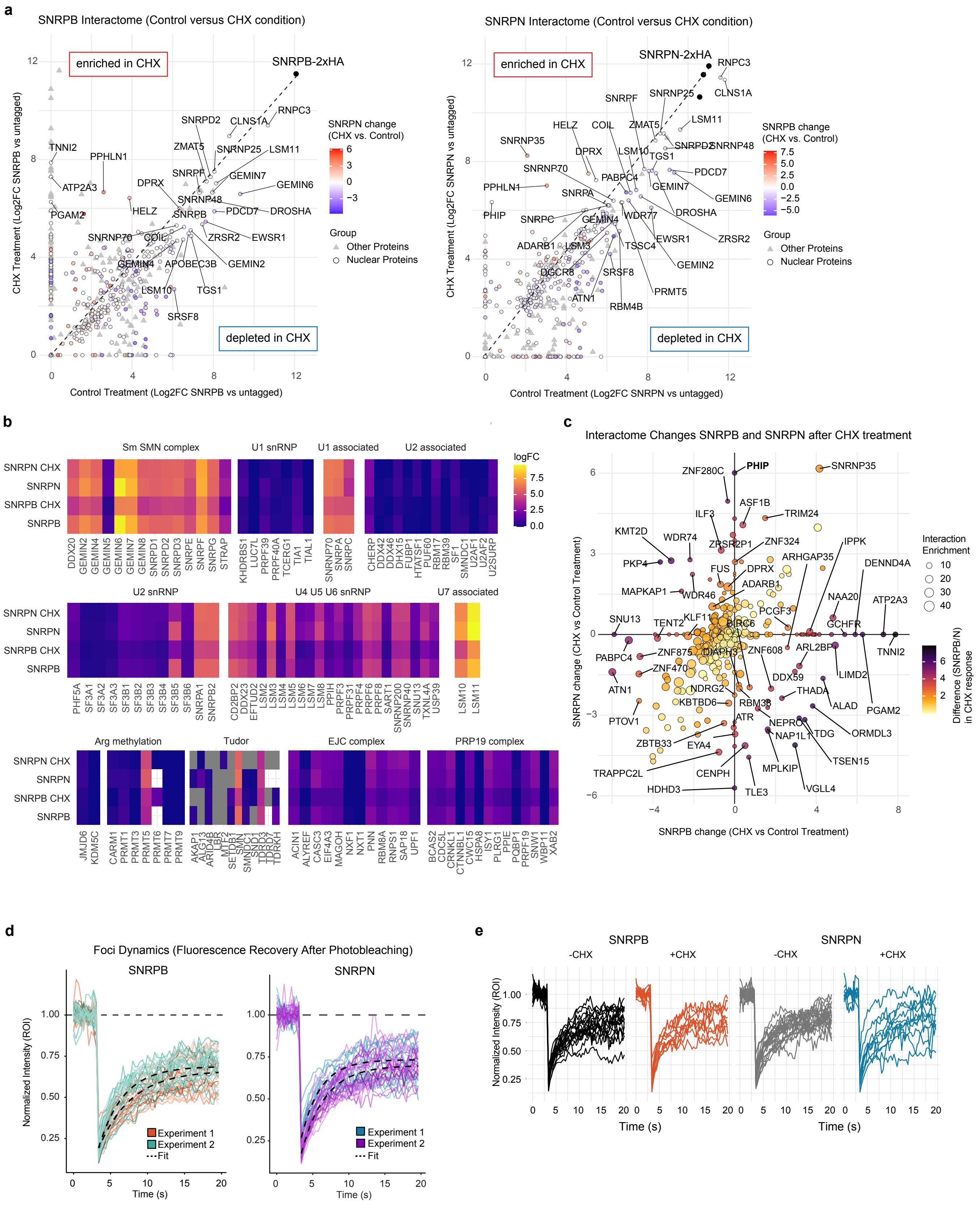
a
SNRPB Interactome (Control versus CHX condition)
enriched in CHX
depleted in CHX
SNRPB-2xHA
CHX Treatment (Log2FC SNRPB vs untagged)
Control Treatment (Log2FC SNRPB vs untagged)
SNRPN change (CHX vs. Control)
Group
Other Proteins
Nuclear Proteins
TNNI2
PPHLN1
ATP2A3
PGAM2
HELZ
DPRX
SNRPD2
CLNS1A
RNPC3
ZMAT5
SNRNP25
LSM11
SNRPF
GEMIN7
GEMIN6
SNRNP48
PDCD7
DROSHA
SNRPB
ZRSR2
EWSR1
SNRNP70
COIL
GEMIN4
APOBEC3B
GEMIN2
LSM10
TGS1
SRSF8
SNRPN Interactome (Control versus CHX condition)
SNRPN-2xHA
CHX Treatment (Log2FC SNRPN vs untagged)
Control Treatment (Log2FC SNRPN vs untagged)
SNRPB change (CHX vs. Control)
SNRPF
SNRNP25
CLNS1A
HELZ
COIL
ZMAT5
LSM11
SNRNP35
DPRX
LSM10
SNRPD2
SNRNP48
TGS1
SNRNP70
PABPC4
PDCD7
PPHLN1
SNRPA
GEMIN7
GEMIN6
PHIP
SNRPC
GEMIN4
WDR77
DROSHA
ADARB1
LSM3
TSSC4
EWSR1
ZRSR2
DGCR8
SRSF8
GEMIN2
ATN1
PRMT5
RBM4B
b
Sm SMN complex
U1 snRNP
U1 associated
U2 associated
logFC
SNRPN CHX
SNRPN
SNRPB CHX
SNRPB
DDX20
GEMIN2
GEMIN4
GEMIN5
GEMIN6
GEMIN7
GEMIN8
SNRPD1
SNRPD2
SNRPD3
SNRPE
SNRPF
SNRPG
STRAP
KHDRBS1
LUC7L
PRPF39
PRPF40A
TCERG1
TIA1
TIAL1
SNRNP70
SNRPA
SNRPC
CHERP
DDX42
DDX46
DHX15
FUBP1
HTATSF1
PUF60
RBM17
RBM39
SF1
SMNDC1
U2AF1
U2AF2
U2SURP
U2 snRNP
U4 U5 U6 snRNP
U7 associated
PHF5A
SF3A1
SF3A2
SF3A3
SF3B1
SF3B2
SF3B3
SF3B4
SF3B5
SF3B6
SNRPA1
SNRPB2
CD2BP2
DDX23
EFTUD2
LSM2
LSM3
LSM4
LSM5
LSM6
LSM7
LSM8
PPIH
PRPF3
PRPF31
PRPF4
PRPF6
PRPF8
SART1
SNRNP200
SNRNP40
SNU13
TXNL4A
USP39
LSM10
LSM11
Arg methylation
Tudor
EJC complex
PRP19 complex
JMJD6
KDM5C
CARM1
PRMT1
PRMT3
PRMT5
PRMT6
PRMT7
PRMT9
AKAP1
ALG13
ARID4B
LBR
MTF2
SETDB1
SMN1
SMNDC1
SND1
TDRD3
TDRD7
TDRKH
ACIN1
ALYREF
CASC3
EIF4A3
MAGOH
NXF1
NXT1
PNN
RBM8A
RNPS1
SAP18
UPF1
BCAS2
CDC5L
CRNKL1
CTNNBL1
CWC15
HSPA8
ISY1
PLRG1
PPIE
PQBP1
PRPF19
SNW1
WBP11
XAB2
c
Interactome Changes SNRPB and SNRPN after CHX treatment
SNRPN change (CHX vs Control Treatment)
SNRPB change (CHX vs Control Treatment)
Interaction Enrichment
Difference (SNRPB/N) in CHX response
ZNF280C
PHIP
SNRNP35
ASF1B
ILF3
TRIM24
KMT2D
WDR74
ZRSR2P1
ZNF324
IPPK
PKP4
FUS
ARHGAP35
DENND4A
DPRX
ADARB1
MAPKAP1
WDR46
NAA20
ATP2A3
PCGF3
SNU13
TENT2
KLF11
BIRC6
GCHFR
TNNI2
PABPC4
ZNF875
DIAPH3
ZNF608
ARL2BP
ZNF470
DDX59
LIMD2
ATN1
NDRG2
THADA
KBTBD6
RBM38
ALAD
PGAM2
PTOV1
ATR
NEPRO
NAP1L1
TDG
ORMDL3
ZBTB33
EYA4
TSEN15
TRAPPC2L
CENPH
MPLKIP
HDHD3
TLE3
VGLL4
d
Foci Dynamics (Fluorescence Recovery After Photobleaching)
SNRPB
SNRPN
Normalized Intensity (ROI)
Time (s)
Experiment 1
Experiment 2
Fit
e
SNRPB
SNRPN
-CHX
+CHX
-CHX
+CHX
Normalized Intensity (ROI)
Time (s)

ED Fig. 7

**Extended Data Fig. 7 - *SNRPB/N* interactome and dynamics after CHX treatment.**

(a) Dot plots of HA IP-mass spectrometry experiment comparing HA-mNeon tagged SNRPB (left) or SNRPN (right) in control versus CHX-treatment conditions. The dashed diagonal indicates equal enrichment between the two interactomes and selected proteins with equal enrichment and changing interactions are labeled. *n*=4 replicates for each SNPRB/N were conducted, Log2FC was obtained with limma [66].

(b) Heatmaps showing enrichment (Log2FC HA-IP versus control IP, with and without CHX treatment) of proteins in selected snRNP and RNA-processing complexes, grouped by function. Also see **Extended Data Fig. 4d.**

(c) Dot plots visualizing the correlation of interactome changes in SNRPB (CHX versus Control) and SNRPN (CHX versus Control). Interaction partners that change equally for both paralogues reside along the diagonal and are colored in white-to-yellow (see color map). Interaction partners that behave differently are outside the diagonal and have red-purple-black colored shades. The dot size reflects the absolute Log2FC of the interactors versus the untagged control (low to high enrichment).

(d) Normalized intensity curves from FRAP experiments, where each line represents the recorded data of one bleached focus in the nucleus. The lines show the recovery of nuclear foci in *SNRPB*- or *SNRPN*-cells, respectively, with overall fitted curves for each set of experimental curves shown in dashed lines (details on normalization and data fitting in methods). The experiment was conducted two times independently, cells were untreated (neither Ethanol nor CHX added).

(e) FRAP as in (**d**) Normalized intensity curves, where each line represents the recorded data of one bleached focus in the nucleus. See also **Fig. 5f** and **Fig. 5g.**

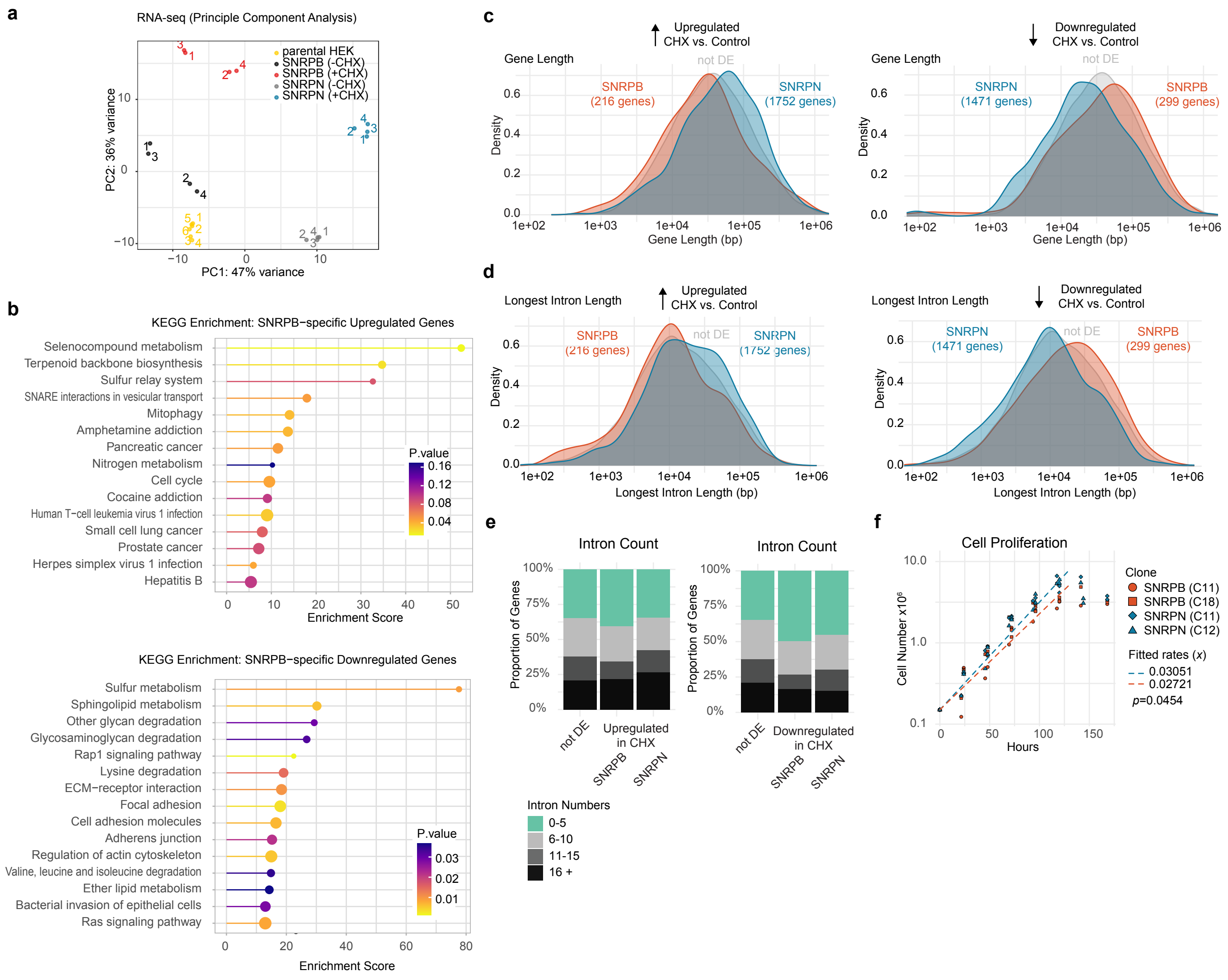
a
RNA-seq (Principle Component Analysis)
parental HEK
SNRPB (-CHX)
SNRPB (+CHX)
SNRPN (-CHX)
SNRPN (+CHX)
PC2: 36% variance
PC1: 47% variance
b
KEGG Enrichment: SNRPB−specific Upregulated Genes
Selenocompound metabolism
Terpenoid backbone biosynthesis
Sulfur relay system
SNARE interactions in vesicular transport
Mitophagy
Amphetamine addiction
Pancreatic cancer
Nitrogen metabolism
Cell cycle
Cocaine addiction
Human T−cell leukemia virus 1 infection
Small cell lung cancer
Prostate cancer
Herpes simplex virus 1 infection
Hepatitis B
P.value
Enrichment Score
KEGG Enrichment: SNRPB−specific Downregulated Genes
Sulfur metabolism
Sphingolipid metabolism
Other glycan degradation
Glycosaminoglycan degradation
Rap1 signaling pathway
Lysine degradation
ECM−receptor interaction
Focal adhesion
Cell adhesion molecules
Adherens junction
Regulation of actin cytoskeleton
Valine, leucine and isoleucine degradation
Ether lipid metabolism
Bacterial invasion of epithelial cells
Ras signaling pathway
P.value
Enrichment Score
c
Gene Length
Upregulated CHX vs. Control
SNRPB (216 genes)
not DE
SNRPN (1752 genes)
Density
Gene Length (bp)
Gene Length
Downregulated CHX vs. Control
SNRPN (1471 genes)
not DE
SNRPB (299 genes)
Density
Gene Length (bp)
d
Longest Intron Length
Upregulated CHX vs. Control
SNRPB (216 genes)
not DE
SNRPN (1752 genes)
Density
Longest Intron Length (bp)
Longest Intron Length
Downregulated CHX vs. Control
SNRPN (1471 genes)
not DE
SNRPB (299 genes)
Density
Longest Intron Length (bp)
e
Intron Count
Proportion of Genes
not DE
Upregulated in CHX
SNRPB
SNRPN
Intron Count
Proportion of Genes
not DE
Downregulated in CHX
SNRPB
SNRPN
Intron Numbers
0-5
6-10
11-15
16 +
f
Cell Proliferation
Cell Number x10⁶
Hours
Clone
SNRPB (C11)
SNRPB (C18)
SNRPN (C11)
SNRPN (C12)
Fitted rates (x)
0.03051
0.02721
p=0.0454

ED Fig. 8

**Extended Data Fig. 8 - RNA-seq of *SNRPB/N*-cells upon CHX treatment.**

(a) Principal component (PC) analysis plot representing the PC1 and PC2 contribution to the variance in gene expression between samples in RNA-seq.
(b) KEGG pathway enrichment analysis of uniquely DE genes in *SNRPB*-cells after CHX treatment. Enrichment scores are plotted on the *x*-axis, with dot size corresponding to gene overlap and color representing adjusted *p*-values.
(c), (d) Density plots comparing gene lengths (**c**) and length of longest introns (**d**) among genes that are uniquely DE (see also **Fig. 6b**) in *SNRPB*- (red) or *SNRPN*- (blue) cells after CHX treatment. The control group ("not DE", grey) are genes without significant changes and >100 mean read counts.
(e) Barplot showing the intron numbers of the same gene groups as in (**c**).
(f) Cell proliferation curves for *SNRPB*- and *SNRPN*-cells. Cell numbers were measured over 150 hours, with each dot representing a biological replicate and clone (indicated by different symbols). In contrast to **Fig. 7f**, cells were not split; instead, replicate wells derived from the same starting culture were counted at successive time points after plating. Statistical significance was evaluated using a linear regression model (fits shown by the dashed line) on log-transformed cell numbers. As cells reached full confluency at the final two time points, no additional growth was detectable; therefore, these data points were excluded from the regression fit.
